# The priming effect of rewarding brain stimulation in rats depends on both the cost and strength of reward but survives blockade of D2-like dopamine receptors

**DOI:** 10.1101/2022.10.20.512941

**Authors:** Czarina Evangelista, Norhan Mehrez, Esthelle Ewusi Boisvert, Wayne G. Brake, Peter Shizgal

## Abstract

Receipt of an intense reward boosts motivation to work for more of that reward. This phenomenon is called the *priming effect of rewards*. Using a novel measurement method, we show that the priming effect of rewarding electrical brain stimulation depends on the cost, as well as on the strength, of the anticipated reward. Previous research on the priming effect of electrical brain stimulation utilized a runway paradigm in which running speed serves as the measure of motivation. In the present study, the measure of motivation was the vigor with which rats executed a two-lever response chain, in a standard operant-conditioning chamber, to earn rewarding electrical stimulation of the lateral hypothalamus. In a second experiment, we introduced a modification that entails self-administered priming stimulation and alternating blocks of primed and unprimed trials. Reliable, consistent priming effects of substantial magnitude were obtained in the modified paradigm, which is closely analogous to the runway paradigm. In a third experiment, the modified paradigm served to assess the dependence of the priming effect on dopamine D2-like receptors. The priming effect proved resilient to the effect of eticlopride, a selective D2-like receptor antagonist. These results are discussed within the framework of a new model of brain reward circuitry in which non-dopaminergic medial forebrain bundle fibers and dopamine axons provide parallel inputs to the final common paths for reward and incentive motivation.

## Introduction

Receipt of an intense reward boosts motivation to seek a subsequent reward. This phenomenon is called the priming effect of rewards. Despite the common terminology, this priming effect of rewards differs from the priming effect observed in the reinstatement model of drug relapse (de Wit & Stewart, 1981, 1983). In that model, drug seeking is learned and then extinguished. Non-contingent delivery of a reward (a “prime”), can re-establish the previously extinguished drug-seeking behavior. That phenomenon is also commonly called priming-induced reinstatement. In contrast, the priming effect of rewards discussed here invigorates a well-established behavior that has not undergone extinction (Evangelista et al., 2019; C. R. Gallistel, 1966, 1969; C. R. Gallistel et al., 1974).

The priming effect of rewarding electrical brain stimulation manifests two key characteristics of motivation: It both directs and energizes goal-seeking behavior. Rats seek out a primed reward in preference to a competing unprimed reward (Deutsch et al., 1964; Wasserman et al., 1982), and they run down a runway faster to obtain a reward after having received priming (Edmonds et al., 1974; Edmonds & Gallistel, 1974; C. R. Gallistel, 1966; C. R. Gallistel et al., 1974).

Previous studies document the dependence of the priming effect of rewarding electrical brain stimulation on the rat’s expectation of reward strength (Edmonds et al., 1974; Edmonds & Gallistel, 1974; C. R. Gallistel, 1966; C. R. Gallistel et al., 1974). In these studies, pre-trial stimulation is delivered by the experimenter or the rat in the start box of a runway or in a waiting box prior to transfer to the runway start box. When a barrier blocking exit from the start box is removed, the rat can run down the alley to the goal box, where a lever press triggers delivery of rewarding stimulation. With the strength and amount of pre-trial stimulation held constant, running speed depends on the strength (e.g., current or pulse frequency) of the stimulation available in the goal box (C. R. Gallistel et al., 1974). The stronger the reward the rat expects to earn in the goal box, the faster it runs.

In Experiment 1, we ask a question complementary to one posed by the studies showing the dependence of running speed on reward strength: How does the vigor of the rat’s reward-seeking behavior vary as a function of its expectation of reward *cost* – the effort and time entailed in meeting a fixed-ratio requirement? To pose this question, we adapted the runway paradigm for use in standard operant-conditioning chambers, substituting lever-pressing behavior for alley running as a measure of motivation. After receiving non-contingent stimulation in Experiment 1, rats pressed a setup lever once to trigger extension of a stimulation lever from the opposite wall of the chamber. To earn a reward, the rat had to complete a fixed-ratio response requirement on the stimulation lever. Using this design, we measured the dependence of the priming effect on the cost and strength of the response-contingent stimulation. We modified the paradigm in Experiment 2 to achieve greater parallelism with Gallistel’s runway paradigm, to reduce putative disruptive effects of non-contingently delivered priming stimulation, and to isolate the effect of priming from within- and across-session drifts in performance. In Experiment 3, the modified paradigm developed in Experiment 2 was employed to assess the modulation of the priming effect by eticlopride, a highly selective dopamine D2-like receptor (D2R) antagonist (Hall et al., 1985; Martelle & Nader, 2008). In all three experiments, the electrodes were aimed at the site most frequently used in previous work on the priming effect of rewarding brain stimulation: the medial forebrain bundle (MFB), at the level of the lateral hypothalamus (LH).

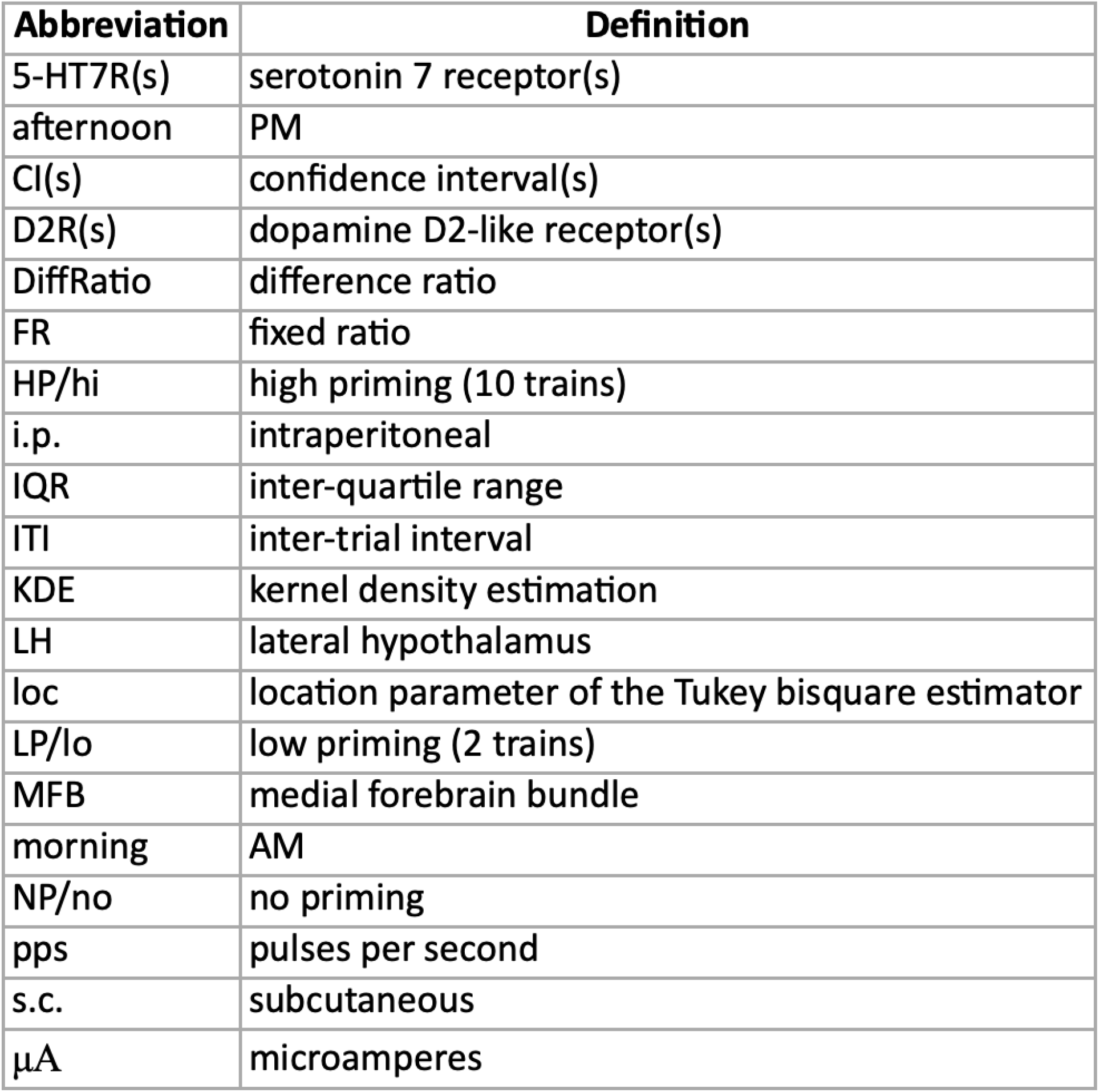

## Experiment 1

Reward seeking is typically a sigmoidal function of pulse frequency: Rats work most vigorously when pulse frequency is high, fail to respond once pulse frequency is low, and respond at intermediate levels when the pulse frequency is moderate (Miliaressis et al., 1986; Yeomans, 1975). An analogous relationship between response vigor and response-contingent stimulation strength holds in the runway paradigm (Edmonds et al., 1974; Edmonds & Gallistel, 1974; C. R. Gallistel, 1966; C. R. Gallistel et al., 1974). In the present study, we assessed whether a similar dependence is also seen in our novel paradigm for measuring the priming effect.

Reward seeking depends not only on stimulation strength but also on two types of reward cost. Opportunity cost is the work time required to obtain a reward (Breton et al., 2009), whereas effort cost is the physical exertion required to earn a reward (e.g., the effort entailed in meeting a fixed-ratio requirement). Reward seeking is highest when the opportunity cost is low and decreases as the cost grows (Hernandez et al., 2010, 2012; Trujillo-Pisanty et al., 2014).

Similarly, rats are less willing to work as the effort cost of reward increases (Aberman & Salamone, 1999; Salamone et al., 2001). Here, we determined whether the magnitude of the priming effect depends on reward costs.

### Experiment 1: Method

#### Subjects

Male Long-Evans rats (bred at Concordia University; n = 8) were pair-housed in Plexiglas^®^ cages (46 cm length x 26 cm width x 21 cm height) located in a vivarium with a reversed 12-hour light-dark cycle (lights off from 0800 to 2000 h). Throughout the study, rats had *ad libitum* access to food and water. A mix of Teklad corncob and Sani-Chips^®^ (Envigo, Madison, Wisconsin, USA) were used as bedding, and cages were enriched with shredded paper and a tunnel toy. After surgery, rats were housed individually for the remainder of the experiment. Behavioral tests were conducted during the dark phase of the diurnal cycle. The protocols used were in accordance with guidelines established by Concordia University’s Animal Research Ethics Committee’s Terms of Reference and the Canadian Council on Animal Care’s Guide to the Care and Use of Experimental Animals.

#### Electrode Implantation

At the time of surgery, rats were 3-4 months old and weighed at least 350 g. A ketamine-xylazine mixture (87 mg/kg of ketamine, 13 mg/kg xylazine), Bioniche, Belleville, ON, Canada; Bayer Healthcare, Toronto, ON, Canada) was administered intraperitoneally (i.p) to induce anesthesia. This was followed by a subcutaneous (s.c.) injection of atropine sulfate (0.05 mg/kg, Sandoz, Boucherville, QC, Canada) to reduce bronchial secretions, and penicillin (0.3 ml, s.c., Vetoquinol, Lavaltrie, QC, Canada) to prevent infections. Xylocaine jelly (AstraZeneca, Mississauga, ON, Canada) was applied to the external auditory meatus to diminish discomfort due to the stereotaxic ear bars. After placing the rat in the stereotaxic frame, a mixture of isoflurane (Pharmaceutical Partners of Canada Inc., Richmond Hill, ON, Canada) and oxygen was delivered through a snout mask to maintain anesthesia. Four to six burr holes were drilled into the skull, and stainless-steel screws were threaded. The free end of the current-return wire was wrapped around two skull screws (which served as the anode), and the opposite end was terminated in a gold-plated Amphenol connector. Monopolar stainless-steel electrodes were fashioned from insect pins (size: 000) insulated with Formvar enamel, leaving 0.5 mm of tip bare. The unsharpened end was soldered to a copper wire that was attached to a gold-plated Amphenol connector. Electrodes were aimed bilaterally at the LH level of the MFB; the target coordinates were: AP: -2.8 from bregma, ML: ±1.7, and DV: -8.8-9.0 from skull surface. The electrodes were secured to the skull with dental acrylic. The Amphenol connectors were inserted into a McIntyre miniature connector (Scientific Technology Centre, Carleton University, Ottawa, ON, Canada) that was attached to the skull and skull-screw anchors using dental acrylic. The rats were allowed at least one week to recover from the surgery before self-stimulation training commenced. Electrode-tip locations are shown in Figure 1.

**Figure 1.**
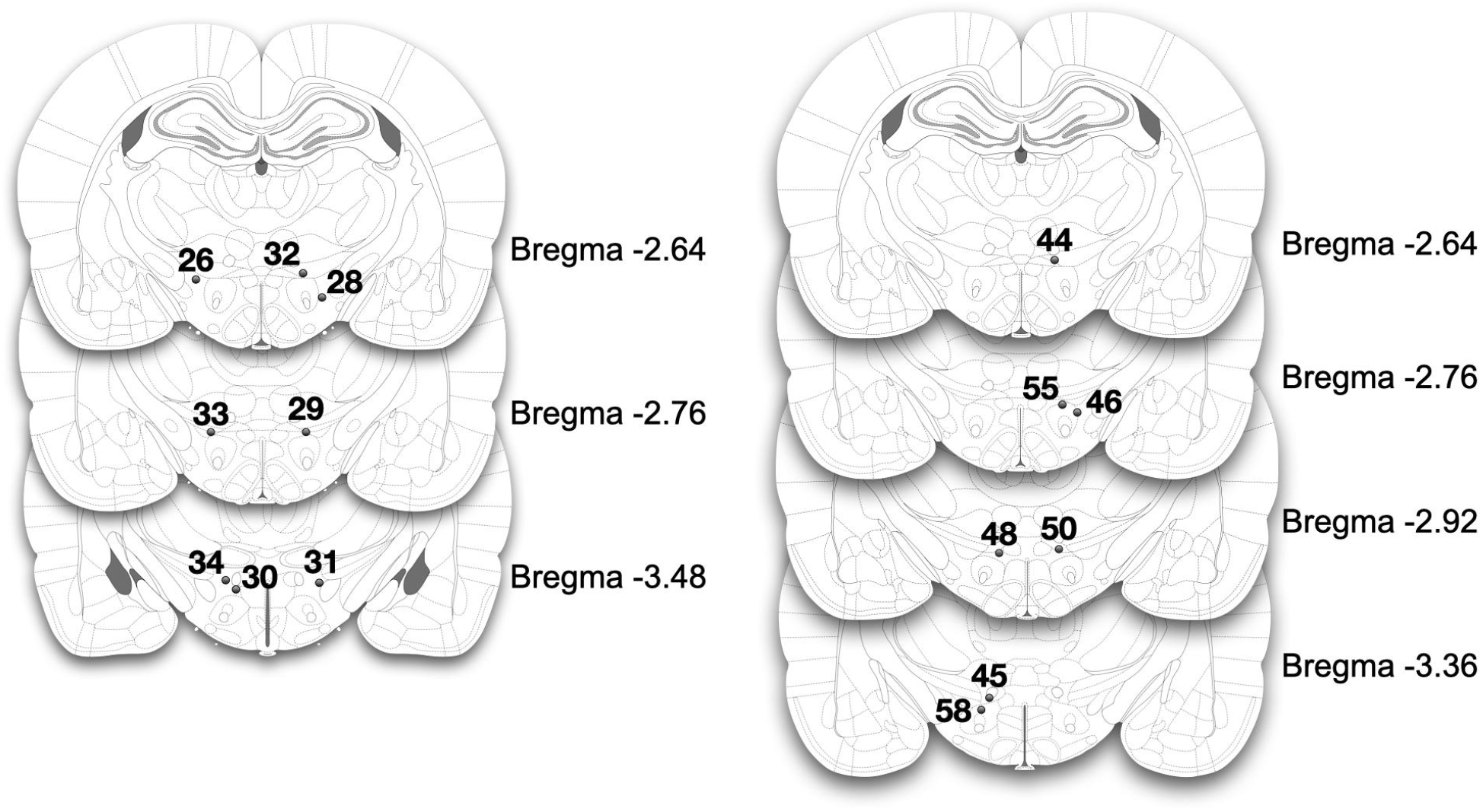
Location of electrode tips of rats in (a) Experiment 1 and (b) Experiments 2 and 3. Each electrode tip was located within or adjacent to the boundary of the LH level of the MFB, as determined by the Paxinos and Watson (2007) atlas. The numbers next to the filled circles identify the rats. Elsewhere in the paper, these numbers are prefixed by ‘AM.’” Due to a problem in tissue collection, the electrode placement for rat AM47 (Experiment 2) is missing.

**Figure 2.**
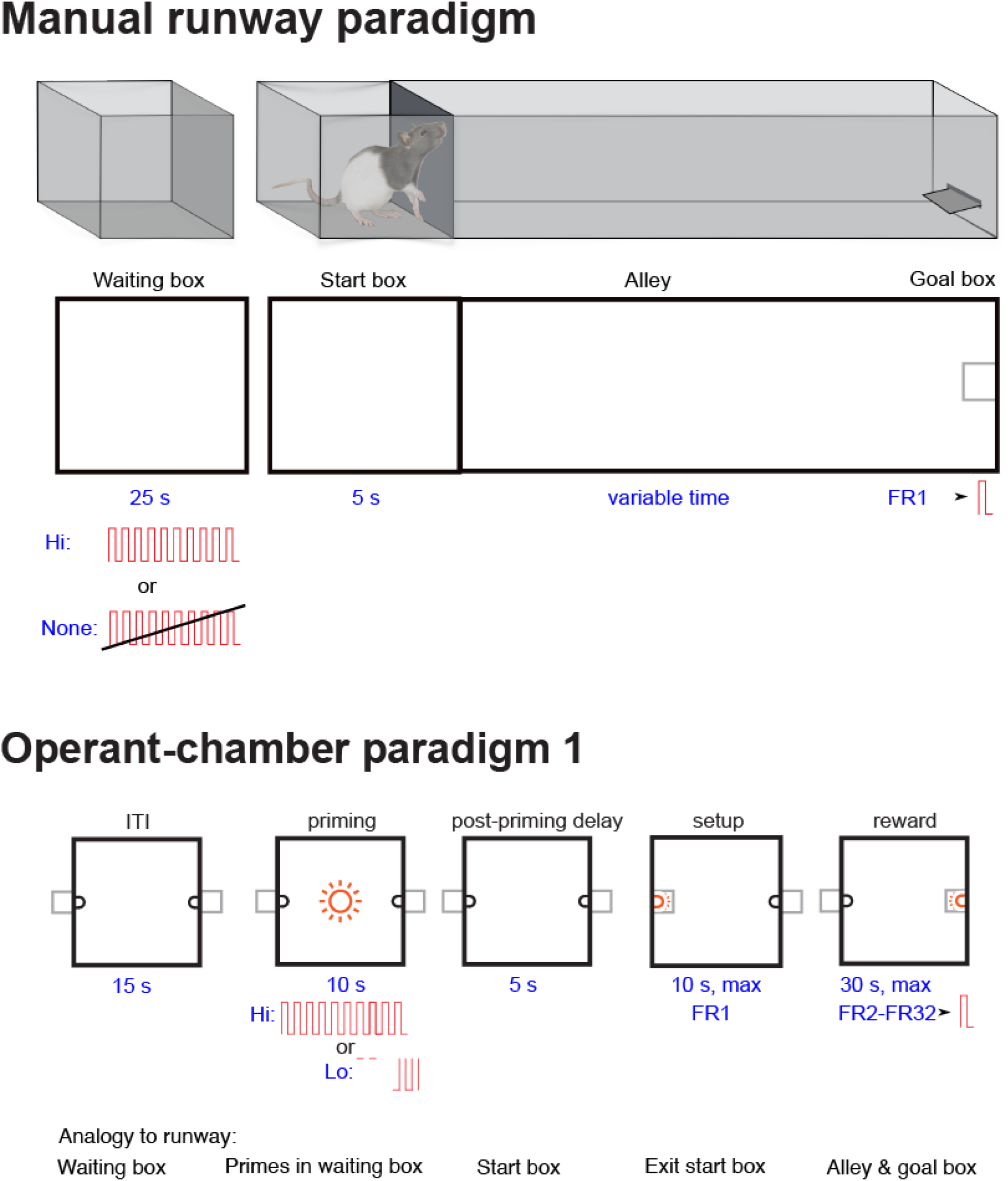
A schematic comparing the manually-operated runway paradigm used by Gallistel and colleagues (top) and the operant-chamber adaptation of the priming paradigm used in Experiment 1 (bottom). That adaptation was designed to be roughly analogous to the runway paradigm. The setup lever in the operant-chamber paradigm is shown on the left and the stimulation lever on the right.

**Figure 3.**
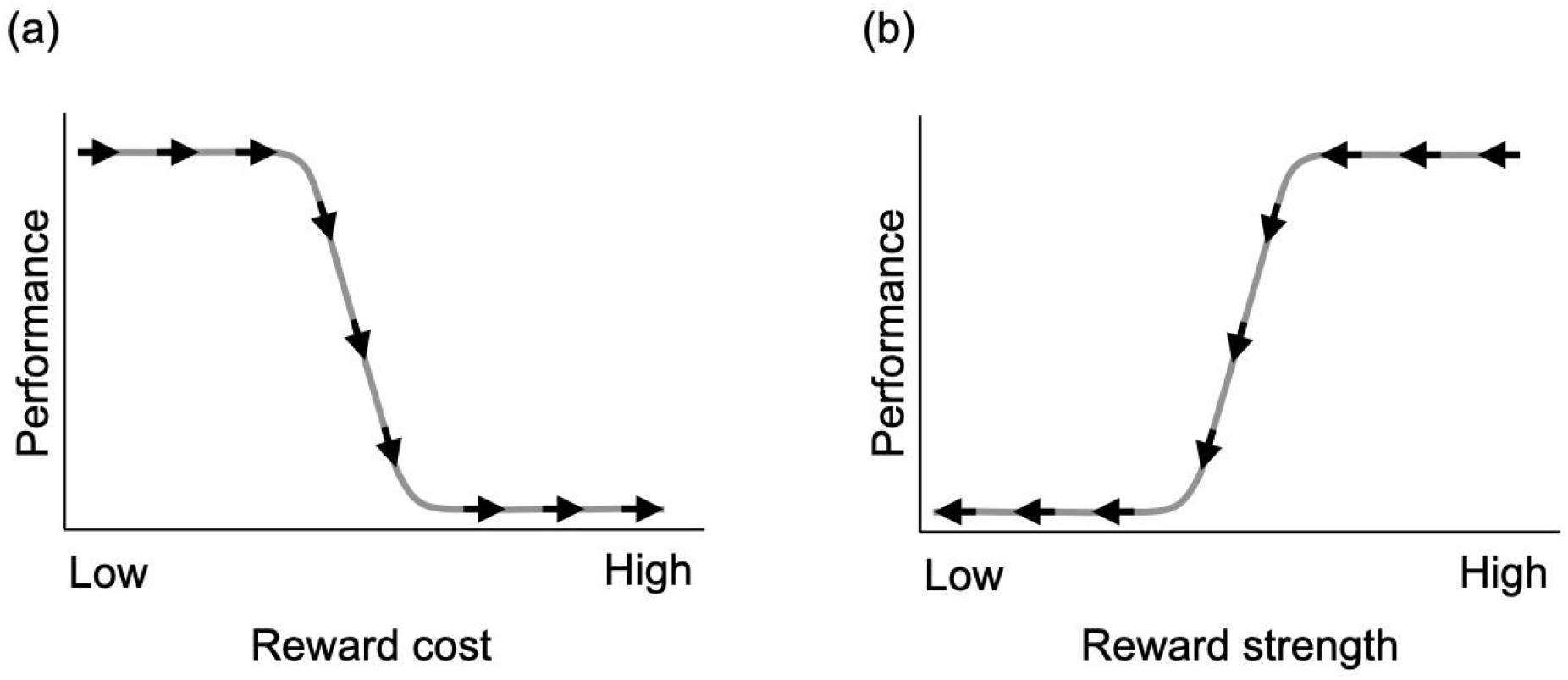
Schematic of the testing procedures in experiment 1. (a) To obtain a rate-cost curve, the stimulation strength was set to a high value and the cost systematically increased across blocks, as illustrated by the arrows pointing from low to high reward cost. (b) To obtain a rate-frequency curve, the cost was set to a constant, low value, and the stimulation strength systematically declined across blocks, as illustrated by the arrows pointing from high to low reward strength. Each arrow in the figures represents a block of trials.

**Figure 4.**
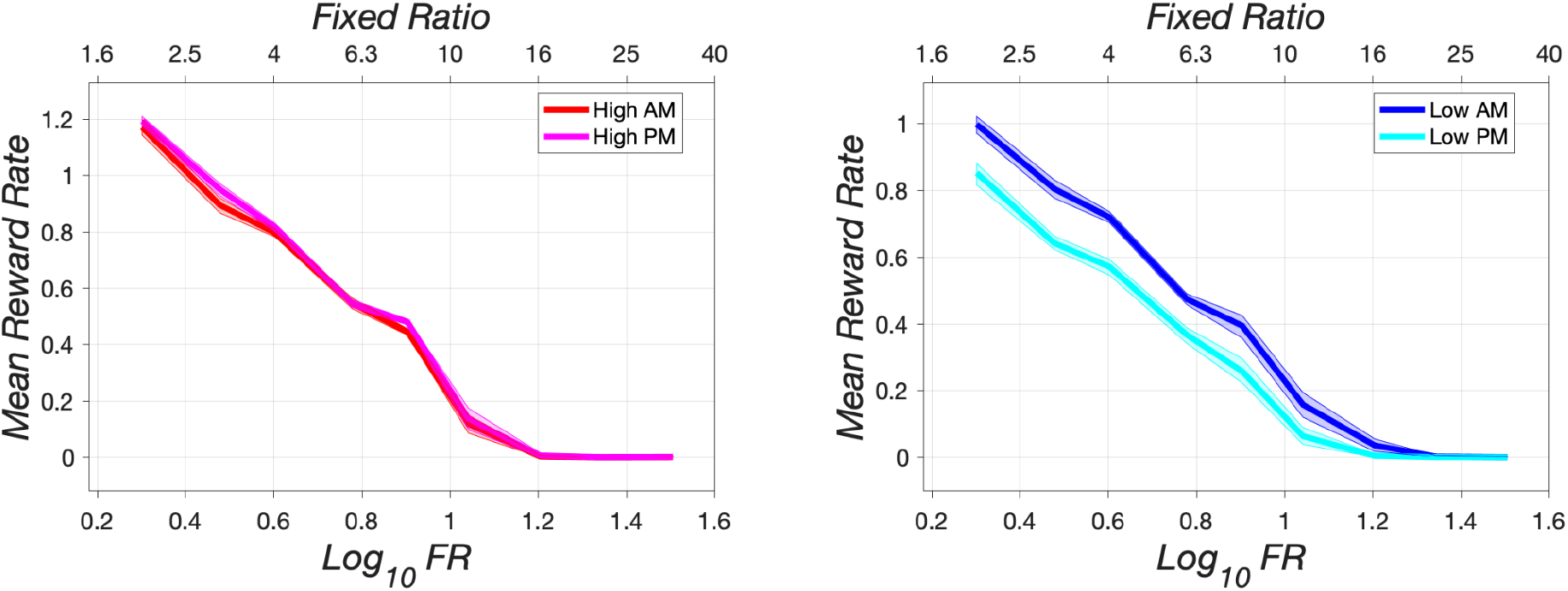
Mean reward rate as a function of reward cost (fixed-ratio requirement) in morning (AM) and afternoon (PM) test sessions. Reward rates are lower in afternoon sessions than in morning sessions under low priming (right panel) but similar in the two sessions under high priming (left panel). The error band shows the 95% CIs around the robust mean. Data are from rat AM26.

**Figure 5.**
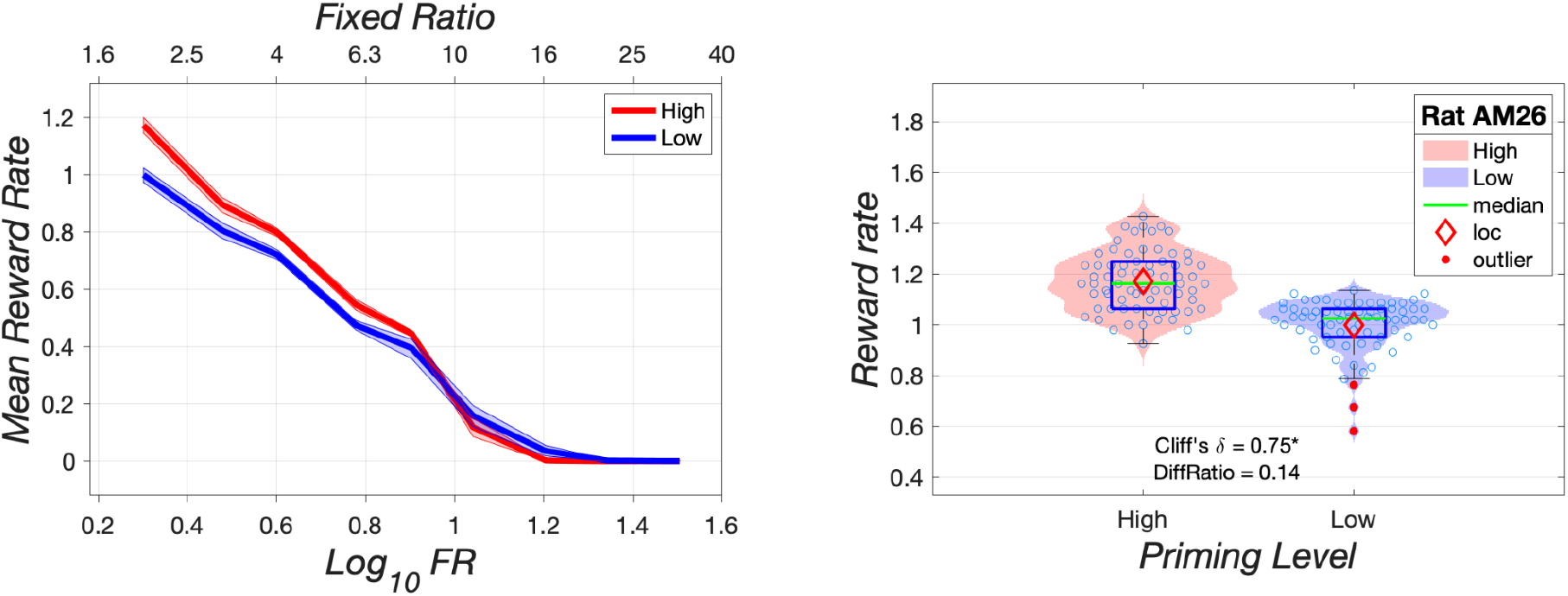
Effect of reward cost (FR requirement) on reward rate under high (red) and low (blue) priming. The violin plots in the right panel show the kernel density of the data obtained at the lowest fixed-ratio tested (FR2). Individual observations are denoted by open circles; their lateral position has been jittered to minimize overlap. The red diamond denotes the robust mean (**loc**ation parameter of the Tukey bisquare estimator), the green line denotes the median, the vertical extent of the blue rectangle shows the interquartile range, and filled red circles denote outliers. Data are from morning sessions carried out with rat AM26.

**Figure 6.**
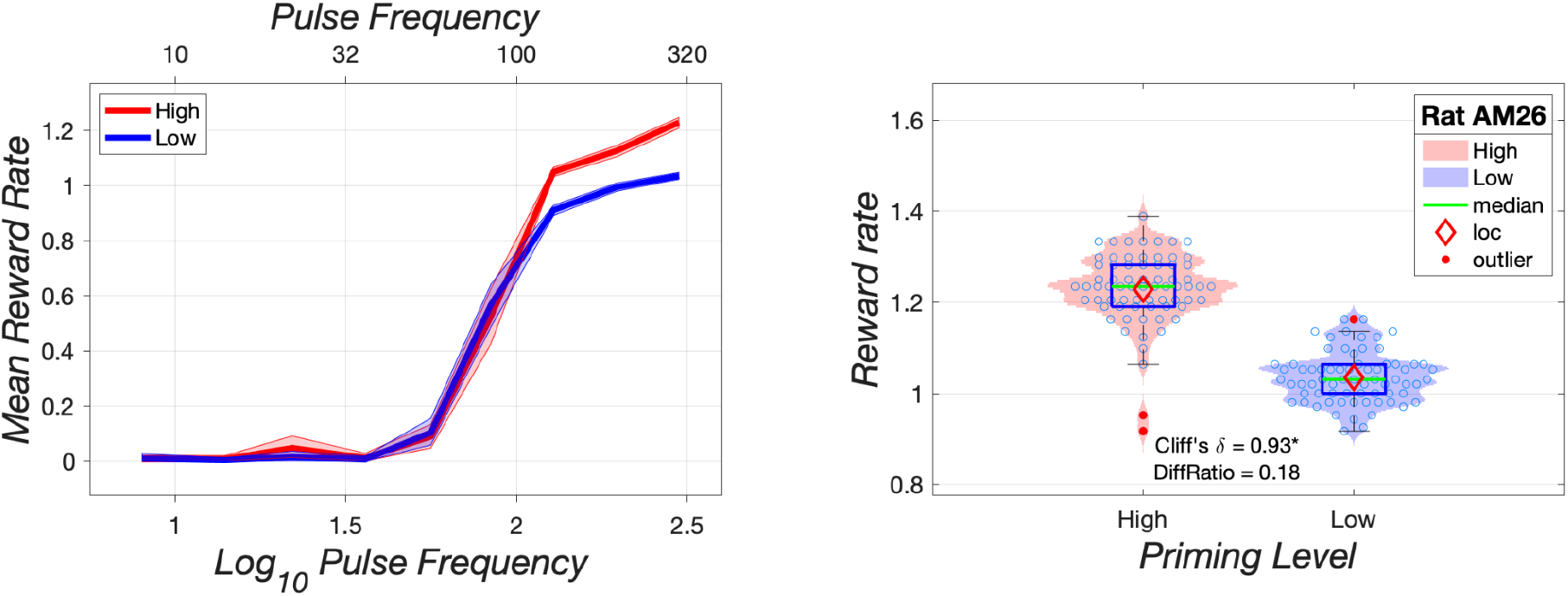
Effect of stimulation strength (pulse frequency) on reward rate under high (red) and low (blue) priming. Please see the Figure 5 caption for details.

**Figure 7.**
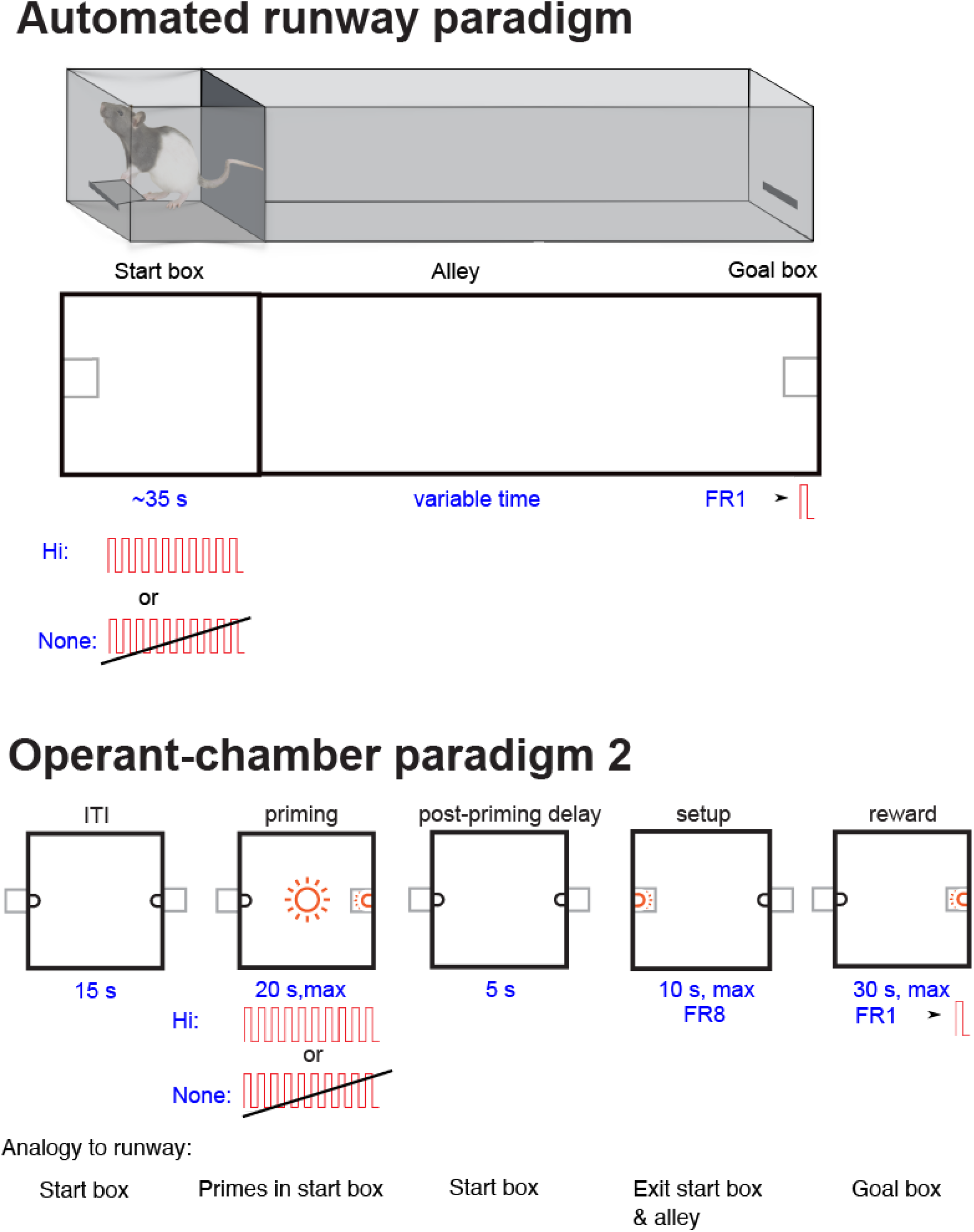
Schematic representation of Gallistel’s automated-runway paradigm (top) and our revised adaptation of the Gallistel’s paradigm for use in standard operant-conditioning chambers. The setup lever is shown on the left and the stimulation lever on the right.

**Figure 8.**
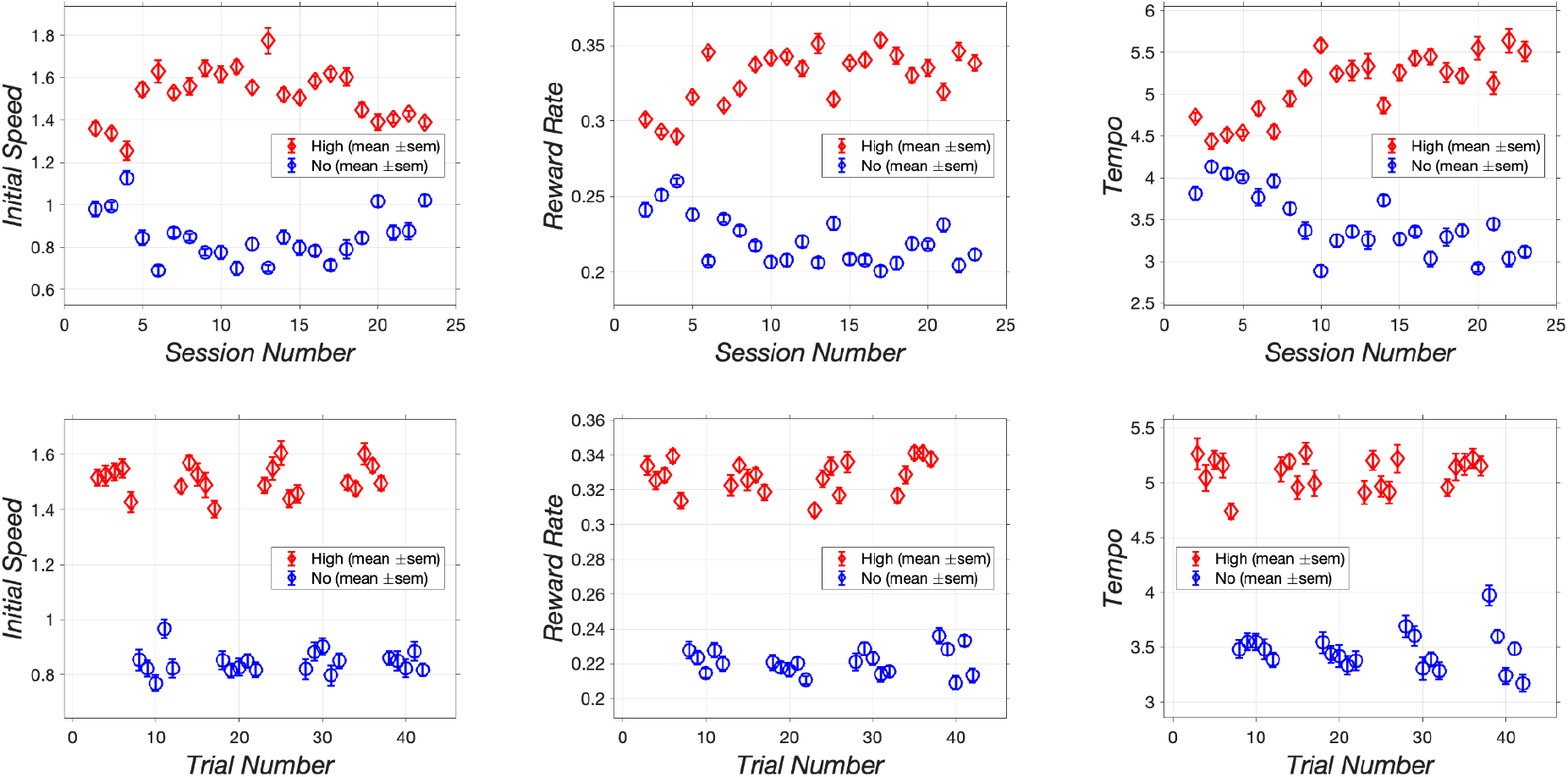
Effect of high (red diamonds) versus no (blue circles) priming as a function of session number collapsed across trials (top) and trial number collapsed across sessions (bottom). The symbols represent robust means, and the error bars represent robust standard errors of the mean. Data are from rat AM46.

#### Apparatus

The operant chambers (34 cm long x 24 cm wide x 66 cm high) were composed of wire-mesh floors (8 cm above the base), a transparent Plexiglas^®^ front panel, an amber house light (10 cm above the mesh floor), and two retractable levers (ENV-112B, MED Associates, St. Albans, Vermont, USA). A lever was located on the left and right sides of the box, and a cue light (1 cm) positioned 4 cm above each lever. An electrical rotary joint centered at the top of the box allowed animals to move freely with the stimulation leads.

The temporal parameters of the electrical stimulation and pulse amplitude were determined by a computer-controlled digital pulse generator and constant-current amplifier, respectively. Experiments were controlled by, and data were collected with, a custom-written computer program (“PREF3”, Steve Cabilio, Concordia University, Montreal, QC, Canada).

#### Procedures

##### Training

Rats were each screened to determine which electrode, electrical current, and pulse frequency promoted vigorous lever pressing with minimal to no disruptive stimulation-induced movements. The rats in these experiments responded for currents between 320 microamperes (μA) to 400 μA. The rewarding stimulation consisted of a single 0.5-s train of 0.1-ms cathodal pulses. The maximum pulse frequency of the stimulation that served as the reward ranged from 266 pulses per second (pps) to 320 pps. The pulse frequency of the priming stimulation was lower, ranging from 198 pps to 222 pps. Consecutive trains of high-frequency priming stimulation produced forced movements, and reducing the pulse frequency served to minimize those disruptive side effects. The settings determined for each rat were used throughout the experiment. Once the operant behavior was stable, rats were trained to press a setup lever, armed on a fixed-ratio 1 (FR1) schedule, that did not deliver reward but instead activated the extension of a stimulation lever located at the opposite side of the operant-conditioning chamber. The stimulation lever was armed on an FR schedule that ranged from two to 32 presses, and completion of this response requirement earned the rat a reward.

To obtain curves expressing response rate as a function of stimulation strength, the response requirement was held constant at FR2 for all trials while the pulse frequency of the reward was systematically decreased. To obtain curves expressing response rate as a function of effort cost, the response requirement was progressively increased while the strength of the reward was held constant at a high value.

##### Testing: Trial structure

We designed our procedures to be roughly analogous to those employed in manual runway testing. The top of Figure 2 depicts the test procedures used in one version of the runway paradigm (C. R. Gallistel et al., 1974). A rat was placed in a waiting box during the 25-s inter-trial interval (ITI). During the last 10 s of the ITI, the rat received 10 trains of priming stimulation. Immediately after the last train of priming stimulation, the rat was transferred to the start box. Five seconds later, the start box door dropped to allow access to the alley. The rat could then run to the goal box at the end of the alley, where a single lever press delivered a train of rewarding brain stimulation.

The bottom of Figure 2 depicts our adaptation of Gallistel’s priming paradigm for use in standard operant-conditioning chambers. A trial cycle started with a 15-s ITI, during which the levers were retracted, and the house light was off. The ITI was followed by a 10 s priming phase during which the amber house lights flashed (10 cycles of 0.5s on, 0.5s off), and priming stimulation was delivered. The priming stimulation consisted of either 10 (“high priming”) or two (“low priming”), non-contingent 0.5-s trains of 0.1 ms cathodal pulses, delivered at a rate of one train per second. On high-priming trials, delivery of the priming stimulation commenced at the start of the priming phase. On low-priming trials, primes were delivered at the eight-s mark of the priming phase. Following the priming phase there was a five-s delay, and then the setup phase began. A single response on the setup lever caused that lever to retract and the stimulation lever to extend, ending the setup phase and initiating the reward phase. The stimulation lever was armed on an FR schedule during the reward phase. Completion of the response requirement on the stimulation lever earned the rat a single train of reward, caused the stimulation lever to retract, and initiated a new ITI. If the response requirement was not satisfied within 30 s, the stimulation lever was retracted, and a new ITI was initiated.

The ITI and priming period in the operant-chamber paradigm can be viewed as analogous to time spent in the waiting box of Gallistel’s runway. The post-priming delay in the operant-chamber paradigm can be viewed as analogous to the sum of the time required to transfer the rat from the waiting box to the runway start box and the subsequent delay until the opening of the start-box door. The time spent approaching and pressing the setup lever in the operant-chamber paradigm can be viewed as analogous to the time between the opening of the start-box door and the exit of the rat from the runway start box. The time spent fulfilling the stimulation-lever response requirement in the operant-chamber paradigm can be viewed as analogous to the time spent running down the alley and pressing the goal-box lever in the runway paradigm.

##### Testing: Block structure

A set of 15 trials was called a block. The stimulation strength or cost remained constant within a block and was varied systematically across blocks (Figure 3a & b). For example, to measure response rate as a function of stimulation strength, pulse frequency was systematically decreased after every block of trials. Each block is denoted by an arrow in Figure 3. Each test session consisted of 10 blocks of trials, totaling 150 trials. A 2-3 minute warm-up session preceded each test session. During the warm-up, the rats could earn trains of rewarding stimulation by lever pressing on an FR2 schedule.

During a single test day, rats underwent one test session in the high-priming condition and a second test session in the low-priming condition. The order of the priming conditions was counterbalanced across days, with the high-priming session run first on odd-numbered days and the low-priming session run first on even-numbered days.

##### Rate-Cost Curves

To obtain curves expressing response rate as a function of reward cost, the ratio requirement was increased systematically (“swept”) while the pulse frequency was held at a constant high value (Figure 3a). Cost was lowest in the first block of trials (FR2) and increased across blocks until the highest cost was reached (FR32). The first block of trials and the first trial of each block served as a warm-up block and a learning trial, respectively, and were excluded from analyses.

##### Rate-Frequency Curves

To measure the dependence of response rate on stimulation strength, the pulse frequency was systematically decreased (“swept”) while the response requirement was held constant at FR2 (Figure 3b). Data from the first block of trials and the first trial of each block were treated as a warm-up block and a learning trial, respectively, and were excluded from analyses. The highest pulse frequency was in effect in both the first and second block; the pulse frequency was then reduced in equal proportional steps across the remaining blocks. The step size between pulse frequencies was adjusted for each rat to capture the sigmoidal shape of the rate-frequency curve. The step sizes used in this experiment ranged from 0.08 to 0.18 common logarithmic units. The pulse frequencies were chosen to yield well-formed sigmoidal curves with multiple data points positioned along clearly defined lower and upper asymptotes and the intervening rising segment.

#### Statistical Analyses

For each rat, data were analyzed for each type of curve (rate-cost, rate-frequency) and each priming condition (high, low). Data were analyzed and graphs were plotted using custom-written MATLAB (The Mathworks, Natick, MA) scripts.

##### Reward Rate

Reward rate served as the measure of response vigor. Reward rate is the inverse of the total time (s) elapsed between the start of the setup phase and the delivery of the reward. Higher reward rates reflect more vigorous reward pursuit. The transformation of elapsed time to rate yields a measure analogous to the running speeds reported in Gallistel’s investigations on the priming effect of electrical brain stimulation (Edmonds et al., 1974; Edmonds & Gallistel, 1974; C. R. Gallistel et al., 1974).

To guard against undue influence of outliers, we used the robust mean as the measure of central tendency. The location parameter returned by Tukey’s bisquare estimator (Hoaglin et al., 2000) was computed using the robustfit function in the MATLAB Statistics Toolbox; this value was used as the robust mean.

##### Distributions

The distribution of the data was visualized with violin plots based on kernel-density estimation (KDE) (Jonas, 2022). This is a non-parametric method for estimating the probability-density function of a random variable, such as reward rate. In contrast to traditional statistics, KDE does not assume a particular probability-density function (e.g., a normal distribution) for the data. Instead, KDE creates a smoothed data distribution using a kernel function. In the present experiment, a Gaussian kernel was employed for this purpose.

##### Effect size

Visual inspection of distribution plots show that the speed measure is often skewed and/or bimodal. Thus, we used Cliff’s Delta (Cliff, 1993) as the measure of effect size. Cliff’s Delta is an effect-size measure used for ordinal data; it is free from assumptions about the form of the data distribution. This statistic provides the difference between the probability that a score in the high-priming condition exceeds a score in the low-priming condition and the probability that a score in the low-priming condition exceeds a score in the high-priming condition. A Cliff’s Delta value reliably greater than zero indicates that faster response speeds were more prevalent following high than low (or no) priming; a value of one is attained when all high-priming values exceed the largest low-priming value (i.e, there is no overlap). A Cliff’s Delta value of zero is obtained when the probabilities of greater speeds in the high- and low-priming conditions are equal, whereas negative values indicate that priming reduces response speeds (with Cliff’s Delta equal to -1 when all low-priming values exceed the largest high-priming value.) Bootstrapping (resampling with replacement) (Efron & Tibshirani, 1993) was used to calculate both Cliff’s Delta and its surrounding 95% confidence interval (CI).

##### Difference Ratio

The effect-size statistic used here, Cliff’s Delta, assesses the overlap between the scores obtained in the high- and low-(or no-) priming conditions. When the variance within the priming condition is very low, Cliff’s Delta (and other common effect-size statistics) can assume high values even when the difference between the central tendencies of the high- and low-priming distributions is small. An additional statistic was developed to express the normalized difference between the central tendencies of the vigor measures obtained in the high- and low-priming conditions independently of the dispersion of the data. Instead of scaling the difference by the pooled standard deviation of the data, as is the case for the commonly-used Cohen’s *d* statistic, we scaled the difference by the central tendency of the pooled distribution across priming conditions. In Experiment 1, the robust mean was used as the measure of central tendency in order to guard against undue influence of outliers. The resulting “Difference Ratio” expresses the discrepancy between the priming-condition robust means as a percentage of the grand robust mean across priming conditions. This normalizes the statistic across sessions and priming conditions. We view the difference ratio as an indication of the meaningfulness of the difference between the robust-mean reward rates in the high- and low-priming conditions. Given sufficiently low within-priming-condition dispersion, a 1% difference between the robust means may be statistically reliable (as assessed by Cliff’s Delta or other effect-size measures), but we do not regard such a small difference as meaningful.

##### Criteria for a priming effect

A two-criterion approach was used to determine whether there is a reliable and meaningful difference between performance in the high- and low-priming conditions. The first criterion was a Cliff’s Delta value greater than zero surrounded by a 95% CI excluding zero. The second criterion was a Difference Ratio equal to or greater than .05. A difference that met the criterion for Cliff’s Delta but not the Difference-Ratio criterion was considered statistically reliable but too small to be regarded as meaningful.

### Experiment 1: Results

As expected, response vigor depended on reward cost. However, this dependence varied as a function of the order in which daily high- and low-priming sessions were run and of the number of priming trains delivered (low priming: two trains; high priming: ten trains).

#### The Session-Order Effect

The right panel of Figure 4 compares rate-cost curves obtained from rat AM26 under low priming in the morning (blue) and afternoon sessions: Response vigor (reward rate) was systematically lower in the afternoon session. In contrast (left panel), similar rate-cost curves were obtained from this rat under high priming in the morning (red) and afternoon (magenta) sessions. Graphs for all rats are shown in Figures S1-S2 and S3-S4, for the reward-cost and reward-strength sweeps, respectively.

Table 1 shows comparisons between reward rates obtained in the morning and afternoon sessions for all rats under low priming. Cliff’s Delta is positive, with a median value of 0.705; the surrounding CI excludes zero in the data from all eight rats. This indicates that there is a statistically reliable difference between the ranks of the reward rates observed in the morning and afternoon sessions under low priming. The Difference Ratio ranges from 0.103 - 0.333, with a median value of 0.192. This means that the robust-mean reward rates in the morning session were 5.15% - 16.67% higher than the grand robust mean of the pooled data from both sessions, and the robust-mean reward rates in the afternoon session were 5.15% - 16.67% lower.

**Table 1.** Effect of session order on reward rates in the lowest-cost trial of the cost sweep under low priming.

| Rat | RobustMeanAM | RobustMeanPM | GrandRobustMean | DiffRatio | Cliffs_delta | Cliffs_delta_CI_hi | Cliffs_delta_CI_lo | met both |
| --- | --- | --- | --- | --- | --- | --- | --- | --- |
| AM26 | 1.022 | 0.870 | 0.937 | 0.163 | 0.727 | 0.849 | 0.576 | TRUE |
| AM28 | 1.077 | 0.956 | 1.009 | 0.120 | 0.693 | 0.828 | 0.533 | TRUE |
| AM29 | 1.110 | 0.786 | 0.969 | 0.333 | 0.585 | 0.676 | 0.493 | TRUE |
| AM30 | 1.044 | 0.852 | 0.949 | 0.203 | 0.799 | 0.886 | 0.700 | TRUE |
| AM31 | 0.816 | 0.681 | 0.739 | 0.181 | 0.631 | 0.734 | 0.511 | TRUE |
| AM32 | 0.935 | 0.843 | 0.891 | 0.103 | 0.450 | 0.598 | 0.292 | TRUE |
| AM33 | 0.735 | 0.558 | 0.636 | 0.278 | 0.717 | 0.820 | 0.603 | TRUE |
| AM34 | 0.937 | 0.731 | 0.825 | 0.249 | 0.860 | 0.927 | 0.780 | TRUE |
| Median | 0.980 | 0.815 | 0.914 | 0.192 | 0.705 | 0.824 | 0.554 |  |
| Max | 1.110 | 0.956 | 1.009 | 0.333 | 0.860 | 0.927 | 0.780 |  |
| Min | 0.735 | 0.558 | 0.636 | 0.103 | 0.450 | 0.598 | 0.292 |  |
| met criterion |  |  |  | 8 |  |  | 8 | 8 |

Under high priming (Table 2), Cliff’s Delta is positive in five of eight cases, with a mean value of 0.124; the surrounding CI excludes zero in four of the eight cases. The Difference Ratio (range: -0.046 - 0.110) exceeds 0.05 in only three cases, with a median value of 0.023.

**Table 2.** Effect of session order on reward rates in the lowest-cost trial of the cost sweep under high priming.

| Rat | RobustMeanAM | RobustMeanPM | GrandRobustMean | DiffRatio | Cliffs_delta | Cliffs_delta_CI_hi | Cliffs_delta_CI_lo | met both |
| --- | --- | --- | --- | --- | --- | --- | --- | --- |
| AM26 | 1.170 | 1.198 | 1.184 | -0.024 | -0.157 | 0.013 | -0.318 | FALSE |
| AM28 | 1.172 | 1.190 | 1.185 | -0.016 | -0.050 | 0.153 | -0.250 | FALSE |
| AM29 | 1.196 | 1.252 | 1.224 | -0.046 | -0.193 | -0.059 | -0.329 | FALSE |
| AM30 | 1.111 | 1.049 | 1.080 | 0.057 | 0.187 | 0.318 | 0.051 | TRUE |
| AM31 | 0.956 | 0.882 | 0.920 | 0.081 | 0.464 | 0.597 | 0.324 | TRUE |
| AM32 | 0.972 | 0.962 | 0.968 | 0.010 | 0.074 | 0.229 | -0.081 | FALSE |
| AM33 | 0.769 | 0.741 | 0.756 | 0.036 | 0.174 | 0.325 | 0.015 | FALSE |
| AM34 | 0.992 | 0.889 | 0.940 | 0.110 | 0.569 | 0.682 | 0.442 | TRUE |
| Median | 1.052 | 1.006 | 1.024 | 0.023 | 0.124 | 0.274 | -0.033 |  |
| Max | 1.196 | 1.252 | 1.224 | 0.110 | 0.569 | 0.682 | 0.442 |  |
| Min | 0.769 | 0.741 | 0.756 | -0.046 | -0.193 | -0.059 | -0.329 |  |
| met criterion |  |  |  | 3 |  |  | 4 | 3 |

We subtracted the Difference-Ratio and Cliff’s-Delta values in Table 1 (low priming) from those in Table 2 (high priming) to produce Table 3, which thus shows the relative strength of the session-order effect under high and low priming. That table shows that the session-order effect is systematically greater in the case of low priming than high priming. The Cliff’s Delta scores are negative and exclude zero in all cases; the Difference Ratios exceed 0.05 in all cases as well. A similar but somewhat weaker pattern is seen in the pulse-frequency sweeps (Figures S3-S4; Tables S1-S3).

**Table 3.** Differences in the effect of session order on reward rates in the lowest-cost trial of the cost sweep under high and low priming (high-low).

| Rat | DiffRatio | Cliffs_delta | Cliffs_delta_CI_hi | Cliffs_delta_CI_lo | met both |
| --- | --- | --- | --- | --- | --- |
| AM26 | -0.187 | -0.883 | -0.835 | -0.894 | TRUE |
| AM28 | -0.135 | -0.743 | -0.675 | -0.783 | TRUE |
| AM29 | -0.379 | -0.778 | -0.735 | -0.822 | TRUE |
| AM30 | -0.146 | -0.612 | -0.568 | -0.649 | TRUE |
| AM31 | -0.101 | -0.167 | -0.136 | -0.187 | TRUE |
| AM32 | -0.093 | -0.376 | -0.368 | -0.373 | TRUE |
| AM33 | -0.242 | -0.543 | -0.495 | -0.588 | TRUE |
| AM34 | -0.140 | -0.291 | -0.245 | -0.338 | TRUE |
| Median | -0.143 | -0.578 | -0.531 | -0.618 |  |
| Max | -0.093 | -0.167 | -0.136 | -0.187 |  |
| Min | -0.379 | -0.883 | -0.835 | -0.894 |  |
| met criterion | 8 |  |  | 8 | 8 |

The fact that the session-order effect is stronger under low than high priming confounds the results obtained in the afternoon sessions. Differences between high and low priming in those sessions reflect some combination of the priming effect and the session-order effect. To avoid that confound, the remaining analyses are based only on the data obtained in the morning sessions.

#### The Priming Effect as a Function of Reward Cost

The curves in Figures 5 and S5-S6 show that response vigor during the morning sessions varied systematically with reward cost: The mean reward rate was highest when the ratio requirement was low and then declined as the ratio requirement increased. Rate-cost curves were roughly similar in shape in all eight rats (Figures S5-S6). High priming boosted performance when the reward cost was low but the priming effect shrank and ultimately disappeared as the ratio requirement increased. In three cases (Figure S5-S6), high priming decreased reward rates at high reward costs, causing the curves to cross.

Table 4 shows effect-size statistics and Difference Ratios for trials conducted at the lowest reward cost, where the priming effect was strongest. Cliff’s Delta on these trials ranged from 0.152 - 0.749, with a median value of 0.343. The surrounding 95% CI excludes zero in seven of eight cases. The Difference Ratio ranged from 0.038 - 0.158, with a median value of 0.069. In six of eight cases, the Difference Ratio was greater than 0.05. Thus, datasets from six of eight rats met both criteria, thereby showing reliable and meaningful increases in the mean reward rate in response to high priming when the reward cost was low. Distributions of the reward rates observed under high and low priming at the lowest reward cost are shown in the right panel of Figure 5 for rat AM26 and in Figures S5-S6 for all rats.

**Table 4.** Priming effect on the lowest-cost trial of the cost sweep (high priming versus low priming).

| Rat | RobustMeanHi | RobustMeanLo | GrandRobustMean | DiffRatio | Cliffs_delta | Cliffs_delta_CI_hi | Cliffs_delta_CI_lo | met both |
| --- | --- | --- | --- | --- | --- | --- | --- | --- |
| AM26 | 1.170 | 1.022 | 1.086 | 0.136 | 0.749 | 0.854 | 0.623 | TRUE |
| AM28 | 1.172 | 1.077 | 1.116 | 0.085 | 0.446 | 0.599 | 0.286 | TRUE |
| AM29 | 1.196 | 1.110 | 1.155 | 0.075 | 0.210 | 0.337 | 0.086 | TRUE |
| AM30 | 1.111 | 1.044 | 1.072 | 0.063 | 0.297 | 0.459 | 0.120 | TRUE |
| AM31 | 0.956 | 0.816 | 0.894 | 0.158 | 0.628 | 0.731 | 0.512 | TRUE |
| AM32 | 0.972 | 0.935 | 0.959 | 0.038 | 0.152 | 0.312 | -0.015 | FALSE |
| AM33 | 0.769 | 0.735 | 0.753 | 0.045 | 0.182 | 0.328 | 0.028 | FALSE |
| AM34 | 0.992 | 0.937 | 0.963 | 0.058 | 0.388 | 0.518 | 0.249 | TRUE |
| Median | 1.052 | 0.980 | 1.017 | 0.069 | 0.343 | 0.488 | 0.185 |  |
| Max | 1.196 | 1.110 | 1.155 | 0.158 | 0.749 | 0.854 | 0.623 |  |
| Min | 0.769 | 0.735 | 0.753 | 0.038 | 0.152 | 0.312 | -0.015 |  |
| met criterion |  |  |  | 6 |  |  | 7 | 6 |

In all rats, cost sweeps were carried out prior to pulse-frequency sweeps. Following the latter, the cost sweeps were rerun (Figures S7-S8). The overall pattern of results is quite similar, but several of the priming effects grew upon retest.

#### The Priming Effect as a Function of Stimulation Strength

The rate-frequency curves in Figures 6 and S9-S10 show that response vigor also varied systematically with stimulation strength: The reward rate was maximal when the stimulation was intense (high pulse frequency) and then decreased as the stimulation weakened (low pulse frequency). High priming reliably increased the reward rate at higher pulse frequencies. As the pulse frequency decreased, the boost in performance due to high priming shrank and ultimately disappeared. In four cases (Figure S3), high priming decreased reward rates at lower pulse frequencies, causing the curves to cross.

**Figure 9.**
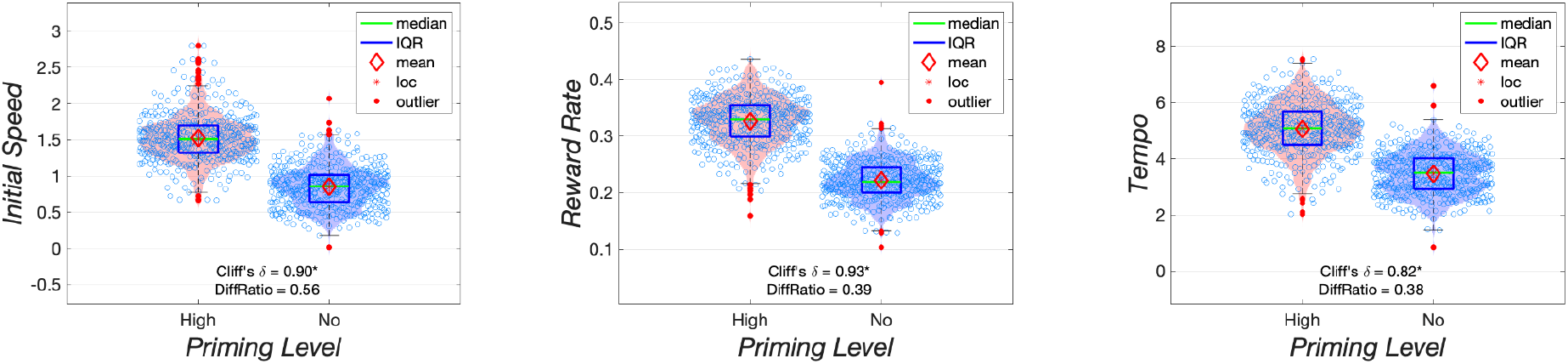
Violin plots showing the distribution of scores on the three measures of vigor. Data are from rat AM46.

**Figure 10.**
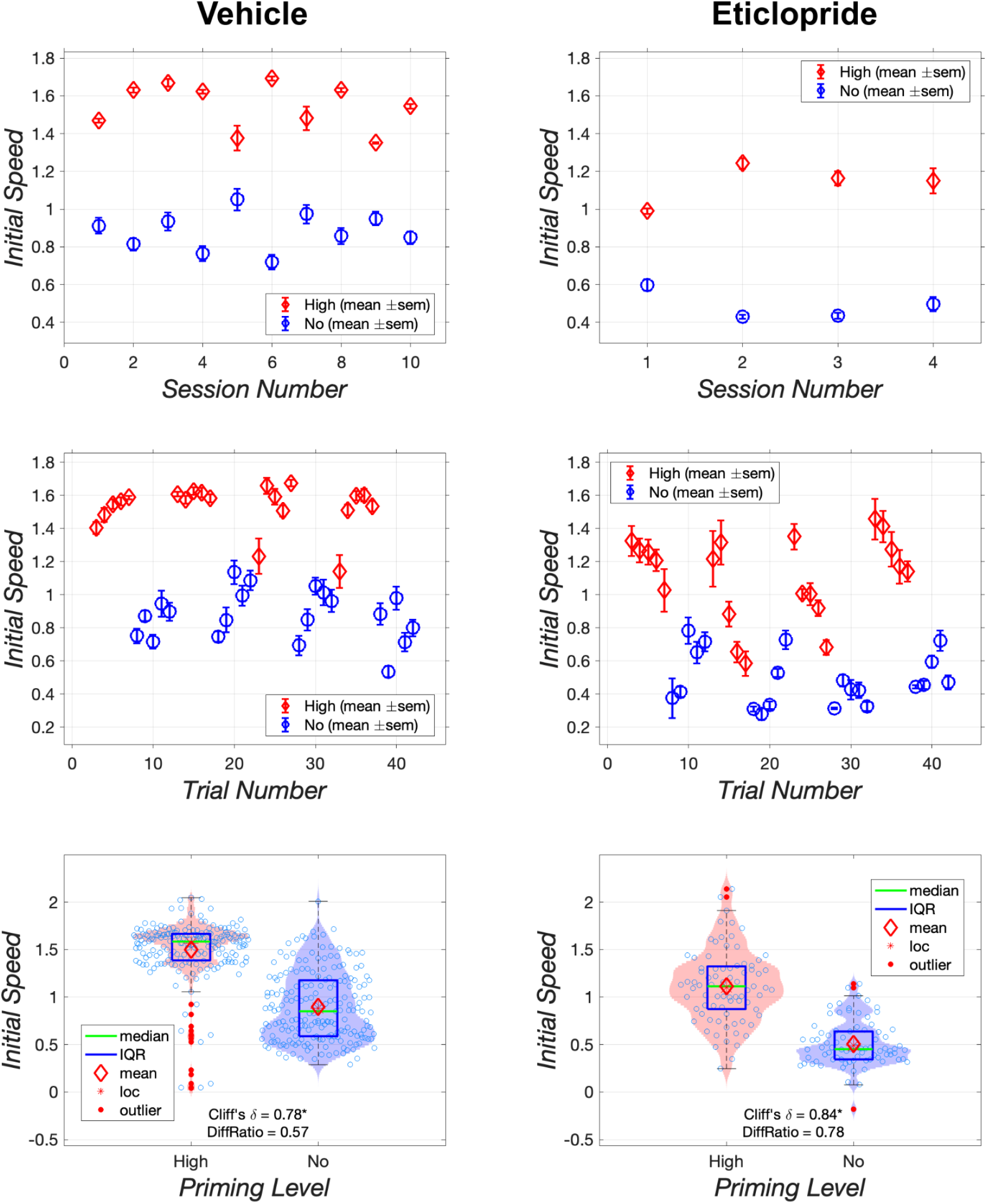
Effect of vehicle and 0.1 mg/kg of eticlopride on the initial speed to initiate responding by rat AM47. The upper row shows results collapsed over trials, whereas the middle row shows results collapsed over sessions. The violin plots in the bottom row show the kernel density of the data. Individual observations are denoted by open circles. The red diamond denotes the robust mean, the green line denotes the median, the vertical extent of the blue rectangle shows the interquartile range, and filled red circles denote outliers.

**Figure 11.**
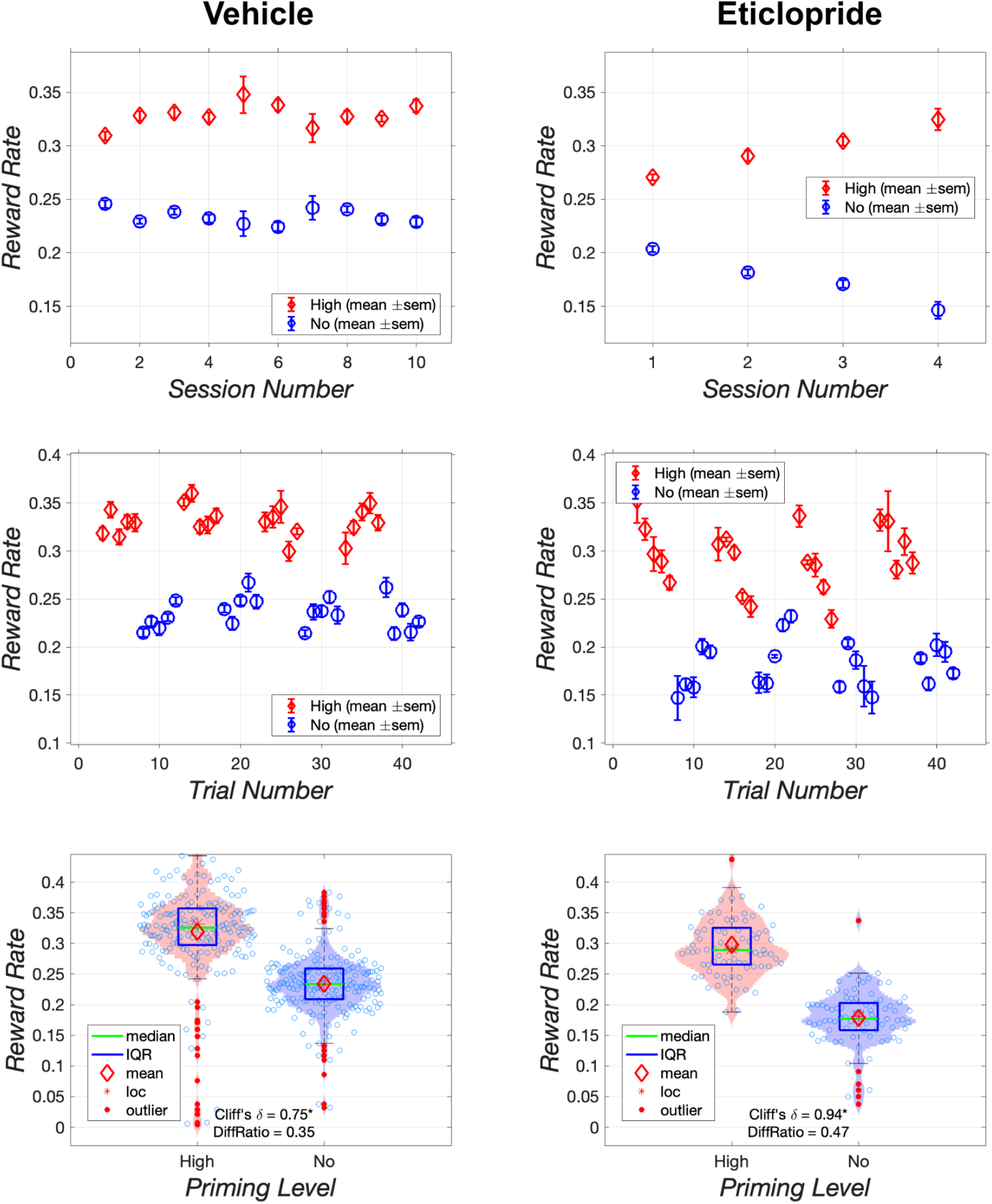
Effect of vehicle and 0.1 mg/kg of eticlopride on reward rate in rat AM47. The upper row shows results collapsed over trials, the middle row shows results collapsed over sessions, and the bottom row shows violin plots of the kernel density of the data.

**Figure 12.**
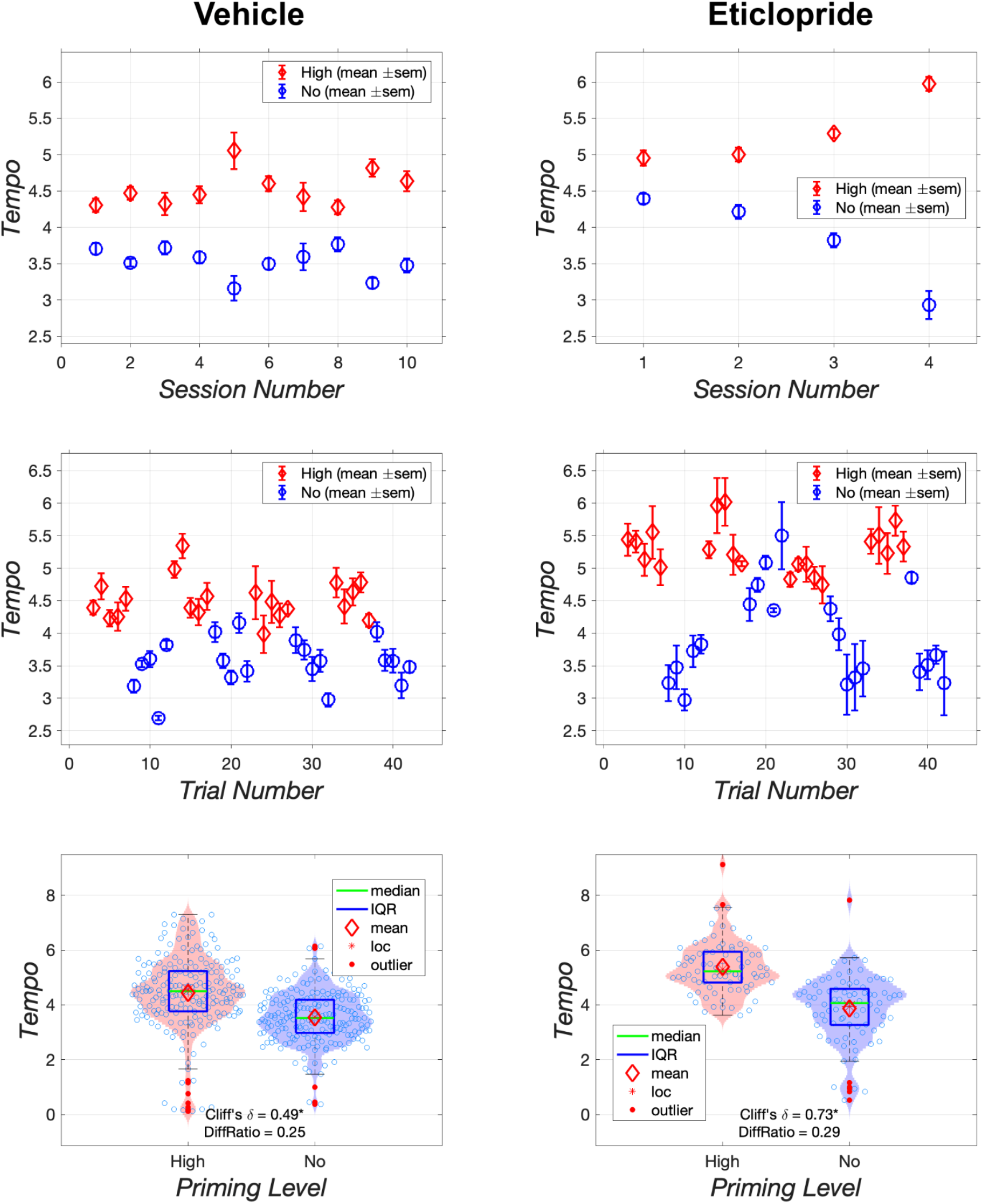
Effect of vehicle and 0.1 mg/kg of eticlopride on the tempo of responding during the setup phase in rat AM47. The upper row shows results collapsed over trials, whereas the middle row shows results collapsed over sessions, and the bottom row shows violin plots of the kernel density of the data.

**Figure 13.**
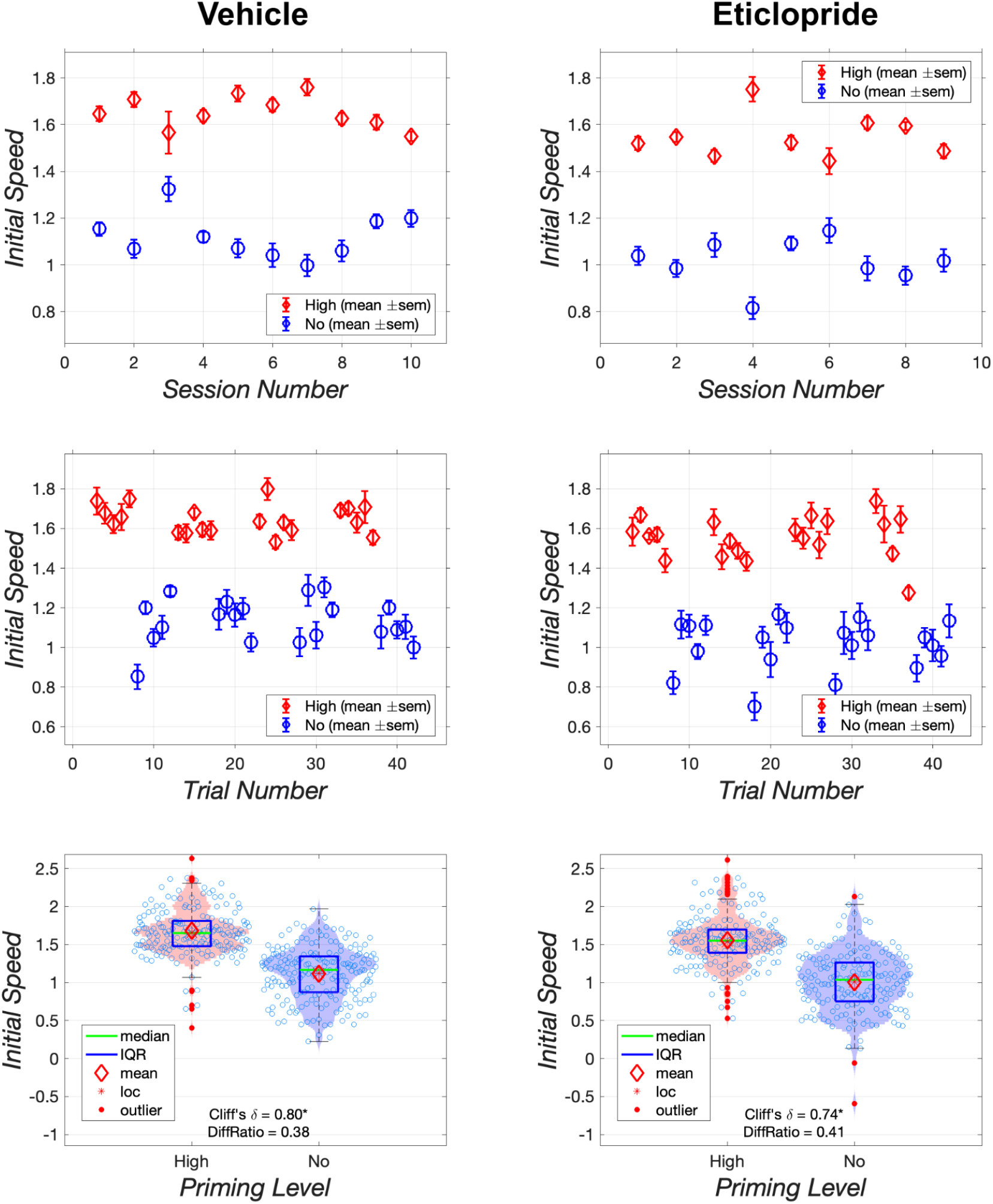
Effect of vehicle and 0.05 mg/kg of eticlopride on the initial speed to initiate responding by rat AM46. The upper row shows results collapsed over trials, the middle row shows results collapsed over sessions, and the bottom row shows violin plots of the kernel density of the data.

**Figure 14:**
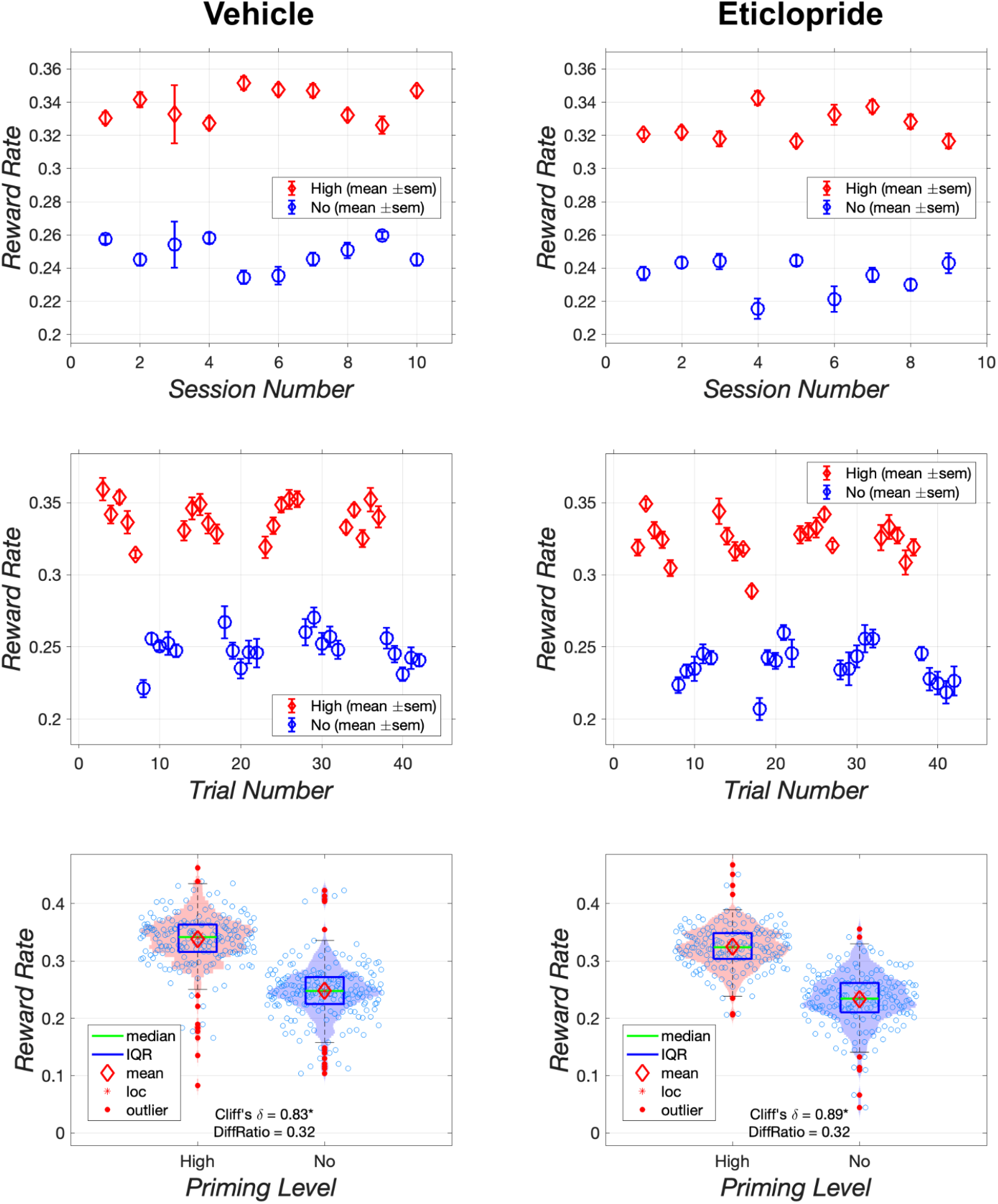
Effect of vehicle and 0.05 mg/kg of eticlopride on reward rate in rat AM46. The upper row shows results collapsed over trials, the middle row shows results collapsed over sessions, and the bottom row shows violin plots of the kernel density of the data.

**Figure 15:**
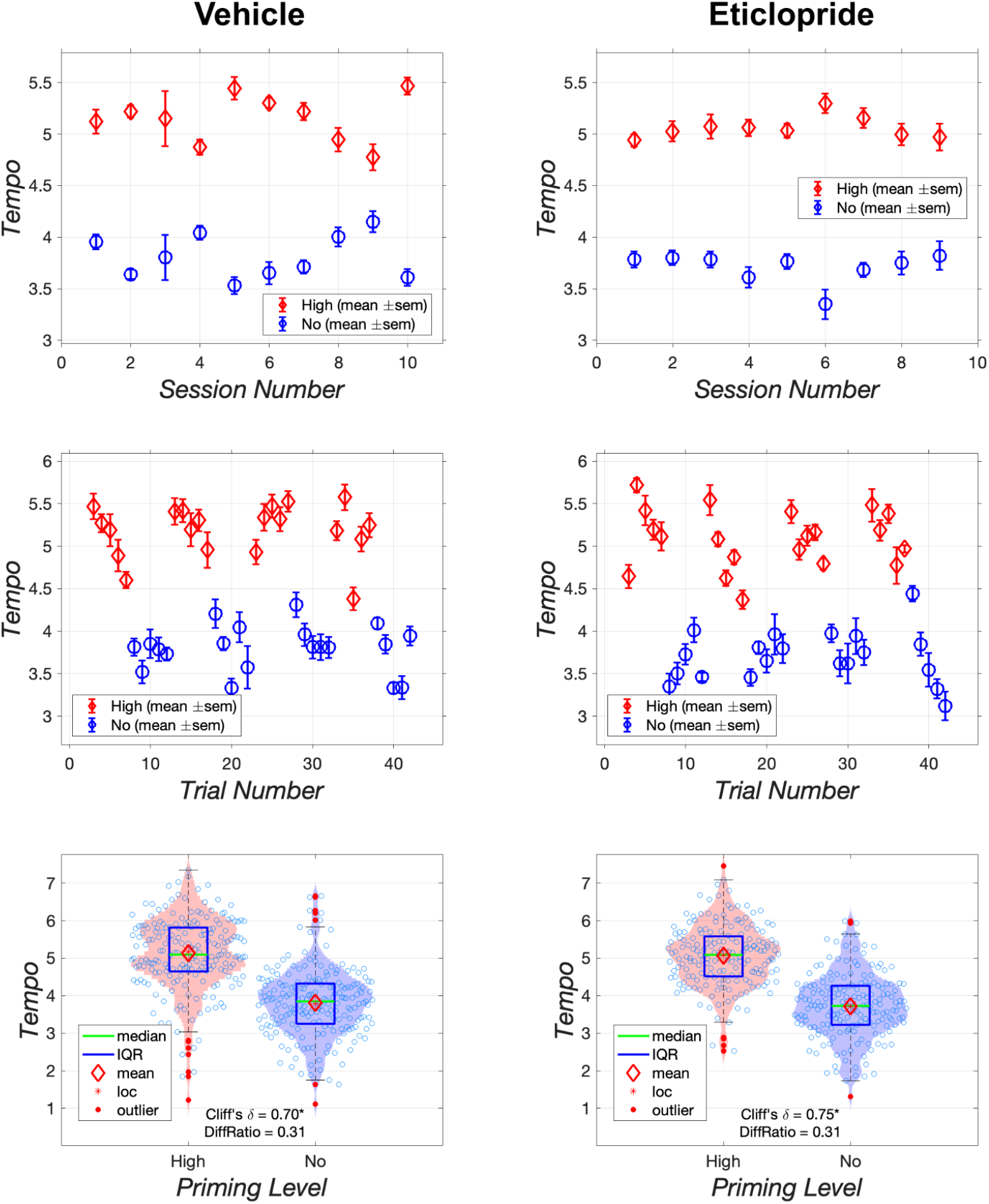
Effect of vehicle and 0.05 mg/kg of eticlopride on the tempo of responding during the setup phase in rat AM46. The upper row shows results collapsed over trials, the middle row shows results collapsed over sessions, and the bottom row shows violin plots of the kernel density of the data.

Table 5 shows effect-size statistics and Difference Ratios for trials conducted at the highest stimulation strength, where the priming effect was strongest. Cliff’s Delta on these trials ranged from 0.304 - 0.928, with a median value of 0.769. The surrounding 95% CI excludes zero in all cases. The Difference Ratio ranged from 0.061 - 0.329, with a median value of 0.199. In all cases, the Difference Ratio was greater than 0.05. Thus, datasets from all eight rats met both criteria, thereby showing reliable and meaningful increases in the mean reward rate in response to high priming when the stimulation strength was highest (Figures 6, S9-S10, Table 5).

**Table 5.** Priming effect on the highest-strength trial of the pulse-frequency sweep (high priming versus low priming).

| Rat | RobustMeanHi | RobustMeanLo | GrandRobustMean | DiffRatio | Cliffs_delta | Cliffs_delta_CI_hi | Cliffs_delta_CI_lo | met both |
| --- | --- | --- | --- | --- | --- | --- | --- | --- |
| AM26 | 1.238 | 1.033 | 1.131 | 0.182 | 0.928 | 0.990 | 0.839 | TRUE |
| AM28 | 1.274 | 1.067 | 1.158 | 0.178 | 0.748 | 0.865 | 0.607 | TRUE |
| AM29 | 1.078 | 0.818 | 0.967 | 0.270 | 0.790 | 0.873 | 0.688 | TRUE |
| AM30 | 1.331 | 0.953 | 1.147 | 0.329 | 0.879 | 0.942 | 0.797 | TRUE |
| AM31 | 0.990 | 0.765 | 0.895 | 0.251 | 0.836 | 0.902 | 0.758 | TRUE |
| AM32 | 0.988 | 0.929 | 0.963 | 0.061 | 0.304 | 0.434 | 0.170 | TRUE |
| AM33 | 0.834 | 0.672 | 0.751 | 0.216 | 0.612 | 0.720 | 0.499 | TRUE |
| AM34 | 0.939 | 0.866 | 0.910 | 0.081 | 0.440 | 0.569 | 0.305 | TRUE |
| Median | 1.034 | 0.898 | 0.965 | 0.199 | 0.769 | 0.869 | 0.648 |  |
| Max | 1.331 | 1.067 | 1.158 | 0.329 | 0.928 | 0.990 | 0.839 |  |
| Min | 0.834 | 0.672 | 0.751 | 0.061 | 0.304 | 0.434 | 0.170 |  |
| met criterion |  |  |  | 8 |  |  | 8 | 8 |

Distributions of the reward rates observed under high and low priming at the highest stimulation strength are shown in the right panel of Figure 6 for rat AM26 and in Figures S9-S10 for all rats.

#### Relative Strength of the Priming Effect on Strength versus Cost Sweeps

On the lowest-cost and highest-strength trials in the cost and pulse-frequency sweeps, the cost and strength of the rewarding train of electrical brain stimulation were identical (FR2, highest pulse frequency the rat could tolerate). Nonetheless, the priming effect was generally stronger when the trial in question composed part of a pulse-frequency sweep than when it composed part of a cost sweep. To evaluate this surprising observation, we subtracted Cliff’s-Delta and difference-ratio values for the cost-sweep case from the values for the pulse-frequency-sweep case, thus producing a table (Table 6) of “differences of differences.” The differences between Cliff’s Delta scores range from 0.052 - 0.582 with a median value of 0.254; all of the surrounding 95% CIs excludes zero. The Difference Ratio scores range from 0.023 - 0.267 with a median value of 0.093. Thus, five of eight datasets meet our criteria for a reliable and meaningful difference.

**Table 6.** Relative strength of the priming effect on the highest-strength trial of the pulse-frequency sweep and the lowest-cost trial of the cost sweep. Reward strength and cost were identical on these trials.

| Rat | DiffRatio | Cliffs_delta | Cliffs_delta_CI_hi | Cliffs_delta_CI_lo | met both |
| --- | --- | --- | --- | --- | --- |
| AM26 | 0.046 | 0.179 | 0.136 | 0.216 | FALSE |
| AM28 | 0.093 | 0.302 | 0.266 | 0.321 | TRUE |
| AM29 | 0.195 | 0.580 | 0.536 | 0.602 | TRUE |
| AM30 | 0.267 | 0.582 | 0.484 | 0.677 | TRUE |
| AM31 | 0.094 | 0.207 | 0.170 | 0.246 | TRUE |
| AM32 | 0.023 | 0.152 | 0.121 | 0.184 | FALSE |
| AM33 | 0.171 | 0.430 | 0.391 | 0.471 | TRUE |
| AM34 | 0.023 | 0.052 | 0.051 | 0.056 | FALSE |
| Median | 0.093 | 0.254 | 0.218 | 0.284 |  |
| Max | 0.267 | 0.582 | 0.536 | 0.677 |  |
| Min | 0.023 | 0.052 | 0.051 | 0.056 |  |
| met criterion | 5 |  |  | 8 | 5 |

### Experiment 1: Discussion

Previous work in the runway paradigm (Edmonds & Gallistel, 1974; C. R. Gallistel et al., 1974; Sax & Gallistel, 1991) shows that the priming effect depends on the strength of the stimulation that serves as the reward; that dependence is replicated here. Experiment 1 extends previous work by showing that the priming effect also depends on the cost of the reward. Below, we discuss these matters as well as implications of Experiment 1 for refining and improving the measurement of priming effects in standard operant-conditioning chambers.

We observed a strong, consistent interaction between session order and the number of priming trains (10 in the high-priming condition; two in the low-priming condition). Reward rates were substantially lower when afternoon low-priming sessions were preceded by morning high-priming sessions than when the low-priming sessions were carried out in the morning. In contrast, the data from only three of eight rats met both of our criteria for a reliable and meaningful difference between reward rates observed when high-priming sessions were carried out in the afternoon (following a morning low-priming sessions) than when high-priming sessions were carried out in the morning. The effect of session order was weaker under high than low priming in all rats (Tables 3, S3; Figures 5, 6; S1-S9).

In the present experiment, afternoon sessions were carried out after the animal had previously completed a morning session. We speculate that some residual fatigue carried over. Perhaps the priming effect that built up over the 10 trains delivered in the high-priming condition was sufficient to overcome such fatigue, but the smaller effect that built up over the two trains delivered in the low-priming condition was not.

Priming was held constant within test sessions (either high or low) and varied across them. The interaction of the session-order effect with the priming condition renders that feature of the design problematic. In Experiments 2 and 3, we minimize this problem by restructuring the test sessions to include alternating blocks of high- and low-priming trials and by detrending the data within sessions.

As predicted by prior experiments carried out in the runway paradigm (Edmonds & Gallistel, 1974; C. R. Gallistel et al., 1974; Sax & Gallistel, 1991), a priming effect in Experiment 1 was manifested only when rewarding stimulation was delivered at the highest pulse frequencies (Figures 6, S8, S9). At lower pulse frequencies, an antagonistic effect of the priming stimulation was uncovered: In half of the subjects, high priming decreased reward rates at low and/or intermediate pulse frequencies (note the crossing curves in Figure S8). An analogous uncovering can also be seen in three of the sets of cost sweeps (Figure S5). This disruptive effect could have arisen from multiple side effects of the automatically delivered priming stimulation including forced movements, superstitious behaviors, and the build-up of aversion (Bower & Miller, 1958; Shizgal & Matthews, 1977). Such side effects would have imposed a handicap on high-priming trials that was overcome only when the rewarding stimulation was particularly intense and inexpensive. We suspect that the disruptive effect could be mitigated by giving the rat control over the timing of the priming stimulation, and we implemented that change in the remaining experiments.

A new finding of this experiment is the dependence of the priming effect on the cost of the rewarding stimulation: The priming effect was manifested only when the fixed-ratio requirements were low (Figures 5, S5, S6). That the priming effect depends on the expectation of reward is well established: Regardless of the strength of the priming, rats typically do not leave the start box of the runway after having learned that rewarding stimulation is no longer available in the goal box. Experiment 1 broadens the domain of the expectation to include reward cost as well as reward intensity. On that view, the priming effect can be seen to arise from the experience of several large, recent rewards (Sax & Gallistel, 1991) given the expectation that a large, inexpensive reward can be earned imminently. Such a broadening of the definition is consistent with the notion of “payoff” (“utility”, “value”) as the ratio of reward intensity to reward cost (Hernandez et al., 2010; Trujillo-Pisanty et al., 2020).

A surprising finding of Experiment 1 is that the priming effect was stronger in the highest stimulation-strength trial of the pulse-frequency sweep than in the lowest fixed-ratio-requirement trial of the cost sweep (Table 6, Figures S5, S8). This result is surprising because the strength and cost of the stimulation are identical on these two trial types. Both types of sweeps progress from “better” to “worse” (i.e, from stronger or cheaper to weaker for pulse-frequency sweeps, from cheaper to or more expensive for cost sweeps). However, there seems to be something about the expectation formed during cost sweeps that moderates the priming effect. We speculate that during repeated cost-sweep testing, the rats learn that arduous work will eventually be required, and this dampens their enthusiasm to seek out additional rewarding stimulation, even on low-cost trials.

Cost sweeps were carried out prior to pulse-frequency sweeps, and the priming effect on cost sweeps appears to grow upon retest in some rats (Figures S5, S7). Thus the order in which the two types of sweeps were run does seem to matter. Nonetheless, the priming effect in the retested cost sweeps (Figure S7) still appears to fall short of the priming effect on pulse-frequency sweeps (Figure S8).

In revising the testing paradigm for use in Experiments 2 and 3, we took advantage of the finding that the priming effect was strongest in the highest stimulation-strength trial of the pulse-frequency sweep and the lowest fixed-ratio-requirement trial of the cost sweep. In the revised paradigm, the cost of the reward delivered by the stimulation lever is always minimal (FR1), and the strength of the stimulation is near the maximum the rat can tolerate without undue side effects. Responding at a relatively high fixed-ratio requirement is dissociated from the reward lever and moved to an earlier phase of the trial, in which it becomes analogous to the work entailed in traversing the runway (Figure 7).

## Experiment 2

Findings of Experiment 1 were applied in Experiment 2 to improve and extend our paradigm for measuring priming effects in standard operant-conditioning chambers. Stimulation was eliminated from the low-priming condition (now called the “no-priming” condition), thus increasing its contrast with the high-priming condition. Priming in the latter condition was self-administered by the rats in the hopes of minimizing disruptive side-effects of the stimulation. The testing paradigm (Figure 7) was rendered more strictly analogous to the automated runway paradigm used by Gallistel et al. (1974). Rats pressed the setup lever eight times to activate the extension of the stimulation lever, which had to be pressed only once to earn a reward. The response requirement on the setup lever can be viewed as comparable to the task of traversing the runway, and the response requirement on the stimulation lever as akin to the work required to earn a reward after entering the runway goal box. Blocks of priming and no-priming trials alternated throughout each session.

### Experiment 2: Method

#### Subjects, Electrode Implantation, Apparatus

Male Long-Evans rats (bred at Concordia University; n = 8) were pair-housed in Plexiglas^®^ cages (46 cm length x 26 cm width x 21 cm height). At the time of surgery, they were 3-4 months old and weighed at least 350 g. The housing conditions, procedures for electrode implantations, and apparatus for operant conditioning are identical to those in Experiment 1.

#### Procedures

##### Training

The procedures for training rats in Experiment 1 were similar to those in Experiment 2. Rats were each screened to determine which electrode (left or right hemisphere) and which electrical current promoted vigorous lever pressing with minimal to no disruptive stimulation-induced movements. The rats responded to currents between 210 to 440 μA. The pulse frequency of the priming and reward stimulation ranged from 184 pps to 242 pps. The rewarding stimulation consisted of a single 0.5-s train of 0.1-ms cathodal pulses. The settings determined for each rat were used throughout the experiment.

Rats were first trained to press a stimulation lever for rewarding brain stimulation on an FR8 schedule. Once the operant behavior was stable, rats were trained in the revised priming paradigm. Rats learned to press a setup lever that did not deliver reward but activated the extension of a stimulation lever located on the opposite wall of the chamber. The setup lever was initially armed on an FR1 schedule, and then the response requirement was increased gradually to FR8. A single depression of the stimulation lever earned the rat a reward.

##### Testing: Trial structure

Figure 7 (top) depicts the procedures used in the automated version of the runway paradigm (Gallistel *et al*., 1974) and thus helps draw the analogy between that procedure and the one implemented here. In Gallistel’s automated-runway paradigm, the rat is confined to the start box during the ITI, priming stimulation, and post-priming delay. The rat self-administers the priming stimulation by means of the start-box lever. Five seconds after the last prime, the start-box door drops to allow access to the alley. The rat runs to the goal box to earn a train of rewarding brain stimulation, which is triggered by a single press on the goal-box lever. Once the rat receives the reward, the goal-box lever retracts and an ITI commences during which the rat can return to the start box. At the end of the ITI the start-box lever extends to enable delivery of the priming stimulation. The first press on the start-box lever delivers one train of stimulation and raises the start-box door, thus confining the rat to the start box. The rat can then earn nine more trains of priming stimulation. Delivery of the last train is accompanied by lowering of the start-box door, thus allowing the rat to run down the alley and gain access to the goal-box lever.

The lower portion of Figure 7 depicts the revised version of our adaptation of the automated runway paradigm for use in standard operant-conditioning chambers.

- Both the setup and stimulation levers remain retracted during the 15 s ITI.
- On high-priming trials, the stimulation lever extends at the start of the priming phase, the light over the lever illuminates, the house light flashes (10 cycles of 0.5s on, 0.5s off), and a 20 s countdown begins. A single lever press triggers a train of priming stimulation and initiates retraction of the setup lever for 1 s, during which time the countdown pauses. The rat can earn up to 9 additional trains of priming before the countdown ends. Initiation of the tenth train or a countdown to zero terminates the priming phase.

○ The priming phase is identical on no priming trials except that the stimulation lever does not extend, the lever light does not illuminate, and no stimulation is available.
- There is a 5-s delay at the end of the priming phase, at which time the setup lever extends, the setup-lever light illuminates, a 10-s countdown is initiated, and the setup phase begins.
- The setup lever is armed on an FR8 schedule. If the rat completes the response requirement before the end of the 10-s countdown, the setup lever retracts, the setup lever light is extinguished, and the setup phase ends, thus triggering the start of the reward phase.

○ The stimulation lever then extends, the stimulation-lever light illuminates, and a 30-s countdown begins. A single depression of the stimulation lever triggers delivery of the rewarding stimulation and terminates the trial. If the rat fails to press the stimulation lever, the trial terminates after 30 s.
- If the rat fails to complete the response requirement during the setup phase before the end of the 10-s countdown, the setup lever retracts, the setup lever light is extinguished, and the setup phase terminates.

○ The stimulation lever does not extend at that time, nor is the stimulation-lever light illuminated. Instead, both levers remain retracted, and the trial ends 30 s later.

##### Testing: Session structure

Each test session is preceded by a warm-up session during which the rats pressed the stimulation lever for reward on an FR1 schedule for 2-3 minutes. Test sessions consist of 42 trials. The first two trials of a session are primed trials that served as learning trials and are excluded from analyses. The remaining 40 trials consist of four cycles of five primed and five unprimed trials.

#### Statistical Analyses

For each rat, data were analyzed for each priming condition (high, none). Means, medians, robust means (the location parameter obtained from application of the Tukey bisquare estimator (Hoaglin et al., 2000) and medians were calculated for each measure. Data were analyzed and graphs were plotted using MATLAB (The Mathworks, Natick, MA). Analysis and application of CIs, distributions, Cliff’s-Delta and Difference-Ratio calculation, and criteria for a priming effect are identical to those described in Experiment 1.

#### Measures of response vigor

##### Initial Speed

Initial speed is the inverse of the latency (s) to press the setup lever following its extension. A higher initial speed number indicates that a rat was quicker to initiate the first press on the setup lever. This is akin to how fast a rat exited the start box in the runway paradigm.

##### Reward Rate

Reward rate is the inverse of the total time (s) elapsed between the start of the trial and the delivery of the reward. Higher reward rates reflect more vigorous reward pursuit. This transformation is analogous to the running speed measure reported in Gallistel’s investigations of the priming effect of electrical brain stimulation (C. R. Gallistel et al., 1974).

##### Tempo

Tempo is the response rate during the setup phase. It is calculated by dividing the number of setup responses by the interval between the first and last responses. Higher tempos indicate faster reward pursuit. This is analogous to running speed in the alley alone.

#### Data transformations

##### Detrending

The key comparisons are between the data acquired in high- and no-priming conditions. Ideally, the differences between the measures in these two conditions would reflect **only** the “true” influence of priming plus random trial-by-trial noise. In reality, there is at least one additional influence. Performance can drift systematically from trial to trial, for example, decreasing as the rat tires or the effect of a drug evolves. Such a drift is seen clearly in the graphs in the upper left panel of Figure S10. Although the reward rate is systematically higher in the high-priming condition, high-priming values obtained late in the session tend to be lower than no-priming values obtained early in the session. When the data are collapsed across trials, as in the box and violin plots, the effect of the drift is confounded with the trial-to-trial noise, thus inflating the variance and the overlap.

To address the confounding effect of the drift, we estimated the linear trends and removed them, thus generating the graphs in the right column of Figure S10. Note that the high-priming data are now less dispersed (bottom right). Removing the confound boosts Cliff’s Delta from 0.70 to 0.77.

The equation for the detrending is

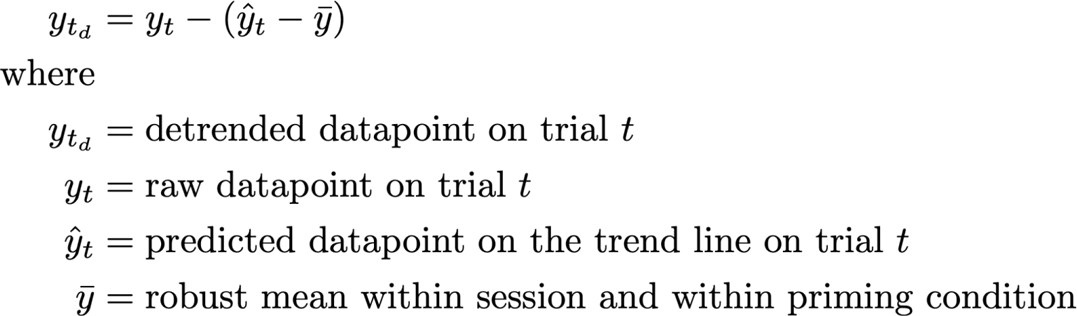

The equation tells us that each data point is moved up or down as a function of how far the trend line is from the mean of the data at the corresponding trial number.

##### Normalization

The effect-size and effect-magnitude statistics compare vigor scores from high- and no-priming conditions across trials and sessions. Whereas the detrending procedure isolates these scores from the linear component of within-session drift, a normalization procedure was implemented to isolate these scores from across-session variance in average within-session vigor (the grand robust mean across priming conditions).

The rats have “good days”, when their performance is particularly vigorous, and “not-so-good days”, when it is less so. This is illustrated in Figure S11. The black open circles represent the average vigor in each session: the grand robust mean across priming conditions. There is substantial variation. This creates a problem for the effect-size statistic. Ideally, that statistic would compare the within-session ranks of the scores under high and no priming.

However, for statistical power and to produce a single effect-size measure for the entire dataset, the scores are pooled across sessions before they are ranked. The upper left panel of Figure S12 shows that average initial speed was greater under high than low priming on all sessions.

However, the high-priming robust means in the early sessions fall below the low-priming robust means in the later sessions. When the data are pooled, the across-session variance in average vigor spuriously increases the overlap on the high- and low-priming scores.

A two-step procedure was implemented to correct for the across-session variation in the average vigor. First, each datapoint is divided by the grand robust mean across priming conditions for the session in question. The resulting values are then rescaled by multiplying them by the grand robust mean across both priming conditions and sessions.

The equation for the normalization is:

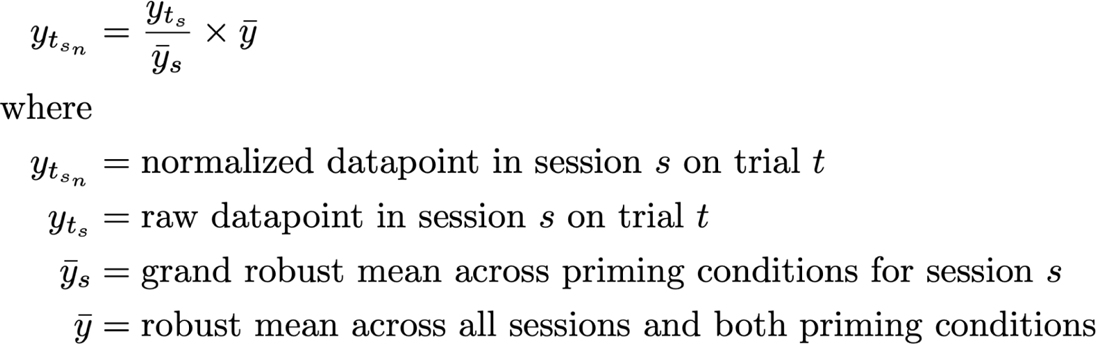

The upper right graph in Figure S12 illustrates the effect of the normalization. The across-session variation in average vigor has been removed (see the magenta diamonds in Figure S10), but the across-session variation in the priming effect (the vertical separation of the red diamonds and blue circles) remains. This latter source of variation is germane to assessing the strength of the priming effect, whereas the variation in average vigor across sessions is not.

### Experiment 2: Results

All results in this section are based on the detrended, normalized data. Figures 8 and 9 show data from rat AM46 illustrating the more vigorous responding observed in high-versus no-priming trials. Priming increased initial speed, reward rate, and tempo. Data from all rats are shown in Figures S13a-S15b.

Tables 7-9 summarize the effect of priming in all rats on the three measures of vigor: initial speed (Table 7), reward rate (Table 8), and tempo (Table 9).

- In 7 of the 8 rats, initial speed to initiate responding was both reliably and meaningfully greater on primed than on unprimed trials. The median Difference Ratio was 0.520, indicating that the median initial speed on primed trials was 26% faster than the median grand robust mean and the median initial speed on unprimed trials was 26% slower.
- Reward rate was both reliably and meaningfully greater on primed than unprimed trials in all rats. The median Difference Ratio was 0.254, indicating that the median reward rate on primed trials was 12.7% higher than the median grand robust mean and the median reward rate on unprimed trials was 12.7% lower.
- Tempo was also both reliably and meaningfully greater on primed than unprimed trials in all rats. The median Difference Ratio was 0.207, indicating that the median tempo on primed trials was 10.35% faster than the median grand robust mean and the median reward rate on unprimed trials was 10.35% slower. Thus priming boosted vigor on 23 of the 24 statistics.

**Table 7.** Effect of priming on initial speed. HP: high priming; NP: no priming; Diff: difference between high and no priming; Grand: grand robust mean across priming conditions; DiffRatio: Difference Ratio; Delta: Cliff’s Delta; DeltaCIhi: high bound of 95% CI around Cliff’s Delta; DeltaCIlo: low bound of 95% CI around Cliff’s Delta; met both: CIl around Cliff’s Delta excludes zero and Difference Ratio ≥ 0.05.

| Rat | HP | NP | Diff | Grand | DiffRatio | Delta | DeltaCIhi | DeltaCIlo | met both |
| --- | --- | --- | --- | --- | --- | --- | --- | --- | --- |
| AM44 | 0.536 | 0.554 | -0.018 | 0.540 | -0.034 | -0.005 | 0.085 | -0.091 | FALSE |
| AM45 | 0.969 | 0.771 | 0.199 | 0.878 | 0.226 | 0.400 | 0.480 | 0.320 | TRUE |
| AM46 | 1.511 | 0.844 | 0.667 | 1.181 | 0.564 | 0.894 | 0.921 | 0.865 | TRUE |
| AM47 | 1.530 | 0.676 | 0.855 | 1.097 | 0.779 | 0.837 | 0.884 | 0.784 | TRUE |
| AM48 | 1.223 | 0.617 | 0.606 | 0.923 | 0.656 | 0.938 | 0.957 | 0.916 | TRUE |
| AM50 | 1.913 | 1.287 | 0.625 | 1.554 | 0.402 | 0.761 | 0.805 | 0.714 | TRUE |
| AM55 | 1.508 | 0.930 | 0.578 | 1.217 | 0.475 | 0.892 | 0.926 | 0.859 | TRUE |
| AM58 | 1.207 | 0.552 | 0.655 | 0.891 | 0.736 | 0.795 | 0.843 | 0.744 | TRUE |
| Median | 1.365 | 0.723 | 0.615 | 1.010 | 0.520 | 0.816 | 0.864 | 0.764 |  |
| Max | 1.913 | 1.287 | 0.855 | 1.554 | 0.779 | 0.938 | 0.957 | 0.916 |  |
| Min | 0.536 | 0.552 | -0.018 | 0.540 | -0.034 | -0.005 | 0.085 | -0.091 |  |
| met criterion |  |  |  |  | 7 |  |  |  | 7 |

**Table 8.** Effect of priming on reward rate. See caption of Table 7 for details.

| Rat | HP | NP | Diff | Grand | DiffRatio | Delta | DeltaCIhi | DeltaCIlo | met both |
| --- | --- | --- | --- | --- | --- | --- | --- | --- | --- |
| AM44 | 0.185 | 0.163 | 0.023 | 0.174 | 0.131 | 0.350 | 0.429 | 0.273 | TRUE |
| AM45 | 0.272 | 0.228 | 0.044 | 0.250 | 0.178 | 0.558 | 0.627 | 0.485 | TRUE |
| AM46 | 0.328 | 0.221 | 0.107 | 0.273 | 0.391 | 0.932 | 0.952 | 0.907 | TRUE |
| AM47 | 0.307 | 0.210 | 0.097 | 0.258 | 0.378 | 0.813 | 0.864 | 0.759 | TRUE |
| AM48 | 0.249 | 0.188 | 0.061 | 0.219 | 0.279 | 0.735 | 0.775 | 0.687 | TRUE |
| AM50 | 0.347 | 0.302 | 0.044 | 0.324 | 0.137 | 0.589 | 0.651 | 0.517 | TRUE |
| AM55 | 0.306 | 0.229 | 0.076 | 0.266 | 0.287 | 0.781 | 0.831 | 0.724 | TRUE |
| AM58 | 0.223 | 0.146 | 0.077 | 0.184 | 0.417 | 0.884 | 0.914 | 0.849 | TRUE |
| Median | 0.289 | 0.215 | 0.069 | 0.254 | 0.283 | 0.758 | 0.803 | 0.706 |  |
| Max | 0.347 | 0.302 | 0.107 | 0.324 | 0.417 | 0.932 | 0.952 | 0.907 |  |
| Min | 0.185 | 0.146 | 0.023 | 0.174 | 0.131 | 0.350 | 0.429 | 0.273 |  |
| met criterion |  |  |  |  | 8 |  |  |  | 8 |

**Table 9.**
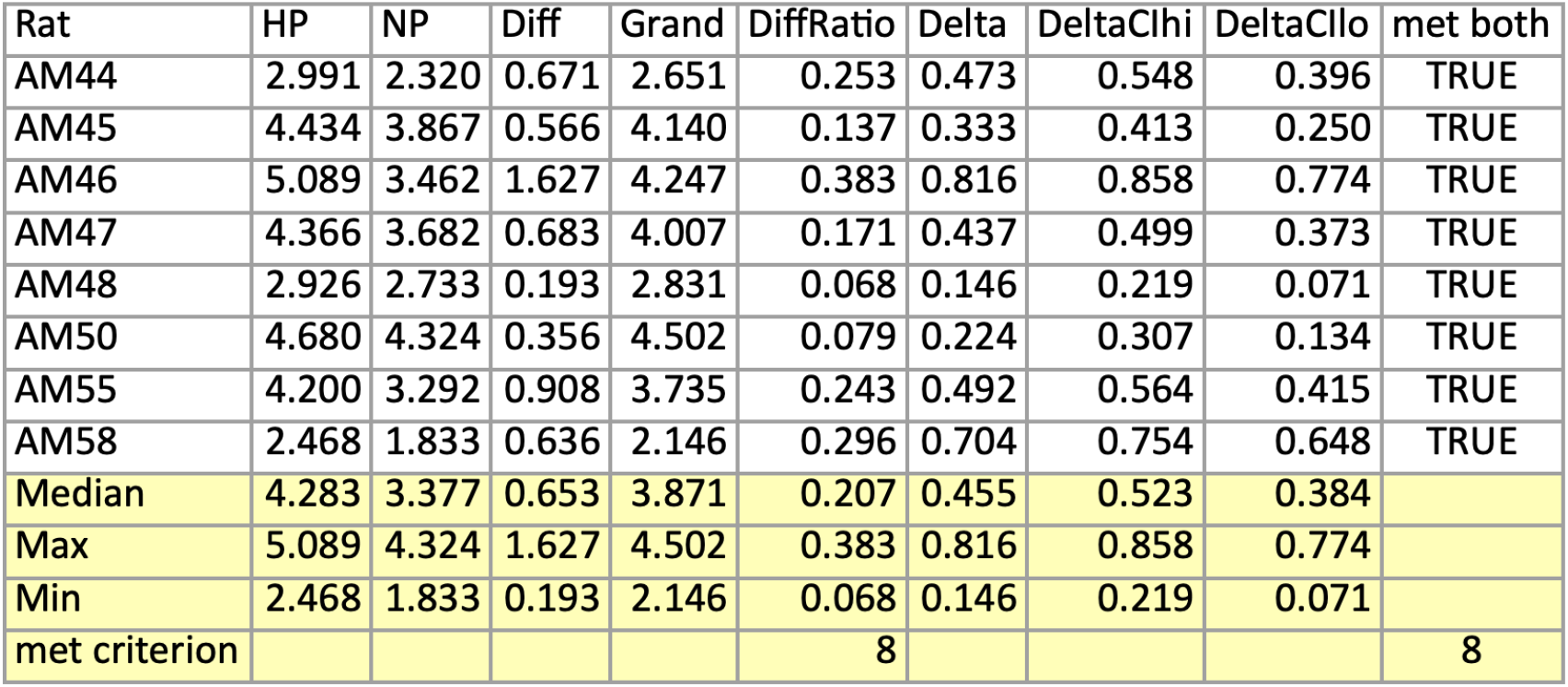
Effect of priming on tempo. See caption of Table 7 for details.

### Experiment 2: Discussion

The vigor of responding was reliably, meaningfully, and consistently greater on primed than unprimed trials. Thus, the modifications of the paradigm for measuring the priming effect and for transforming the data appear to have been beneficial. Initiation of responding during the setup phase was nearly twice as fast (across-rat medians of the initial-speed measures) on primed trials, and the reward rate was 34% higher. Effect sizes for all three measures of vigor are high (0.816, 0.758, and 0.455 for the initial-speed, reward-rate, and tempo measures, respectively, with a maximum possible value of 1.) Consistency across subjects was excellent: The effect of priming on the reward-rate and tempo measures met our criteria for reliability and meaningfulness in all eight rats, and the effect on initial speed did so in seven rats.

In the modified procedure employed in Experiment 2, most of the time required to procure the reward is consumed by satisfaction of the response requirement on the setup lever. The rat typically stands in place while it performs this component of the task. Switching between the setup and stimulation levers is mostly a matter of making a half-turn. Thus, little locomotion is entailed. MFB stimulation has been shown to elicit stepping movements (Levy & Sinnamon, 1990; Sinnamon, 1990; Sinnamon et al., 1987). This could make a non-motivational contribution to the running-speed measurements obtained in the runway. The geometry of the operant-chamber priming paradigm should remove or greatly reduce that contribution.

The priming effect measured in the standard operant-conditioning chamber closely resembles the priming effect measured in the runway. For example, work carried out in the runway paradigm established that the priming effect is immediate: Performance attains a new stable value on the very first trial on which a change in the strength or amount of priming stimulation has been introduced (Gallistel, Stellar & Bubis, 1974). In contrast, several trials are required in order for performance to fully adjust to a change in the strength of the rewarding stimulation on offer at the end of the runway (C. R. Gallistel et al., 1974). In the operant-chamber paradigm used in Experiment 2, blocks of high- and no-priming trials alternate, thus providing ample opportunity to determine whether or not the change in performance at the junction of the trial blocks is immediate. As expected, largely step-like changes in vigor are evident in the plots of vigor versus trial number in Figure 8 (bottom), S14a, and S14b.

The magnitude of priming effects in both the runway and standard operant-conditioning chamber paradigms likely depends on the value of task variables, e.g., the length of the runway and the FR requirement in the setup phase of Experiment 2. We attempted a comparison nonetheless. As explained above (Experiment 2, Method, Testing: Trial structure), the reward-rate measure obtained in the operant-chamber paradigm is analogous to the running-speed measure obtained in the runway paradigm. We derived running-speed values by visual inspection of Figures 2 and 3 in Edmonds and Gallistel’s (1974) report on their runway study and then converted those values to Difference Ratios, as described above (Experiment 1, Statistical Analyses, Difference Ratio). The Difference Ratios for the reward rates measured in Experiment 2 range from 0.131 to 0.417, overlapping those for the runway data, which range from 0.232 to 0.752. This shows that even with current task-variable settings, the priming effects measured in the operant chamber approach and sometimes exceed the magnitude of those measured in the runway. Conceivably, increasing the FR requirement in the operant-chamber paradigm would bring the values into even closer alignment.

Given the reliability and magnitude of the priming effects obtained in Experiment 2, we used the Experiment-2 paradigm in Experiment 3, in which we assess the dependence of the priming effect on dopaminergic neurotransmission.

## Experiment 3

Performance for rewarding electrical brain stimulation depends on the integrity of dopaminergic neurotransmission (Crow, 1970; Franklin, 1978; Franklin & McCoy, 1979; C. R. Gallistel & Karras, 1984; Hernandez et al., 2010, 2012; Miliaressis et al., 1986; Trujillo-Pisanty et al., 2014; Wise & Rompré, 1989). Dopaminergic neurotransmission is also implicated in motivation (Berridge & Robinson, 1998; Ikemoto & Panksepp, 1999; Robinson & Berridge, 1993). The incentive-salience theory (Berridge & Robinson, 1998; Robinson & Berridge, 1993) has been particularly influential in this regard. That theory posits separate neural systems mediate *wanting* (incentive salience) and *liking* (hedonic value) of rewards; wanting is attributed to activation of dopamine neurons.

Given the motivational character of the priming effect and the implication of dopamine signaling in motivation, one might expect that blockade of dopamine receptors would weaken the priming effect. That commonsensical prediction was not borne out when the priming effect was challenged in the runway paradigm by the administration of the dopamine D2R antagonist, pimozide (Wasserman et al., 1982). Even after receiving a very high dose, the rats showed a clear priming effect during the initial trials after the drug had taken effect. (They then ceased to respond.) Pimozide also failed to block the priming-induced preference reversal: Following delivery of priming stimulation, thirsty rats continued to choose brain stimulation in lieu of water.

We used the operant-chamber paradigm validated in Experiment 2 to revisit the question of whether the priming effect of rewarding electrical brain stimulation depends on dopaminergic neurotransmission. Given that pimozide has affinity for serotonin 7 receptors (5HT_7_Rs) we used a more specific D2R blocker instead, eticlopride (Hall et al., 1985; Martelle & Nader, 2008).

### Experiment 3: Method

#### Subjects, Apparatus

The subjects were six male Long-Evans rats from Experiment 2. The same apparatus was employed.

#### Procedures

The rats advanced from Experiment 2 to Experiment 3 without further training. Session structure was the same as in Experiment 2, and the same statistical procedures were employed.

#### Drug treatment

Eticlopride (0.1 or 0.05 mg/kg, Sigma, St. Louis, MO) was dissolved in physiological saline (0.9%). These doses were based on data from a previous pilot study (data not shown) and from Lazenka et al. (2016). Vehicle (0.9% physiological saline) or one of the two doses of eticlopride was administered intraperitoneally (i.p) 30 minutes prior to testing. Tests were conducted in three-day cycles consisting of a vehicle day, drug day, and washout day. We aimed to conduct ten of these 3-day cycles for each drug dose. However, there were instances where rats did not perform at all at the highest dose (0.1 mg/kg) of eticlopride and thus testing at that dose was discontinued. Tests were first conducted at the highest dose of eticlopride (0.1 mg/kg). Six rats underwent testing with the highest dose of eticlopride. However, only three rats (AM47, AM55, AM58) successfully completed testing at that dose, which induced catatonia in the other rats (AM46, AM48, AM50) following repeated administration. Those latter three rats underwent testing at the lowest dose of eticlopride. The rats that were tested at the highest dose of eticlopride were not tested with the lowest dose of eticlopride due to attrition (e.g., health, electrode stability, etc.).

### Experiment 3: Results

The priming effect was seen in both the vehicle and drug conditions, even at the highest dose of eticlopride (0.1 mg/kg). This is illustrated in Figures 10-12 in the case of rat AM47. Only four testing sessions were completed by this rat under the effects of the 0.1 mg/kg dose, but the variance is low, and a clear effect of priming stimulation is evident. In the remaining drug sessions, rat AM47 initiated responding on the setup lever on a total of only 35 of the 240 trials (6 sessions × 40 trials/session) and only harvested a reward on 17 of these trials. In contrast, all 10 test sessions were completed successfully following vehicle administration. Although priming continued to boost performance substantially under the influence of 0.1 mg/kg of eticlopride in rat AM47, the initiation of responding in the setup phase (Figure 10) and rate of responding on the setup and reward phases (Figure 3) were slowed. However, once responding had been initiated, response tempo was equal or greater than that observed in the vehicle condition (Figure 12).

Results from the other two rats that completed at least four sessions under the effects of the 0.1 mg/kg dose (AM55, AM58) are shown in Figures S1-S6. Rat AM55 completed all 10 vehicle sessions and five drug sessions, whereas rat AM58 completed nine of 10 vehicle sessions and five drug sessions. Figures 10-12 and S1-S6 show only the results obtained on the completed sessions.

Tables 10, 12, and 14 compile the values of the Difference Ratio and Cliff’s Delta on the three measures of vigor in the vehicle condition for all three rats that received the 0.1 mg/kg dose of eticlopride. The results from the drug condition are shown in Tables 11, 13, and 15. The reward-rate and tempo results meet our criteria for a meaningful (Difference Ratio ≧ 0.05) and reliable (CI around Cliff’s Delta excludes 0) effect in both the vehicle and drug conditions in all three rats. All three rats showed a meaningful and reliable priming effect in the vehicle condition on the initial-speed measure, whereas in the eticlopride condition, rats AM47 and AM55 met the criteria for a meaningful and reliable priming effect but AM58 did not. Both the Difference Ratio and Cliff’s Delta scores for all three vigor measures were higher in the eticlopride data from rat AM47 than in the vehicle data, and the CIs around Cliff’s Delta for the drug condition do not overlap those for the vehicle condition in the cases of the reward-rate and tempo measures. In other words, the priming effect in this rat was generally stronger under the influence of eticlopride than in the vehicle condition.

**Table 10.** Initial speed to initiate responding in the setup phase in rats AM47, AM55, and AM58 following administration of the vehicle (top) or 0.1 mg/kg eticlopride (bottom).

Initial Speed: Vehicle
| Rat | HP | NP | Diff | Grand | DiffRatio | Delta | DeltaCIhi | DeltaCIlo | met both |
| --- | --- | --- | --- | --- | --- | --- | --- | --- | --- |
| AM47 | 1.565 | 0.881 | 0.684 | 1.203 | 0.569 | 0.779 | 0.849 | 0.708 | TRUE |
| AM55 | 1.233 | 0.711 | 0.522 | 0.969 | 0.539 | 0.665 | 0.744 | 0.578 | TRUE |
| AM58 | 1.228 | 0.981 | 0.248 | 1.108 | 0.224 | 0.313 | 0.432 | 0.197 | TRUE |
| Median | 1.233 | 0.881 | 0.522 | 1.108 | 0.539 | 0.665 | 0.744 | 0.578 |  |
| Max | 1.565 | 0.981 | 0.684 | 1.203 | 0.569 | 0.779 | 0.849 | 0.708 |  |
| Min | 1.228 | 0.711 | 0.248 | 0.969 | 0.224 | 0.313 | 0.432 | 0.197 |  |
| met criterion |  |  |  |  | 3 |  |  | 3 | 3 |

**Table 11.** Initial speed to initiate responding in the setup phase in rats AM47, AM55, and AM58 following administration of the vehicle (top) or 0.1 mg/kg eticlopride (bottom).

Initial Speed: 0.1 mg/kg Eticlopride
| Rat | HP | NP | Diff | Grand | DiffRatio | Delta | DeltaCIhi | DeltaCIlo | met both |
| --- | --- | --- | --- | --- | --- | --- | --- | --- | --- |
| AM47 | 1.101 | 0.488 | 0.613 | 0.788 | 0.778 | 0.842 | 0.920 | 0.738 | TRUE |
| AM55 | 0.648 | 0.555 | 0.092 | 0.603 | 0.153 | 0.140 | 0.263 | 0.014 | TRUE |
| AM58 | 0.692 | 0.641 | 0.051 | 0.653 | 0.078 | 0.169 | 0.330 | -0.009 | FALSE |
| Median | 0.692 | 0.555 | 0.092 | 0.653 | 0.153 | 0.169 | 0.330 | 0.014 |  |
| Max | 1.101 | 0.641 | 0.613 | 0.788 | 0.778 | 0.842 | 0.920 | 0.738 |  |
| Min | 0.648 | 0.488 | 0.051 | 0.603 | 0.078 | 0.140 | 0.263 | -0.009 |  |
| met criterion |  |  |  |  | 3 |  |  | 2 | 2 |

**Table 12.** Reward rate in rats AM47, AM55, and AM58 following administration of the vehicle (top) or 0.1 mg/kg eticlopride (bottom).

Reward Rate: Vehicle
| Rat | HP | NP | Diff | Grand | DiffRatio | Delta | DeltaCIhi | DeltaCilo | met both |
| --- | --- | --- | --- | --- | --- | --- | --- | --- | --- |
| AM47 | 0.331 | 0.233 | 0.098 | 0.278 | 0.351 | 0.758 | 0.836 | 0.671 | TRUE |
| AM55 | 0.250 | 0.189 | 0.060 | 0.218 | 0.277 | 0.704 | 0.779 | 0.618 | TRUE |
| AM58 | 0.222 | 0.195 | 0.027 | 0.210 | 0.130 | 0.387 | 0.500 | 0.272 | TRUE |
| Median | 0.250 | 0.195 | 0.060 | 0.218 | 0.277 | 0.704 | 0.779 | 0.618 |  |
| Max | 0.331 | 0.233 | 0.098 | 0.278 | 0.351 | 0.758 | 0.836 | 0.671 |  |
| Min | 0.222 | 0.189 | 0.027 | 0.210 | 0.130 | 0.387 | 0.500 | 0.272 |  |
| met criterion |  |  |  |  | 3 |  |  | 3 | 3 |

**Table 13.** Reward rate in rats AM47, AM55, and AM58 following administration of the vehicle (top) or 0.1 mg/kg eticlopride (bottom).

Reward Rate: 0.1 mg/kg Eticlopride
| Rat | HP | NP | Diff | Grand | DiffRatio | Delta | DeltaCIhi | DeltaCilo | met both |
| --- | --- | --- | --- | --- | --- | --- | --- | --- | --- |
| AM47 | 0.291 | 0.181 | 0.110 | 0.236 | 0.467 | 0.940 | 0.986 | 0.871 | TRUE |
| AM55 | 0.206 | 0.183 | 0.024 | 0.193 | 0.123 | 0.284 | 0.396 | 0.162 | TRUE |
| AM58 | 0.184 | 0.148 | 0.036 | 0.170 | 0.213 | 0.371 | 0.528 | 0.190 | TRUE |
| Median | 0.206 | 0.181 | 0.036 | 0.193 | 0.213 | 0.371 | 0.528 | 0.190 |  |
| Max | 0.291 | 0.183 | 0.110 | 0.236 | 0.467 | 0.940 | 0.986 | 0.871 |  |
| Min | 0.184 | 0.148 | 0.024 | 0.170 | 0.123 | 0.284 | 0.396 | 0.162 |  |
| met criterion |  |  |  |  | 3 |  |  | 3 | 3 |

**Table 14.** Response tempo during the setup phase in rats AM47, AM55, and AM58 following administration of the vehicle (top) or 0.1 mg/kg eticlopride (bottom).

Tempo: Vehicle
| Rat | HP | NP | Diff | Grand | DiffRatio | Delta | DeltaCIhi | DeltaCilo | met both |
| --- | --- | --- | --- | --- | --- | --- | --- | --- | --- |
| AM47 | 4.552 | 3.551 | 1.000 | 4.006 | 0.250 | 0.495 | 0.593 | 0.394 | TRUE |
| AM55 | 3.446 | 2.807 | 0.639 | 3.116 | 0.205 | 0.411 | 0.524 | 0.296 | TRUE |
| AM58 | 2.622 | 2.371 | 0.251 | 2.507 | 0.100 | 0.308 | 0.421 | 0.178 | TRUE |
| Median | 3.446 | 2.807 | 0.639 | 3.116 | 0.205 | 0.411 | 0.524 | 0.296 |  |
| Max | 4.552 | 3.551 | 1.000 | 4.006 | 0.250 | 0.495 | 0.593 | 0.394 |  |
| Min | 2.622 | 2.371 | 0.251 | 2.507 | 0.100 | 0.308 | 0.421 | 0.178 |  |
| met criterion |  |  |  |  | 3 |  |  | 3 | 3 |

**Table 15.** Response tempo during the setup phase in rats AM47, AM55, and AM58 following administration of the vehicle (top) or 0.1 mg/kg eticlopride (bottom).

Tempo: 0.1 mg/kg Eticlopride
| Rat | HP | NP | Diff | Grand | DiffRatio | Delta | DeltaCIhi | DeltaCilo | met both |
| --- | --- | --- | --- | --- | --- | --- | --- | --- | --- |
| AM47 | 5.304 | 3.938 | 1.366 | 4.659 | 0.293 | 0.722 | 0.827 | 0.609 | TRUE |
| AM55 | 3.507 | 3.023 | 0.484 | 3.251 | 0.149 | 0.345 | 0.452 | 0.239 | TRUE |
| AM58 | 2.501 | 1.832 | 0.669 | 2.181 | 0.307 | 0.463 | 0.597 | 0.314 | TRUE |
| Median | 3.507 | 3.023 | 0.669 | 3.251 | 0.293 | 0.463 | 0.597 | 0.314 |  |
| Max | 5.304 | 3.938 | 1.366 | 4.659 | 0.307 | 0.722 | 0.827 | 0.609 |  |
| Min | 2.501 | 1.832 | 0.484 | 2.181 | 0.149 | 0.345 | 0.452 | 0.239 |  |
| met criterion |  |  |  |  | 3 |  |  | 3 | 3 |

Figures 13-15 illustrate the effect of the lower dose of eticlopride (0.05 mg/kg) on the three measures of vigor in rat AM46, one of the three rats that were unable to complete at least four sessions under the influence of the higher dose (0.1 mg/kg). This rat completed 9 of ten test sessions in both the vehicle and drug conditions, and a clear priming effect is seen in both conditions.

The results from the other two rats tested at this dose (AM48, AM50) are shown in Figures S7-S12. Rat AM48 completed 9 of ten test sessions in the vehicle condition and four of 10 sessions in the drug condition, whereas rat AM50 completed all ten test sessions in the vehicle condition and nine of 10 sessions in the drug condition.

Tables 16, 18, and 20 compile the values of the Difference Ratio and Cliff’s Delta on the three measures of vigor in the vehicle condition for all three rats that received the 0.0.05 mg/kg dose of eticlopride. The results from the drug condition are shown in Tables 17, 19, and 21. The initial-speed and reward-rate results meet our criteria for a meaningful (Difference Ratio ≧ 0.05) and reliable (CI around Cliff’s Delta excludes 0) effect in both the vehicle and drug conditions in all three rats. This was also the case for the tempo measure in rats AM46 and AM50. Rat AM48 did not show a priming effect on the tempo measure in either the vehicle or the drug condition.

**Table 16.**
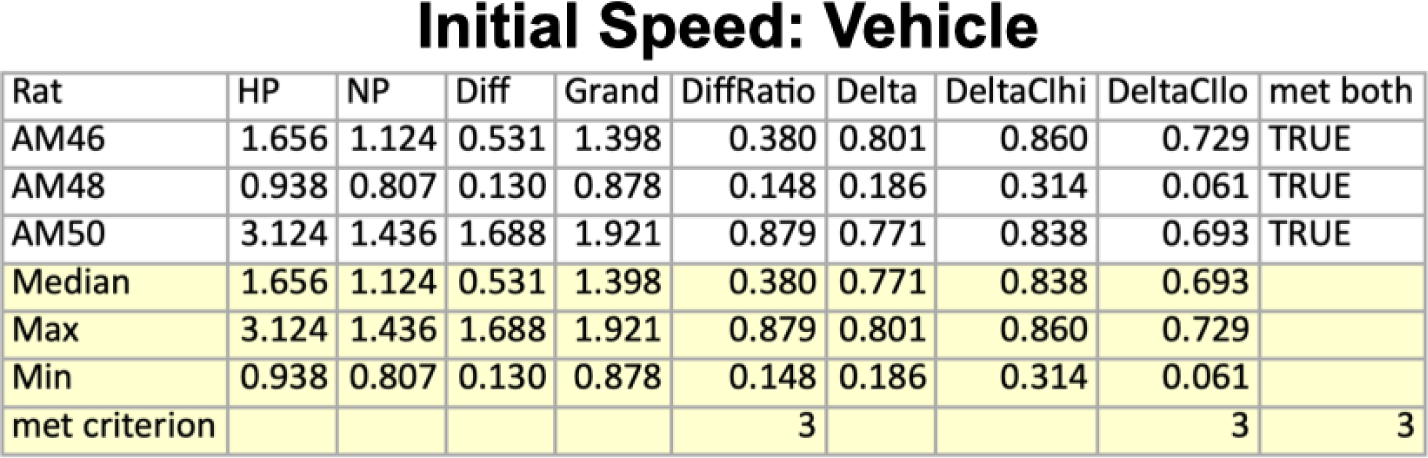
Initial speed to initiate responding in the setup phase in rats AM46, AM48, and AM50 following administration of the vehicle (top) or 0.05 mg/kg eticlopride (bottom).

**Table 17.** Initial speed to initiate responding in the setup phase in rats AM46, AM48, and AM50 following administration of the vehicle (top) or 0.05 mg/kg eticlopride (bottom).

Initial Speed: 0.05 mg/kg Eticlopride
| Rat | HP | NP | Diff | Grand | DiffRatio | Delta | DeltaClhi | DeltaCllo | met both |
| --- | --- | --- | --- | --- | --- | --- | --- | --- | --- |
| AM46 | 1.540 | 1.010 | 0.531 | 1.291 | 0.411 | 0.740 | 0.818 | 0.655 | TRUE |
| AM48 | 0.975 | 0.804 | 0.171 | 0.890 | 0.192 | 0.283 | 0.445 | 0.115 | TRUE |
| AM50 | 2.288 | 1.298 | 0.991 | 1.709 | 0.580 | 0.710 | 0.779 | 0.621 | TRUE |
| Median | 1.540 | 1.010 | 0.531 | 1.291 | 0.411 | 0.710 | 0.779 | 0.621 |  |
| Max | 2.288 | 1.298 | 0.991 | 1.709 | 0.580 | 0.740 | 0.818 | 0.655 |  |
| Min | 0.975 | 0.804 | 0.171 | 0.890 | 0.192 | 0.283 | 0.445 | 0.115 |  |
| met criterion |  |  |  |  | 3 |  |  | 3 | 3 |

**Table 18.**
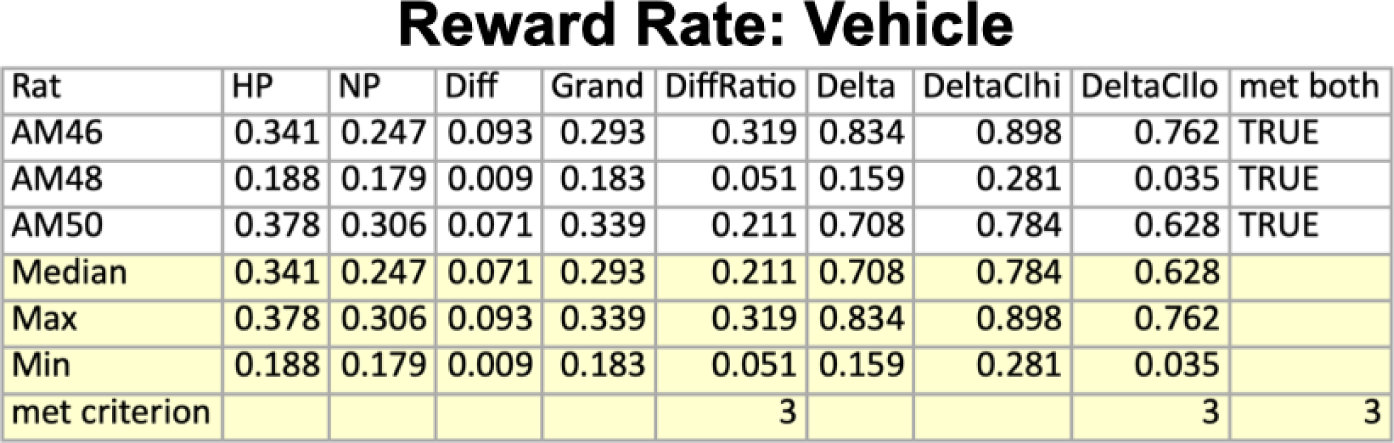
Reward rate in rats AM46, AM48, and AM50 following administration of the vehicle (top) or 0.05 mg/kg eticlopride (bottom).

Reward Rate: Vehicle
| Rat | HP | NP | Diff | Grand | DiffRatio | Delta | DeltaClhi | DeltaCllo | met both |
| --- | --- | --- | --- | --- | --- | --- | --- | --- | --- |
| AM46 | 0.341 | 0.247 | 0.093 | 0.293 | 0.319 | 0.834 | 0.898 | 0.762 | TRUE |
| AM48 | 0.188 | 0.179 | 0.009 | 0.183 | 0.051 | 0.159 | 0.281 | 0.035 | TRUE |
| AM50 | 0.378 | 0.306 | 0.071 | 0.339 | 0.211 | 0.708 | 0.784 | 0.628 | TRUE |
| Median | 0.341 | 0.247 | 0.071 | 0.293 | 0.211 | 0.708 | 0.784 | 0.628 |  |
| Max | 0.378 | 0.306 | 0.093 | 0.339 | 0.319 | 0.834 | 0.898 | 0.762 |  |
| Min | 0.188 | 0.179 | 0.009 | 0.183 | 0.051 | 0.159 | 0.281 | 0.035 |  |
| met criterion |  |  |  |  | 3 |  |  | 3 | 3 |

**Table 19.** Reward rate in rats AM46, AM48, and AM50 following administration of the vehicle (top) or 0.05 mg/kg eticlopride (bottom).

Reward Rate: 0.05 mg/kg Eticlopride
| Rat | HP | NP | Diff | Grand | DiffRatio | Delta | DeltaClhi | DeltaCllo | met both |
| --- | --- | --- | --- | --- | --- | --- | --- | --- | --- |
| AM46 | 0.325 | 0.236 | 0.089 | 0.281 | 0.316 | 0.889 | 0.933 | 0.838 | TRUE |
| AM48 | 0.191 | 0.174 | 0.016 | 0.181 | 0.090 | 0.354 | 0.513 | 0.189 | TRUE |
| AM50 | 0.371 | 0.287 | 0.083 | 0.327 | 0.255 | 0.821 | 0.885 | 0.755 | TRUE |
| Median | 0.325 | 0.236 | 0.083 | 0.281 | 0.255 | 0.821 | 0.885 | 0.755 |  |
| Max | 0.371 | 0.287 | 0.089 | 0.327 | 0.316 | 0.889 | 0.933 | 0.838 |  |
| Min | 0.191 | 0.174 | 0.016 | 0.181 | 0.090 | 0.354 | 0.513 | 0.189 |  |
| met criterion |  |  |  |  | 3 |  |  | 3 | 3 |

**Table 20.** Response tempo during the setup phase in rats AM46, AM48, and AM50 following administration of the vehicle (top) or 0.05 mg/kg eticlopride (bottom).

Tempo: Vehicle
| Rat | HP | NP | Diff | Grand | DiffRatio | Delta | DeltaClhi | DeltaCllo | met both |
| --- | --- | --- | --- | --- | --- | --- | --- | --- | --- |
| AM46 | 5.173 | 3.785 | 1.387 | 4.455 | 0.311 | 0.707 | 0.778 | 0.625 | TRUE |
| AM48 | 2.165 | 2.171 | -0.006 | 2.169 | -0.003 | 0.008 | 0.143 | -0.112 | FALSE |
| AM50 | 4.695 | 4.205 | 0.490 | 4.432 | 0.111 | 0.295 | 0.402 | 0.180 | TRUE |
| Median | 4.695 | 3.785 | 0.490 | 4.432 | 0.111 | 0.295 | 0.402 | 0.180 |  |
| Max | 5.173 | 4.205 | 1.387 | 4.455 | 0.311 | 0.707 | 0.778 | 0.625 |  |
| Min | 2.165 | 2.171 | -0.006 | 2.169 | -0.003 | 0.008 | 0.143 | -0.112 |  |
| met criterion |  |  |  |  | 2 |  |  | 2 | 2 |

**Table 21.** Response tempo during the setup phase in rats AM46, AM48, and AM50 following administration of the vehicle (top) or 0.05 mg/kg eticlopride (bottom).

Tempo: 0.05 mg/kg Eticlopride
| Rat | HP | NP | Diff | Grand | DiffRatio | Delta | DeltaClhi | DeltaCllo | met both |
| --- | --- | --- | --- | --- | --- | --- | --- | --- | --- |
| AM46 | 5.072 | 3.714 | 1.359 | 4.393 | 0.309 | 0.753 | 0.819 | 0.673 | TRUE |
| AM48 | 2.197 | 2.099 | 0.098 | 2.146 | 0.045 | 0.135 | 0.317 | -0.045 | FALSE |
| AM50 | 5.099 | 4.020 | 1.079 | 4.569 | 0.236 | 0.589 | 0.682 | 0.490 | TRUE |
| Median | 5.072 | 3.714 | 1.079 | 4.393 | 0.236 | 0.589 | 0.682 | 0.490 |  |
| Max | 5.099 | 4.020 | 1.359 | 4.569 | 0.309 | 0.753 | 0.819 | 0.673 |  |
| Min | 2.197 | 2.099 | 0.098 | 2.146 | 0.045 | 0.135 | 0.317 | -0.045 |  |
| met criterion |  |  |  |  | 2 |  |  | 2 | 2 |

## Experiment 3: Discussion

Wasserman, Gomita and Gallistel (1982) used the runway paradigm to determine whether the priming effect of rewarding electrical brain stimulation persists under the influence of moderate to very high doses of pimozide. They found that the drug failed to block the energizing effect of the priming stimulation: On the first few trials following the onset of drug action, the rats traversed the runway more quickly after having received priming stimulation prior to being placed in the start box than on trials without priming stimulation. Wasserman et al. (1982) also showed that the directing effect of the priming stimulation persisted under the influence of the drug: Thirsty rats chose brain stimulation in preference to water following priming but preferred water in the absence of priming.

We assessed the role of D2Rs in the priming effect in a manner complementary to the methods used by Wasserman et al. (1982) Instead of administering pimozide, which blocks serotonin 5HT_7_Rs in addition to blocking D2Rs, we administered eticlopride, a more specific D2R antagonist (Hall et al., 1985; Martelle & Nader, 2008). We used our operant-conditioning chamber paradigm (Experiment 2) to assess the priming effect in lieu of the runway paradigm. Within each session, blocks of high-priming and no-priming trials alternated, thus providing multiple measurements of the priming effect. Our results are entirely consistent with the findings of Wasserman et al. (1982).

All three rats tested at the 0.1 mg/kg dose of eticlopride and all three rats tested at the 0.05 mg/kg dose showed reliable and meaningful priming effects under the influence of the drug on two of the three measures of vigor; two rats in each dose group also showed reliable and meaningful priming effects under the influence of the drug on the third measure of vigor. In five of six rats, the robust grand mean of the initial-speed and reward-rate measures was lower under the influence of the drug than in the vehicle condition. This shows that both doses of the drug were behaviorally effective but nonetheless failed to block the priming effect. Similar findings have been demonstrated with a natural reward: D2R antagonism does not block the priming effect of palatable foods (Evangelista et al., 2019).

Below, we discuss the implications of these findings for contemporary models of brain reward circuitry and the role of dopaminergic neurotransmission within them.

## General Discussion

The results of the three experiments provide new findings about the priming effect, its measurement and neuropharmacological basis while also confirming several prior results. The experiments introduce, refine, and apply a method for measuring the priming effect of rewarding electrical brain stimulation in standard operant-conditioning chambers. These are substantially more compact than the runway used in much prior work on the priming phenomenon and are already installed in most behavioral neuroscience laboratories. We anticipate that advantages in compactness and availability will promote further priming studies by facilitating the concurrent testing of multiple subjects.

Performance in the runway implies that the priming effect of rewarding electrical brain stimulation depends on the expected value of the stimulation available in the goal box (C. R. Gallistel et al., 1974). When the goal box stimulation is strongly rewarding and priming stimulation has been delivered in the start box, the rat typically shows great excitement and may attempt to vault over the barrier blocking access to the runway. In contrast, once the goal-box stimulation has been turned off, and the rat has had several trials to ascertain this, it typically behaves apathetically in the start box and is in no hurry to leave it once the barrier has been removed. Thus, the priming effect in the runway depends on the rat’s expectation of goal-box reward. The results of Experiment 1 confirm that is also the case in the standard operant-conditioning chamber paradigm. The priming effect measured in that paradigm declined as either the strength of the response-contingent reward decreased or the cost of that reward increased.

The priming effect in the runway paradigm accumulates during delivery of a series of priming trains (C. R. Gallistel, 1969; Sax & Gallistel, 1991). Similarly, delivery of 10 priming trains in the operant-chamber paradigm (Experiment 1) produced more vigorous subsequent responding than delivery of two priming trains.

### Neural circuitry underlying priming and reward

The account of intracranial self-stimulation that Deutsch derived from his theory of reward and reinforcement (Deutsch, 1960; Deutsch & Howarth, 1963) predicts that two different neural systems are directly activated by rewarding brain stimulation, one responsible for priming and the second for reward. Subsequent psychophysical measurements of strength-duration functions (Matthews, 1977) and recovery from refractoriness (Gallistel, Shizgal & Yeomans, 1981) fail to provide support for that hypothesis. The simplest explanation of this failure is that the very same directly activated neurons give rise to both the priming and reward effects.

Sax and Gallistel (1991) used psychophysical methods to infer temporal integration characteristics of the neural substrates for priming and reward. These characteristics (e.g., reciprocity between the current and pulse frequency required to produce an effect of a given strength) almost certainly arise in postsynaptic targets of the directly stimulated substrate. Sax and Gallistel found that the resulting trade-off functions differ only in scale and not in shape. The similarity in shape is consistent with a common neural substrate for priming and reward encompassing at least the first two stages of the circuitry. Sax and Gallistel interpreted the scale difference in terms of the hypothesis that it is “the experience of several recent large rewards that excites the transient motivational state called the priming effect of the stimulation, the state that makes the animal avidly pursue more brain stimulation.” Stated otherwise, the quantity of stimulation required to produce a behaviorally measurable priming effect exceeds the quantity required to produce a behaviorally measurable reward effect.

If common neural circuitry subserves both the priming and rewarding effects, how is one to explain the fact that the priming effect is transient and the reward effect is enduring? The neural mechanisms underlying the two effects must ultimately diverge. Sax and Gallistel’s account places this divergence beyond the mono-synaptically activated stage. They propose that after the divergence, the branch subserving the reward effect translates the transient stimulation-induced activation into an engram, whereas the branch subserving the priming effect provides a potentiating input to the neural circuitry subserving the goal-seeking actions. The results of Experiment 3 and those of Wasserman et al. (1982) bear directly on the question of where, in the diverging pathway, the midbrain dopamine neurons intervene.

### The role of dopaminergic neurotransmission

Extensive psychophysical data argue that the directly activated stage of the circuitry responsible for self-stimulation of the MFB consists largely of neurons with myelinated, non-dopaminergic axons that are continuous between the LH and ventral tegmental area (Bielajew et al., 1982; C. Gallistel et al., 1981; Shizgal et al., 1980). The behaviorally consequential direction of conduction in at least some of these fibers is rostro-caudal (Bielajew & Shizgal, 1986). The reason that these non-dopaminergic, myelinated axons predominate in the directly stimulated stage is that they are much more highly excitable to extracellular currents than the fine, unmyelinated, dopaminergic axons interspersed among them (Anderson et al., 1996; Yeomans et al., 1988).

To reconcile the extensive data implicating dopaminergic neurotransmission in MFB self-stimulation with the evidence that the directly activated neurons are at least largely non-dopaminergic, a “series-circuit” model was proposed (Bielajew & Shizgal, 1986; Shizgal et al., 1980; Wise, 1980). On that view, the midbrain dopamine neurons relay the activation of the directly stimulated fibers to the later stages of the neural circuitry, the stages that subserve the reward-seeking actions and engram formation. The combination of the series-circuit model and the evidence suggesting that both priming and reward arise from excitation of the same directly stimulated neurons thus predict that blockade of dopaminergic neurotransmission will block both the priming and reward effects. However, Wasserman et al. (1982) showed that this is not the case: The priming effect survives blockade of dopamine D2Rs by pimozide. We show in Experiment 3 that this is also the case with the more specific action of eticlopride on these receptors. An alternative to the series-circuit model is required to explain these findings. The convergence model proposed by Trujillo-Pisanty and Shizgal (2020) fits the bill.

The convergence model holds that signals arising in the directly activated myelinated fibers composing the MFB substrate reach the final common path for the rewarding effect in parallel to signals arising from activation of midbrain dopamine neurons. That view is readily combined with the hypothesis that the initial stages of the neural circuitry subserving the priming and rewarding effects are the same (Sax & Gallistel, 1991) and with the data demonstrating the resistance of the priming effect to perturbation of dopaminergic neurotransmission. The key concept that enables this combination is the bifurcation Sax and Gallistel propose in the causal pathways for the priming and rewarding effects. They argue that this bifurcation occurs beyond both the directly stimulated stage and the subsequent stage(s) underlying spatio-temporal integration. Prior to the bifurcation point, the electrically induced activity has been integrated to yield a unidimensional signal. Beyond the bifurcation, one branch accumulates the unidimensional signals produced by multiple stimulation trains, provided that they are separated by 1-2 s and provided that their strength exceeds the threshold for energizing behavioral performance (Sax & Gallistel, 1991). Following the termination of the series of trains, the accumulated excitation decays over tens of seconds. That is the excitation that is expressed as the priming effect. The second branch generates the rewarding effect under dopaminergic modulation, translating the transient stimulation-induced activation into an enduring entry into multiple mnemonic representations, thus capturing, at the very least, the magnitude of the activation as well as the time and place at which it was triggered (C. R. Gallistel, 2020).

Work by Shizgal and colleagues (Hernandez et al., 2010, 2012; Trujillo-Pisanty et al., 2014) constrains the manner in which the dopaminergic modulation may occur. The option that meshes most cleanly with the Sax and Gallistel hypothesis is that dopaminergic drive scales the reward-intensity value encoded in memory. On that view, this is why the rats that had medium-high doses of pimozide ceased to run down the alley in the Wasserman et al. (1982) study and why three of the rats in the present study ceased to work in the operant-chamber priming paradigm on most sessions following administration of the higher dose of eticlopride. Under the influence of the lower dose of eticlopride in the present study or a low dose of pimozide (0.1 mg/kg) in the Trujillo-Pisanty et al. study (2014), performance could be sustained, but the latter study shows that the cost of the reward had to be reduced to compensate for the effect of the drug.

### Priming, dopamine and incentive motivation

The priming effect of rewarding electrical brain stimulation is a special case of incentive motivation (Bindra, 1968, 1969), one in which the motivation-enhancing effect of exposure to a recent reward arises from direct stimulation of central neurons. The results of this experiment, coupled with the prior findings of Wasserman et al. (1982) show that this effect is resistant to blockade of dopamine D2Rs. That finding generalizes beyond the effects of MFB stimulation. We have shown that the priming effect produced by consumption of highly palatable food also resists blockade of dopamine D2Rs as well as blockade of dopamine D1-like receptors (Evangelista et al., 2019). How are such findings to be reconciled with the abundant data implicating dopaminergic neurotransmission in incentive motivation (e.g., (Berridge & Robinson, 2003; Robinson & Berridge, 1993))? The key is to keep firmly in mind the convergent causation of so many behavioral phenomena and to resist the temptation to assume covertly that the neural substrate for any particular causal factor is the final common path for all the others.

The convergence model of brain stimulation reward in no way disputes the ability of dopaminergic activation to produce powerfully rewarding effects. Instead, it merely asserts that there are other means of generating these effects that operate in parallel to dopaminergic neurons. Similarly, we argue here that non-dopaminergic neurons can produce incentive-motivational effects in parallel to the circuitry underlying those generated by dopaminergic activation. On that view, the final common path for incentive motivational effects lies beyond the point of convergence between inputs that relay signals of dopaminergic and non-dopaminergic origin. We know a great deal about the dopaminergic inputs. The results reported here and in prior work on the priming effects of rewarding electrical brain stimulation and palatable food (Evangelista et al., 2019) argue that it would be worthwhile to learn much more about the significance, function, and neural instantiation of the non-dopaminergic inputs.

## Data accessibility statement

Data are available at Open Science Foundation website: https://osf.io/xe968/

## Notes

### Competing Interest Statement

The authors have declared no competing interest.

### Summary of Updates

This version updates and merges two prior manuscripts, BIORXIV/2022/512941 and, BIORXIV/2022/512944, as well as their supporting-information files.

