## supplemental figures for "The priming effect of rewarding brain stimulation in rats depends on both the cost and strength of reward but survives blockade of D2-like dopamine receptors"

Peter Shizgal*

Concordia University

### Supporting information

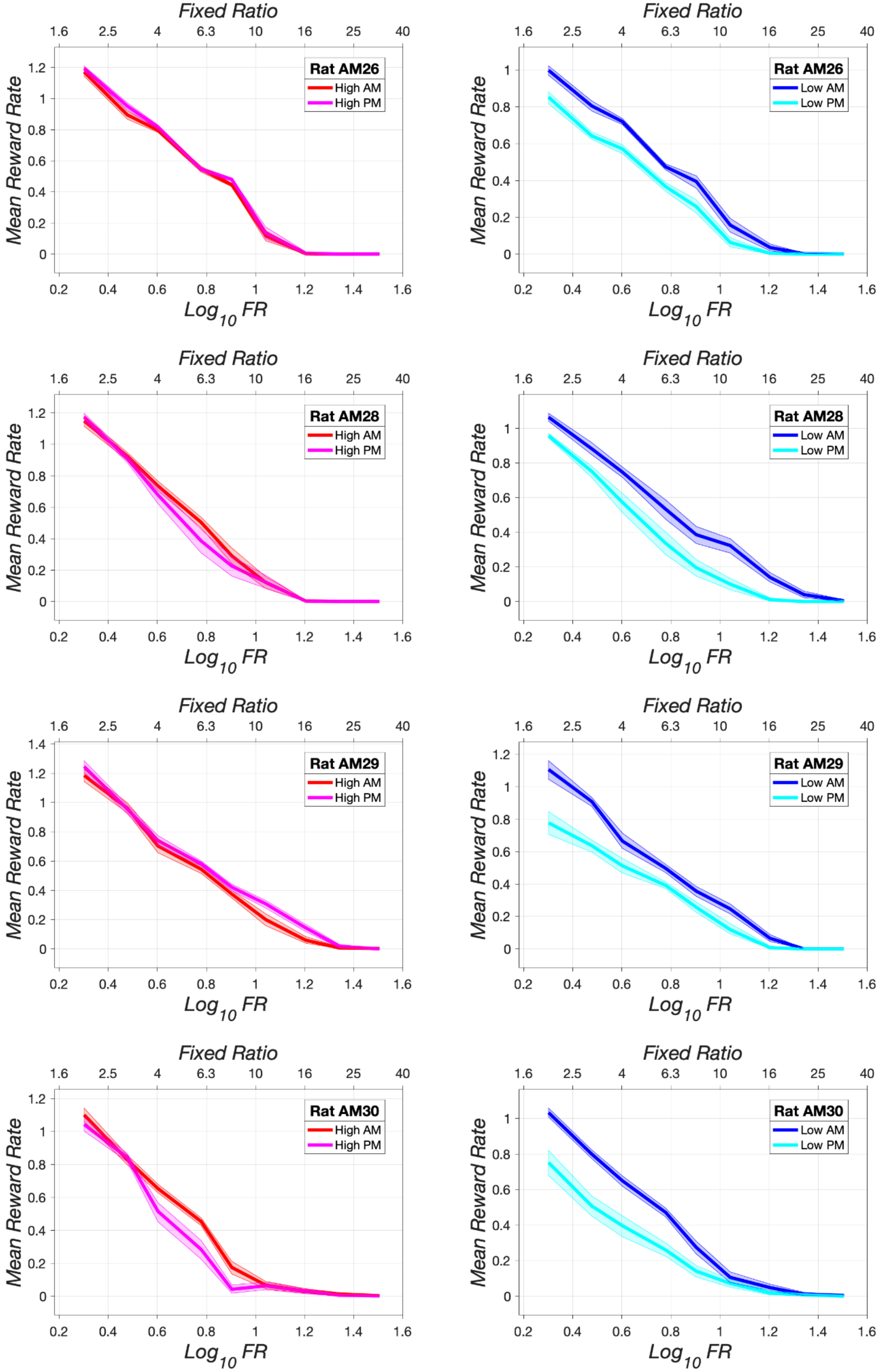
Fixed Ratio
1.6
2.5
4
6.3
10
16
25
40
Mean Reward Rate
Log10 FR
Rat AM26
High AM
High PM
Low AM
Low PM
Rat AM28
Rat AM29
Rat AM30

Priming as a function of reward cost, reward strength, and D2-like blockade: Supporting information

***Figure S1:*** Mean reward rate as a function of reward cost (fixed-ratio requirement) in morning (AM) and afternoon (PM) test sessions under low(right) and high(left)  priming for rats AM26, AM28, AM29, and AM30. The error band shows the 95% confidence interval around the robust mean.

Priming as a function of reward cost, reward strength, and D2-like blockade: Supporting information

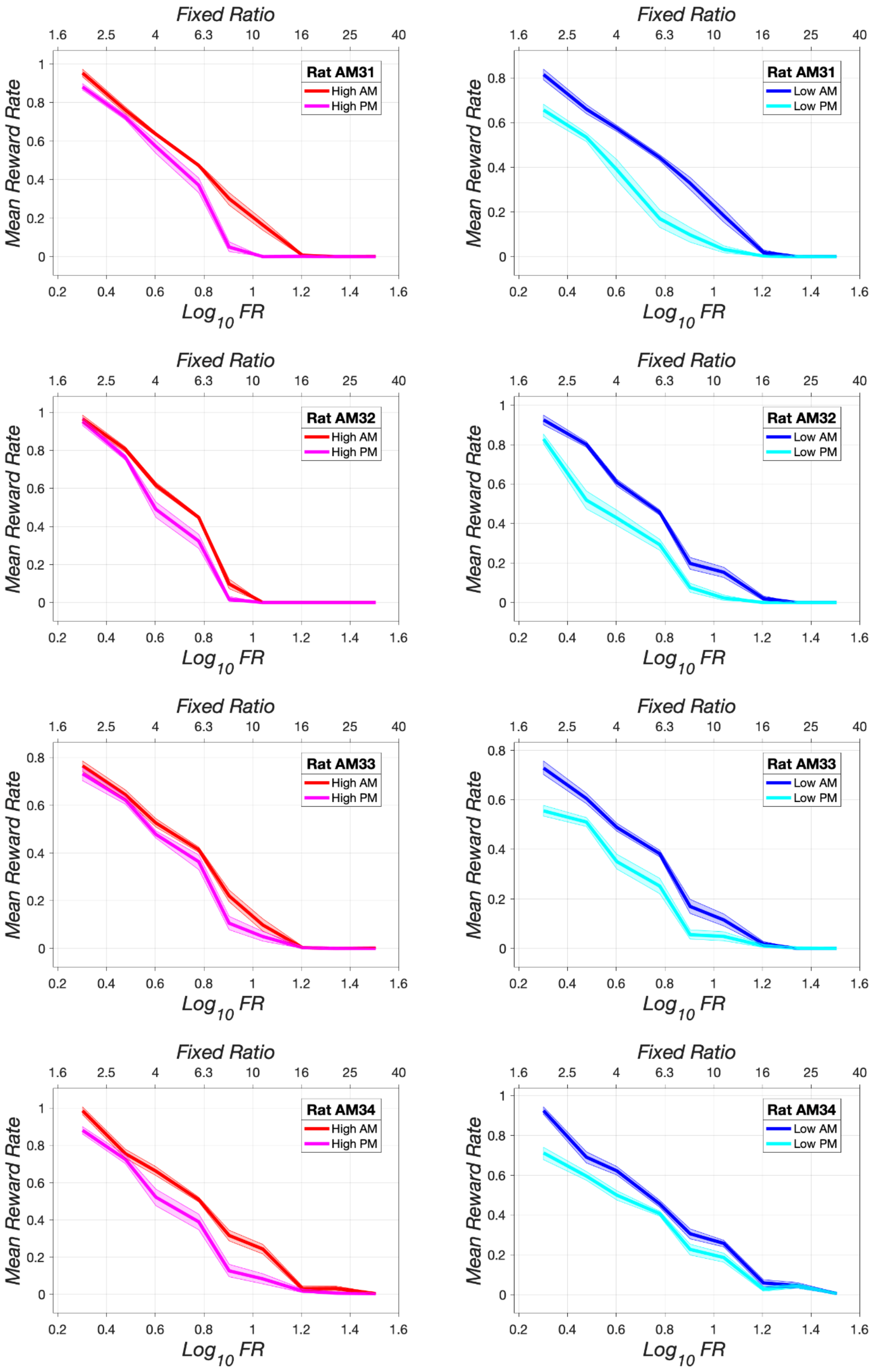

***Figure S2:*** Mean reward rate as a function of reward cost (fixed-ratio requirement) in morning (AM) and afternoon (PM) test sessions under low(right) and high(left)  priming for rats AM31, AM32, AM33, and AM34. The error band shows the 95% confidence interval around the robust mean.

Priming as a function of reward cost, reward strength, and D2-like blockade: Supporting information

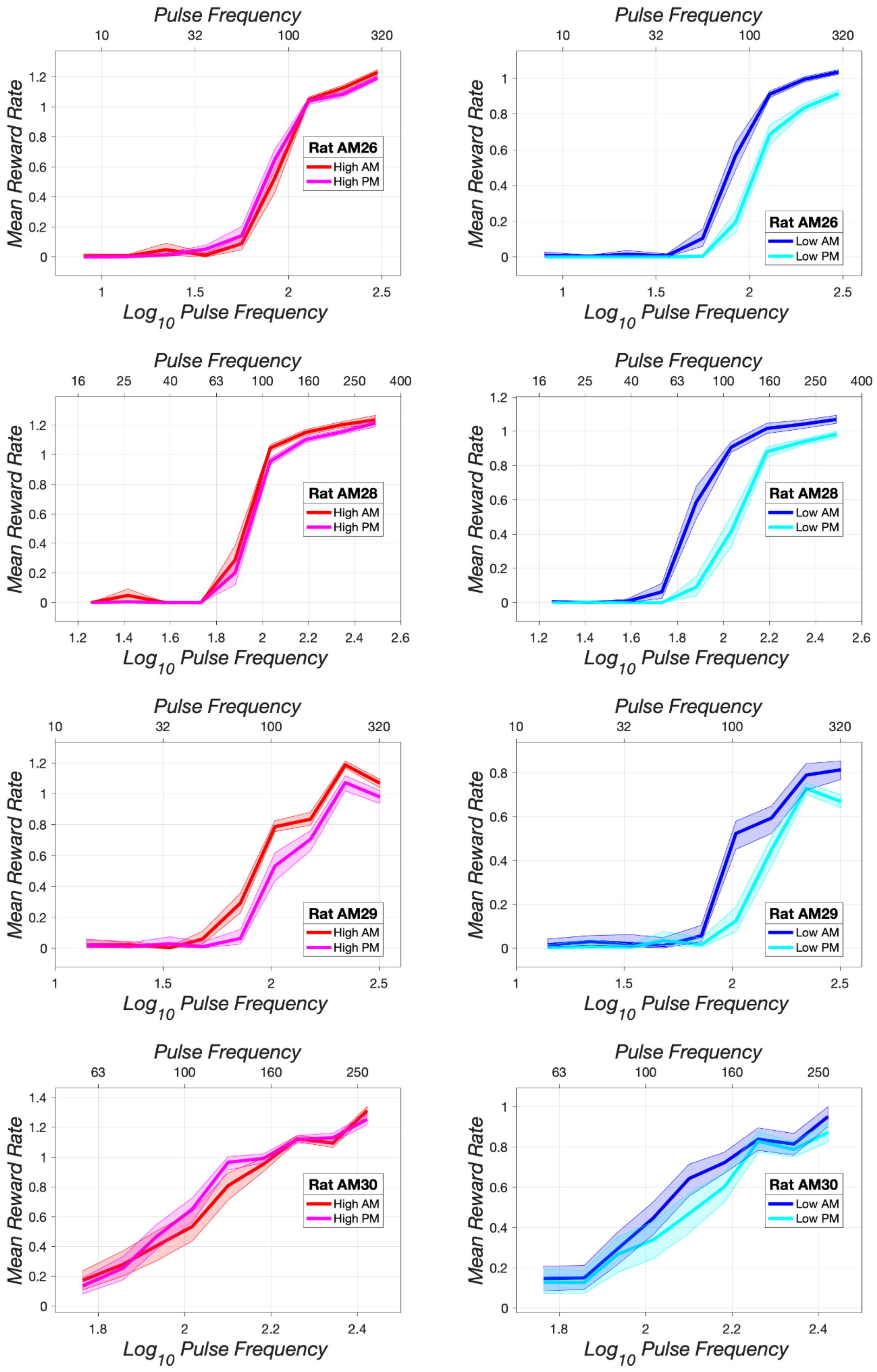

***Figure S3:*** Mean reward rate as a function of stimulation strength (pulse frequency) in morning (AM) and afternoon (PM) test sessions under low(right) and high(left) priming for rats AM26, AM28, AM29, and AM30. The error band shows the 95% confidence interval around the robust mean.

Priming as a function of reward cost, reward strength, and D2-like blockade: Supporting information

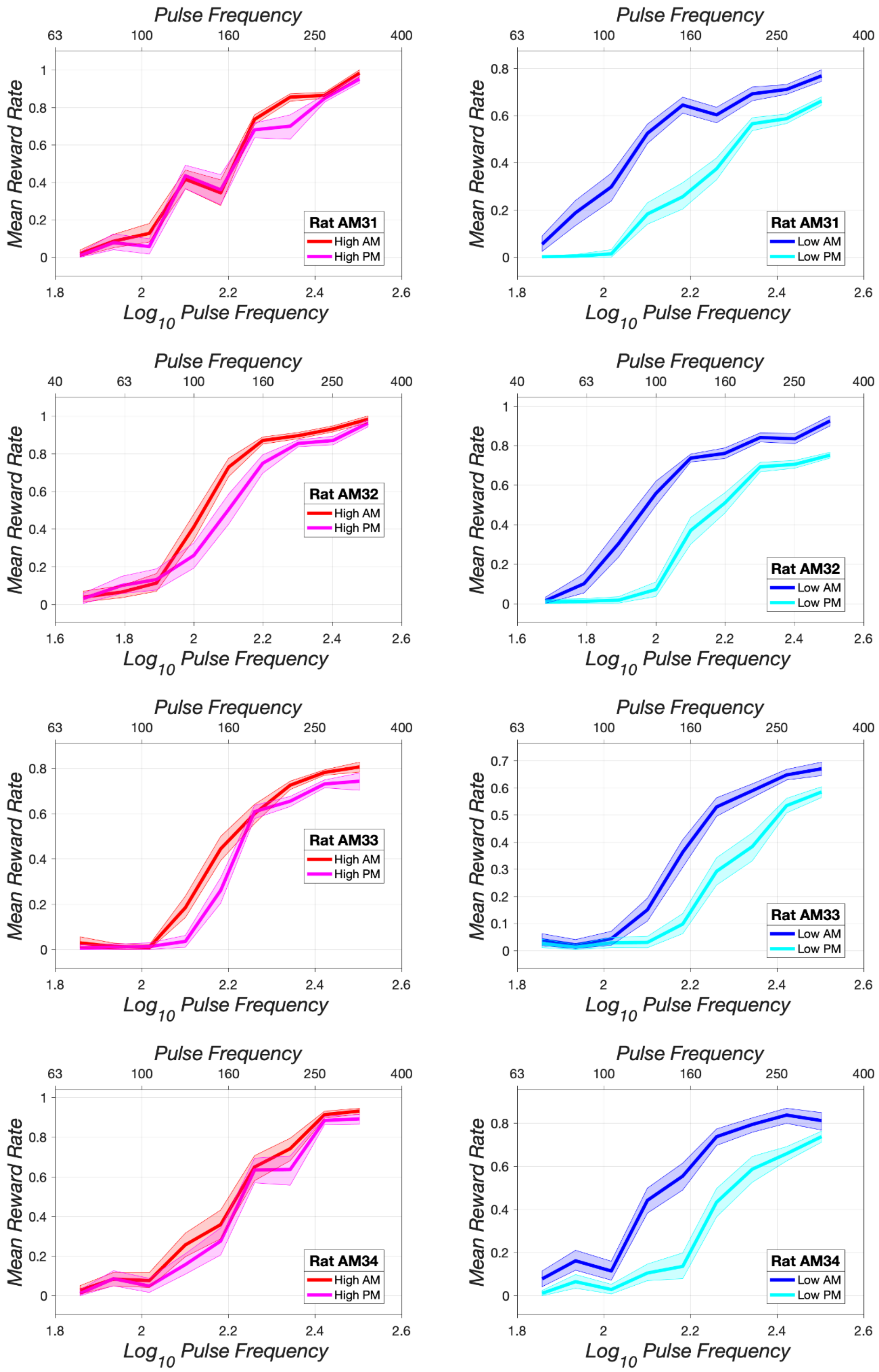

***Figure S4:*** Mean reward rate as a function of stimulation strength (pulse frequency) in morning (AM) and afternoon (PM) test sessions under low(right) and high(left) priming for rats AM31, AM232, AM33, and AM34. The error band shows the 95% confidence interval around the robust mean.

Priming as a function of reward cost, reward strength, and D2-like blockade: Supporting information

| Rat | RobustMeanAM | RobustMeanPM | GrandRobustMean | DiffRatio | Cliffs_delta | Cliffs_delta_CI_hi | Cliffs_delta_CI_lo | met both |
|---|---|---|---|---|---|---|---|---|
| AM26 | 1.033 | 0.922 | 0.985 | 0.113 | 0.753 | 0.853 | 0.632 | TRUE |
| AM28 | 1.067 | 0.987 | 1.024 | 0.078 | 0.497 | 0.651 | 0.328 | TRUE |
| AM29 | 0.818 | 0.671 | 0.740 | 0.199 | 0.486 | 0.647 | 0.306 | TRUE |
| AM30 | 0.953 | 0.867 | 0.915 | 0.094 | 0.189 | 0.419 | -0.047 | FALSE |
| AM31 | 0.765 | 0.662 | 0.704 | 0.146 | 0.528 | 0.651 | 0.392 | TRUE |
| AM32 | 0.929 | 0.747 | 0.822 | 0.222 | 0.761 | 0.851 | 0.661 | TRUE |
| AM33 | 0.672 | 0.593 | 0.627 | 0.126 | 0.398 | 0.528 | 0.257 | TRUE |
| AM34 | 0.866 | 0.746 | 0.802 | 0.149 | 0.468 | 0.614 | 0.315 | TRUE |
| Median | 0.898 | 0.747 | 0.812 | 0.136 | 0.492 | 0.649 | 0.322 | |
| Max | 1.067 | 0.987 | 1.024 | 0.222 | 0.761 | 0.853 | 0.661 | |
| Min | 0.672 | 0.593 | 0.627 | 0.078 | 0.189 | 0.419 | -0.047 | |
| met criterion | | | | 8 | | | 7 | 7 |

***Table S1:*** Effect of session order on reward rates in the lowest-cost trial of the stimulation-strength (pulse frequency) sweep under low priming.

| Rat | RobustMeanAM | RobustMeanPM | GrandRobustMean | DiffRatio | Cliffs_delta | Cliffs_delta_CI_hi | Cliffs_delta_CI_lo | met both |
|---|---|---|---|---|---|---|---|---|
| AM26 | 1.238 | 1.195 | 1.217 | 0.035 | 0.352 | 0.528 | 0.162 | FALSE |
| AM28 | 1.274 | 1.224 | 1.250 | 0.040 | 0.247 | 0.444 | 0.044 | FALSE |
| AM29 | 1.078 | 0.983 | 1.042 | 0.091 | 0.351 | 0.501 | 0.194 | TRUE |
| AM30 | 1.331 | 1.272 | 1.300 | 0.045 | 0.272 | 0.464 | 0.069 | FALSE |
| AM31 | 0.990 | 0.958 | 0.977 | 0.032 | 0.208 | 0.358 | 0.050 | FALSE |
| AM32 | 0.988 | 0.969 | 0.979 | 0.020 | 0.152 | 0.298 | -0.003 | FALSE |
| AM33 | 0.834 | 0.788 | 0.816 | 0.057 | 0.204 | 0.347 | 0.060 | TRUE |
| AM34 | 0.939 | 0.909 | 0.926 | 0.032 | 0.189 | 0.336 | 0.044 | FALSE |
| Median | 1.034 | 0.976 | 1.011 | 0.037 | 0.227 | 0.401 | 0.055 | |
| Max | 1.331 | 1.272 | 1.300 | 0.091 | 0.352 | 0.528 | 0.194 | |
| Min | 0.834 | 0.788 | 0.816 | 0.020 | 0.152 | 0.298 | -0.003 | |
| met criterion | | | | 2 | | | 7 | 2 |

***Table S2:*** Effect of session order on reward rates in the lowest-cost trial of the stimulation-strength (pulse frequency) sweep under high priming.

| Rat | DiffRatio | Cliffs_delta | Cliffs_delta_CI_hi | Cliffs_delta_CI_lo | met both |
|---|---|---|---|---|---|
| AM26 | -0.078 | -0.401 | -0.325 | -0.470 | TRUE |
| AM28 | -0.039 | -0.250 | -0.207 | -0.284 | FALSE |
| AM29 | -0.107 | -0.135 | -0.146 | -0.112 | TRUE |
| AM30 | -0.049 | 0.083 | 0.045 | 0.116 | FALSE |
| AM31 | -0.114 | -0.320 | -0.293 | -0.342 | TRUE |
| AM32 | -0.202 | -0.609 | -0.553 | -0.664 | TRUE |
| AM33 | -0.068 | -0.194 | -0.182 | -0.198 | TRUE |
| AM34 | -0.117 | -0.279 | -0.278 | -0.271 | TRUE |
| Median | -0.092 | -0.265 | -0.242 | -0.277 | |
| Max | -0.039 | 0.083 | 0.045 | 0.116 | |
| Min | -0.202 | -0.609 | -0.553 | -0.664 | |
| met criterion | 7 | | | 7 | 6 |

***Table S3:*** Differences in the effect of session order on reward rates in the lowest-cost trial of the stimulation-strength (pulse frequency) sweep under high and low priming (high-low).

Priming as a function of reward cost, reward strength, and D2-like blockade: Supporting information

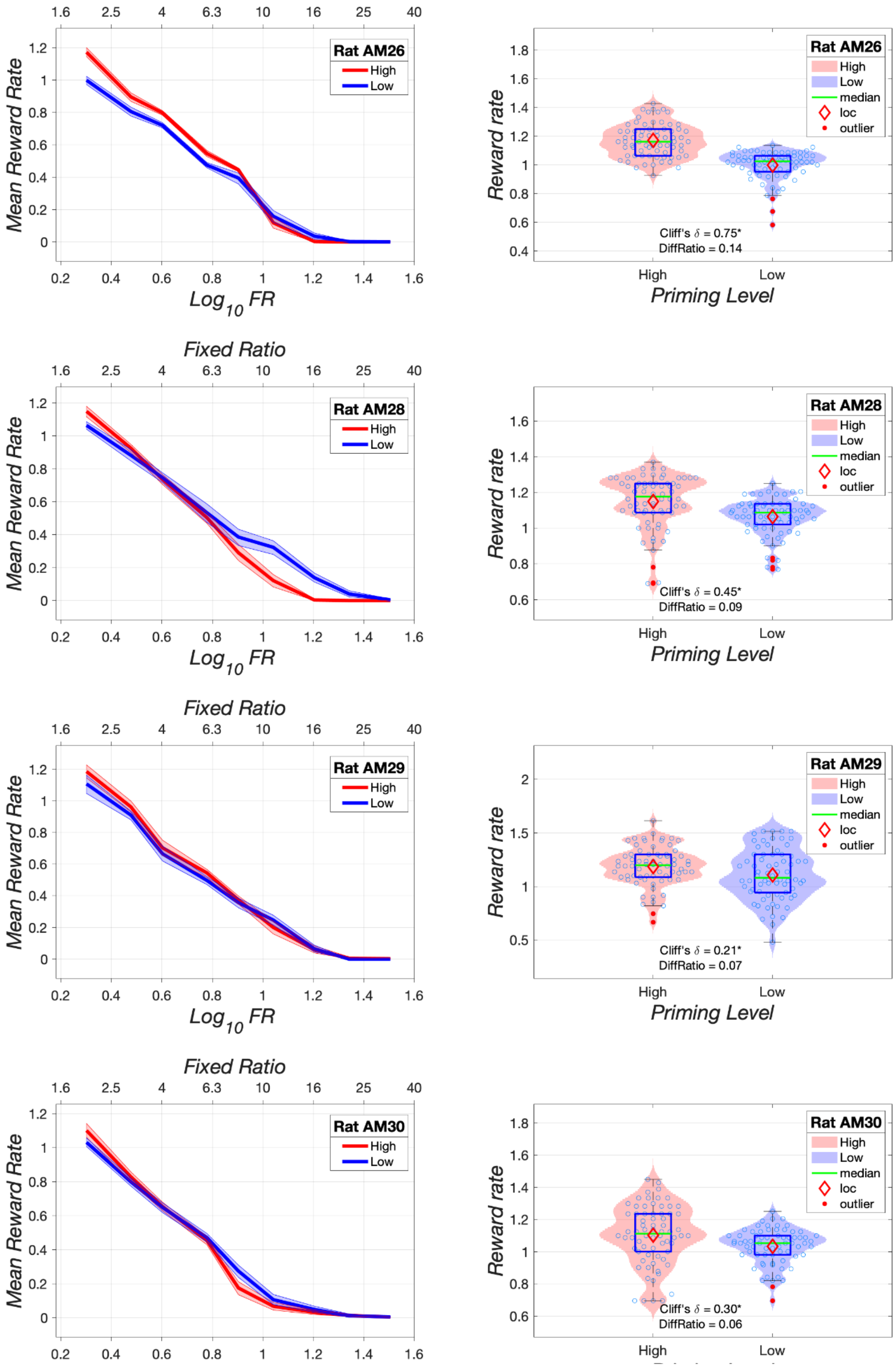

***Figure S5***: Effect of reward cost (fixed-ratio requirement) on reward rate under high (red) and low (blue) priming for rats AM26, AM28, AM29, and AM30. The blue box in the violin plots represents the interquartile range, and the red diamond denotes the robust mean (the location parameter of the Tukey bisquare estimator). The lateral position of the open blue circles in the violin plots has been jittered to minimize overlap. Data are from morning sessions.

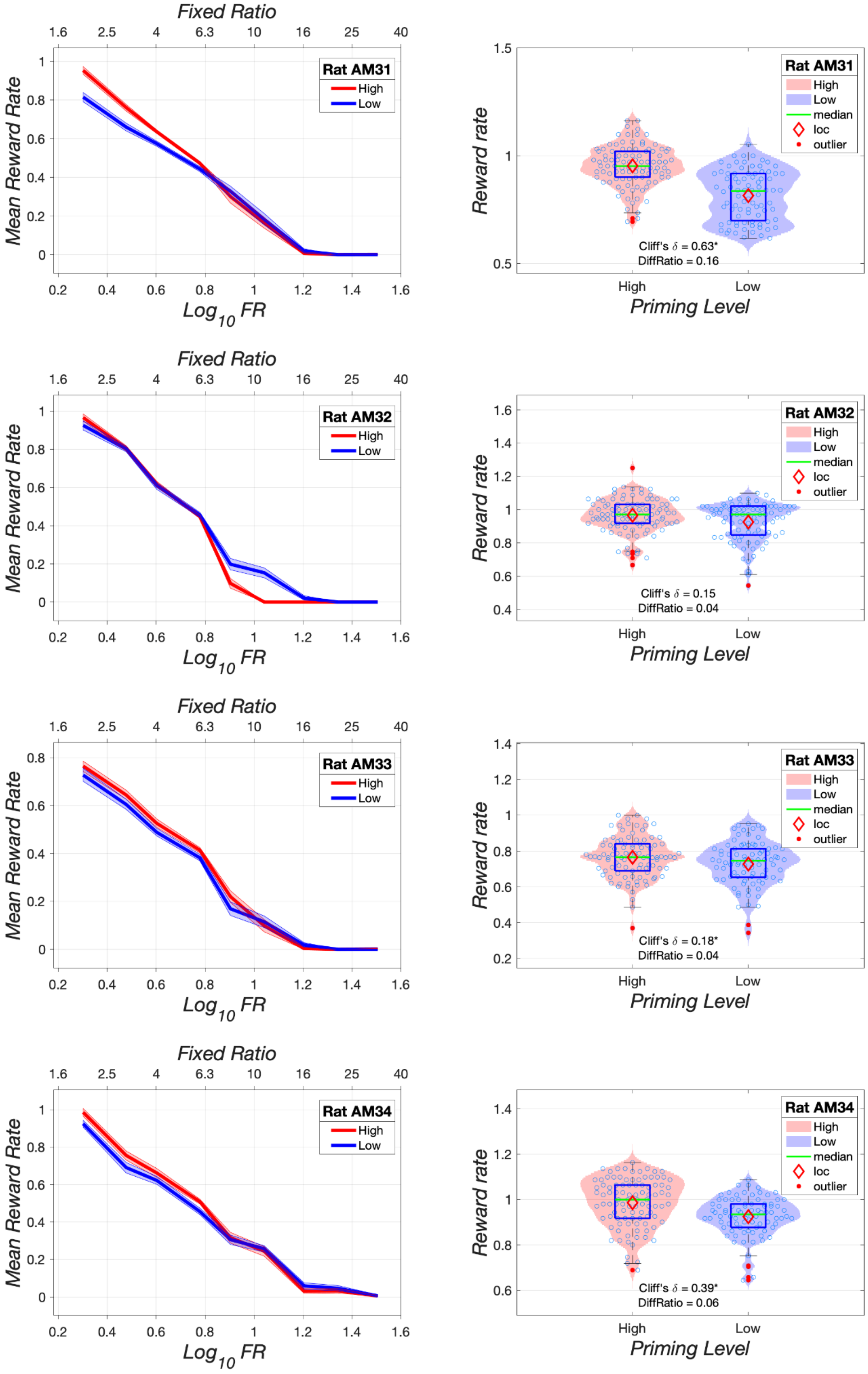
Fixed Ratio
Mean Reward Rate
$Log_{10}$ FR
Rat AM31
High
Low
Reward rate
Priming Level
median
loc
outlier
Cliff's δ = 0.63*
DiffRatio = 0.16
Rat AM32
Cliff's δ = 0.15
DiffRatio = 0.04
Rat AM33
Cliff's δ = 0.18*
DiffRatio = 0.04
Rat AM34
Cliff's δ = 0.39*
DiffRatio = 0.06

***Figure S6***: Effect of reward cost (fixed-ratio requirement) on reward rate under high (red) and low (blue) priming for rats AM31, AM232, AM33, and AM34. The blue box in the violin plots represents the interquartile range, and the red diamond denotes the robust mean (the location parameter of the Tukey bisquare estimator). The lateral position of the open blue circles in the violin plots has been jittered to minimize overlap. Data are from morning sessions.

Priming as a function of reward cost, reward strength, and D2-like blockade: Supporting information

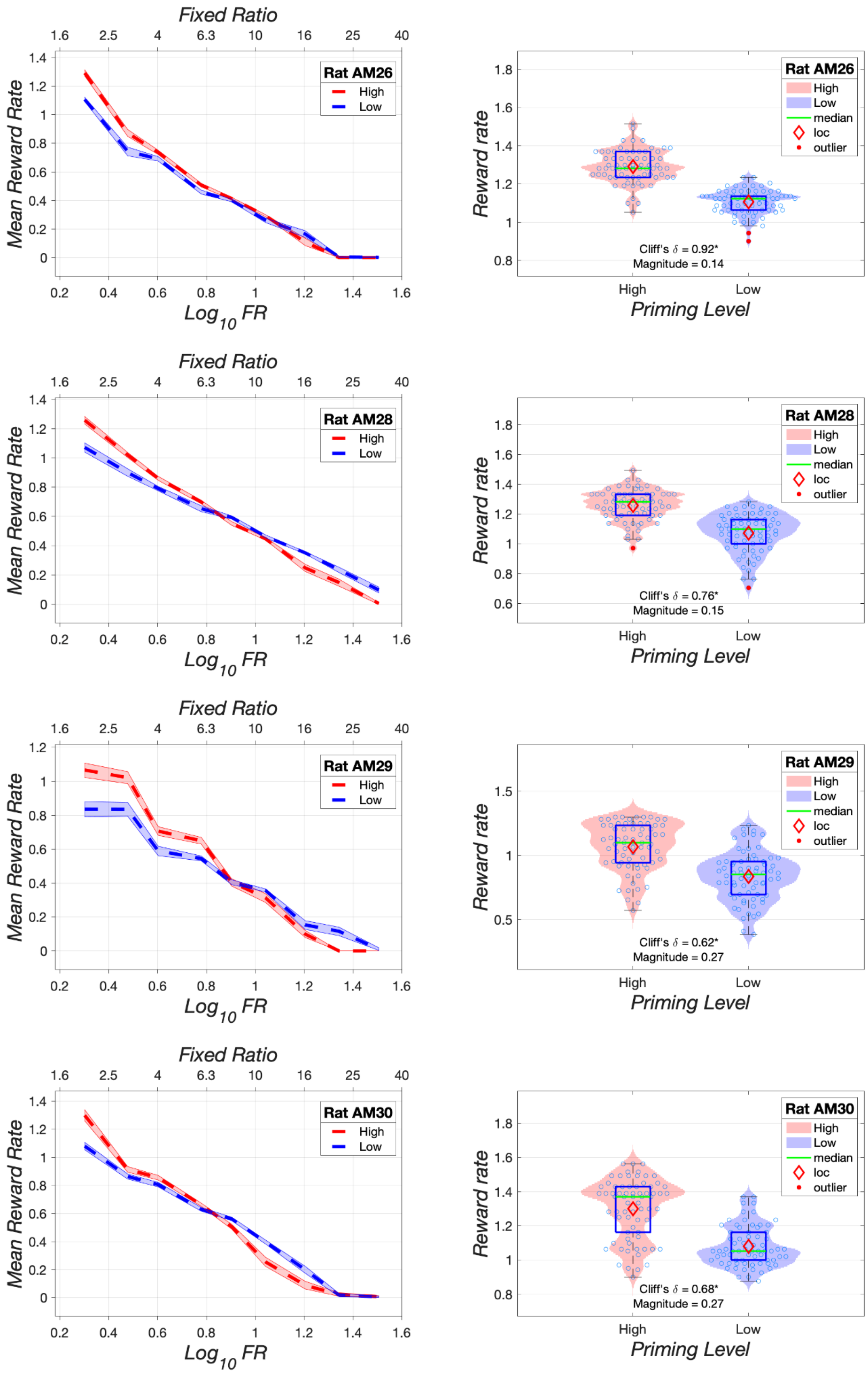

***Figure S7***: Retest of the effect of reward cost (fixed-ratio requirement) on reward rate under high (red) and low (blue) priming for rats AM26, AM28, AM29, and AM30. The blue box in the violin plots represents the interquartile range, and the red diamond denotes the robust mean (the location parameter of the Tukey bisquare estimator). The lateral position of the open blue circles in the violin plots has been jittered to minimize overlap. Data are from morning sessions.

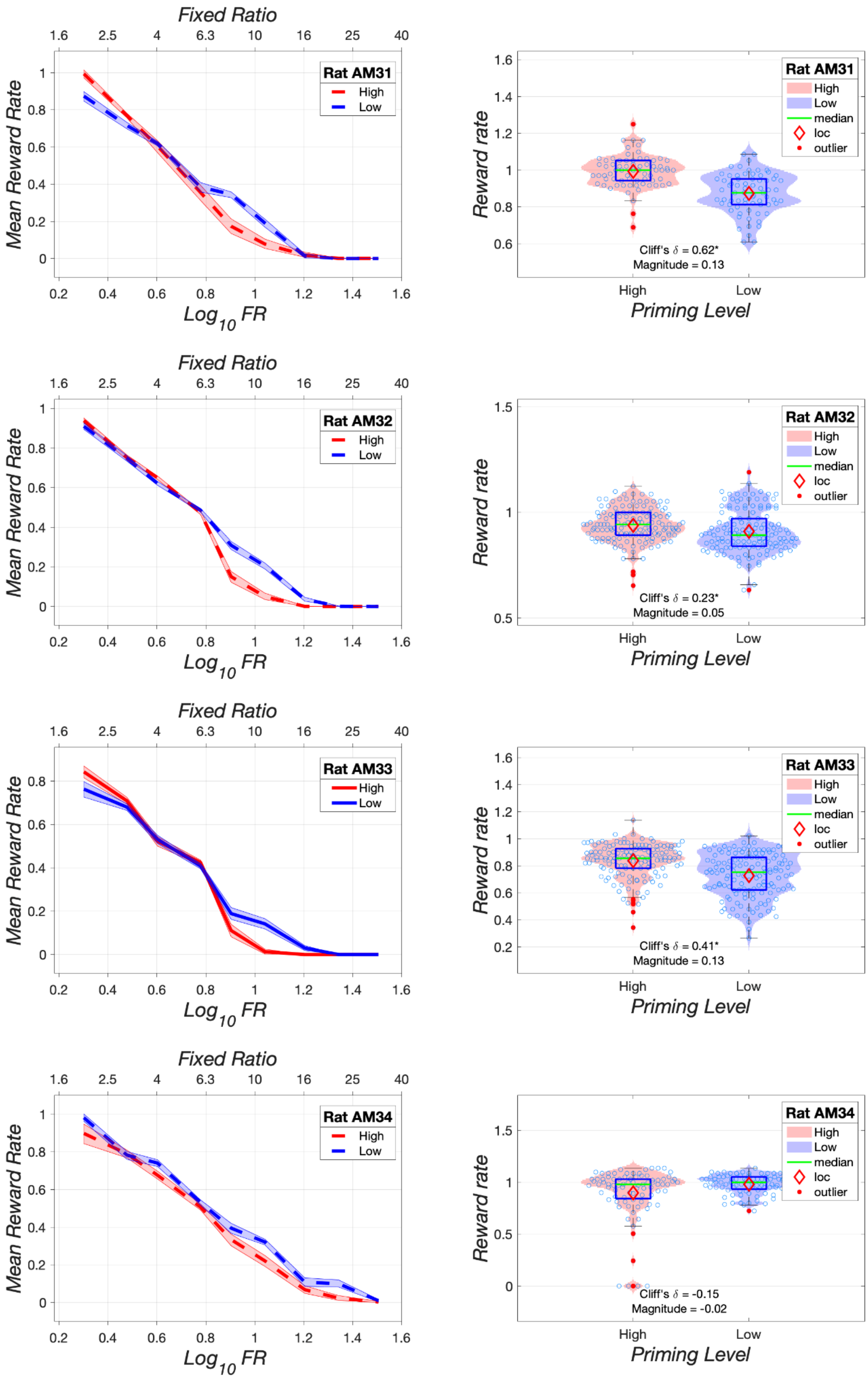
Fixed Ratio
1.6 2.5 4 6.3 10 16 25 40
Mean Reward Rate
Log$_{10}$ FR
0.2 0.4 0.6 0.8 1 1.2 1.4 1.6
Rat AM31
High
Low
Reward rate
Priming Level
median
loc
outlier
Cliff's δ = 0.62*
Magnitude = 0.13
Rat AM32
Cliff's δ = 0.23*
Magnitude = 0.05
Rat AM33
Cliff's δ = 0.41*
Magnitude = 0.13
Rat AM34
Cliff's δ = -0.15
Magnitude = -0.02

***Figure S8***: Retest of the effect of reward cost (fixed-ratio requirement) on reward rate under high (red) and low (blue) priming for rats AM31, AM232, AM33, and AM34.  The blue box in the violin plots represents the interquartile range, and the red diamond denotes the robust mean (the location parameter of the Tukey bisquare estimator). The lateral position of the open blue circles in the violin plots has been jittered to minimize overlap. Data are from morning sessions.

Priming as a function of reward cost, reward strength, and D2-like blockade: Supporting information

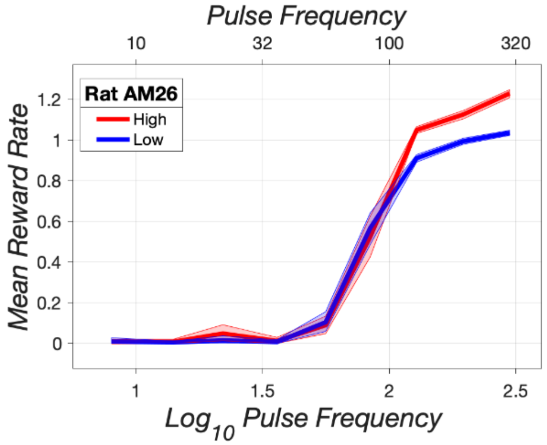


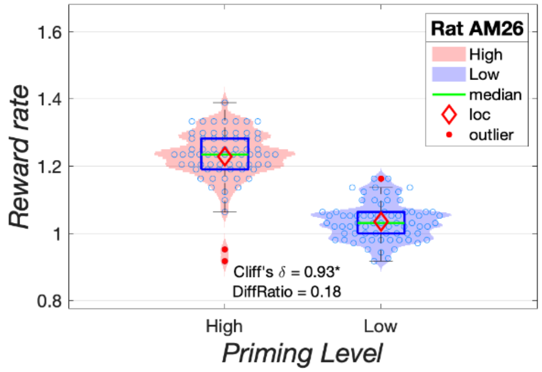


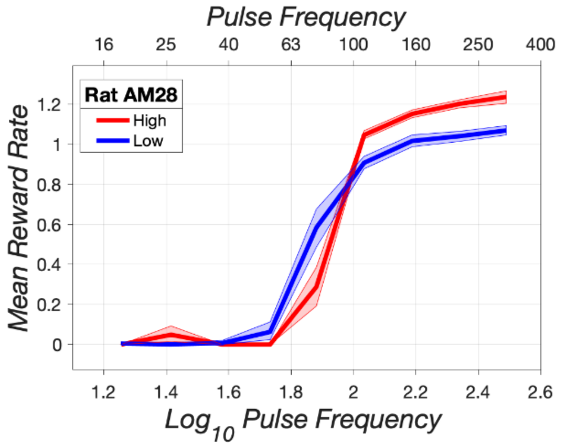


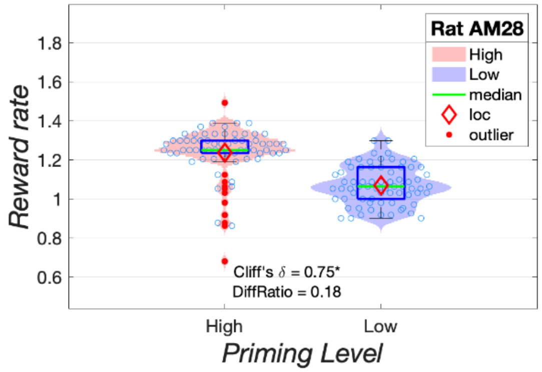


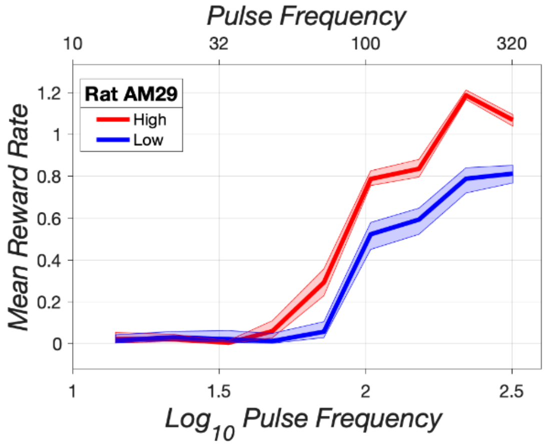


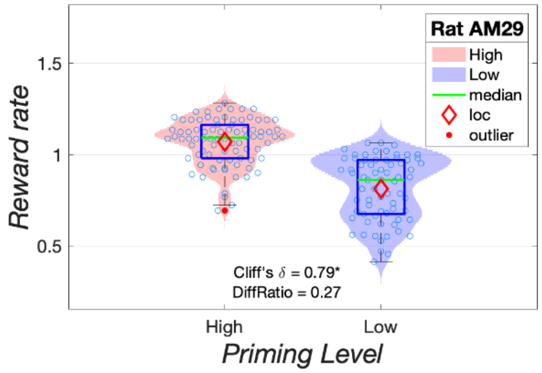


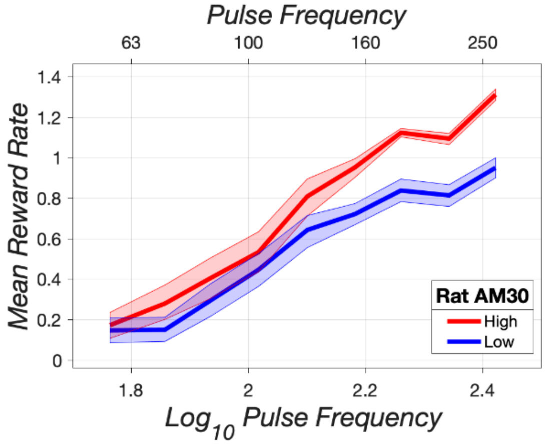


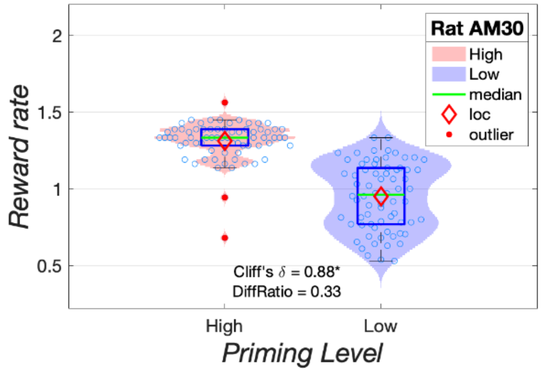

***Figure S9***: Effect of reward strength (pulse frequency) on reward rate under high (red) and low (blue) priming for rats AM26, AM28, AM29, and AM30. The blue box in the violin plots represents the interquartile range, and the red diamond denotes the robust mean (the location parameter of the Tukey bisquare estimator). The lateral position of the open blue circles in the violin plots has been jittered to minimize overlap. Data are from morning sessions.

Priming as a function of reward cost, reward strength, and D2-like blockade: Supporting information

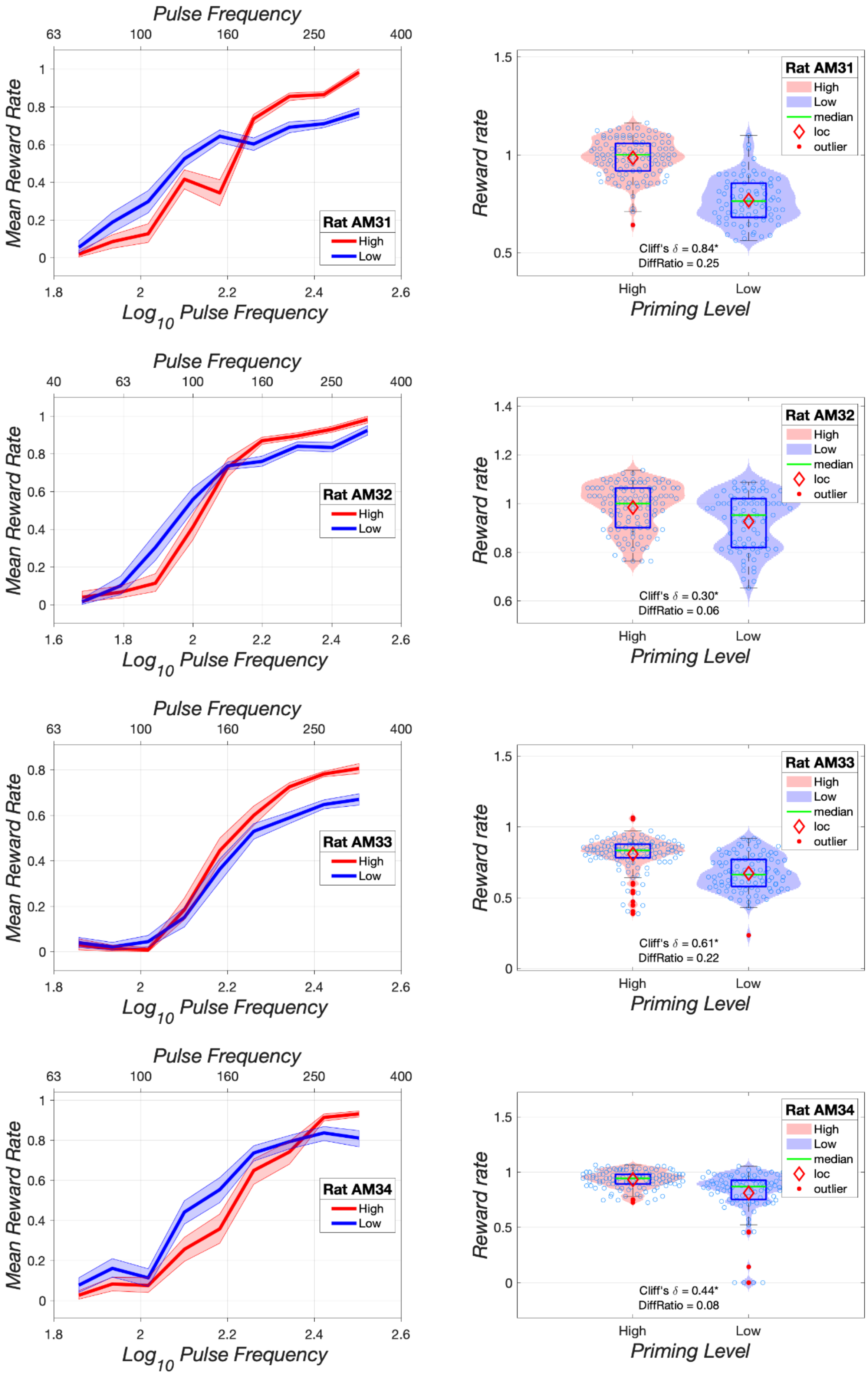

***Figure S10***: Effect of reward strength (pulse frequency) on reward rate under high (red) and low (blue) priming for rats AM31, AM232, AM33, and AM34. The blue box in the violin plots represents the interquartile range, and the red diamond denotes the robust mean (the location parameter of the Tukey bisquare estimator). The lateral position of the open blue circles in the violin plots has been jittered to minimize overlap. Data are from morning sessions.

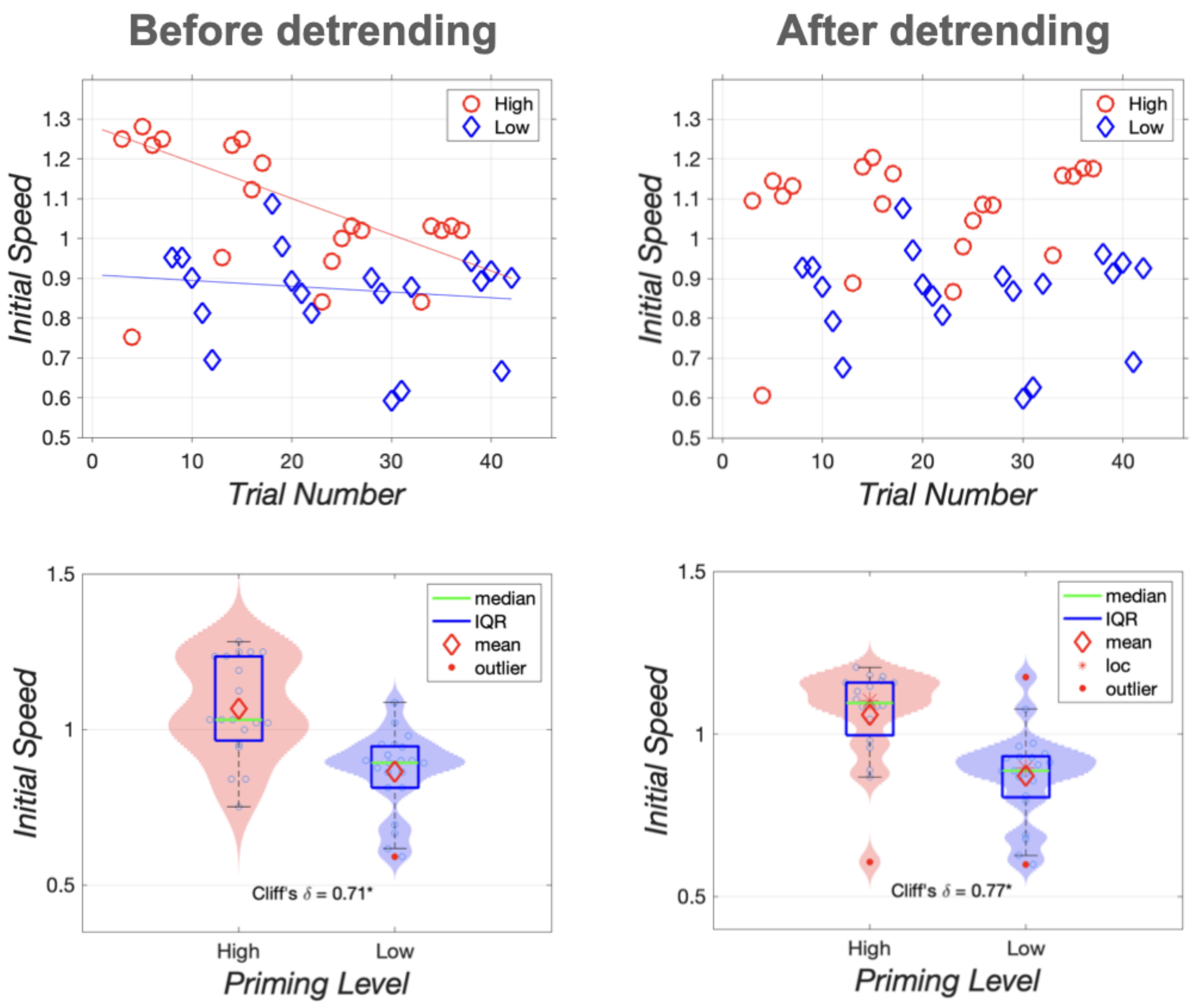


***Figure S10***: Effect of detrending. The left column shows the untransformed data and their distributions, whereas the right column shows the transformed data and distributions after detrending. Data are from rat AM46, session 6.

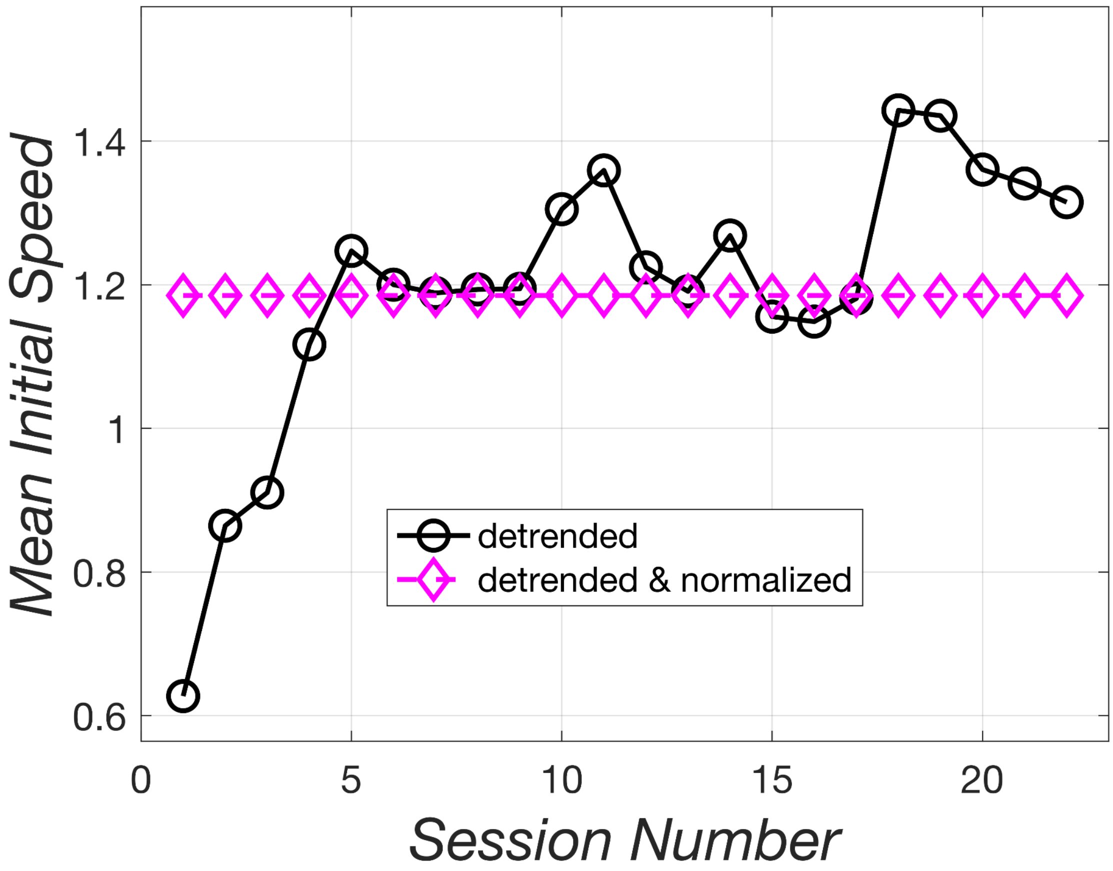


Figure S11: Average vigor varies from session to session. Black circles are the robust means of the initial-speed measure across priming conditions, whereas the magenta diamonds show the corresponding normalized values.

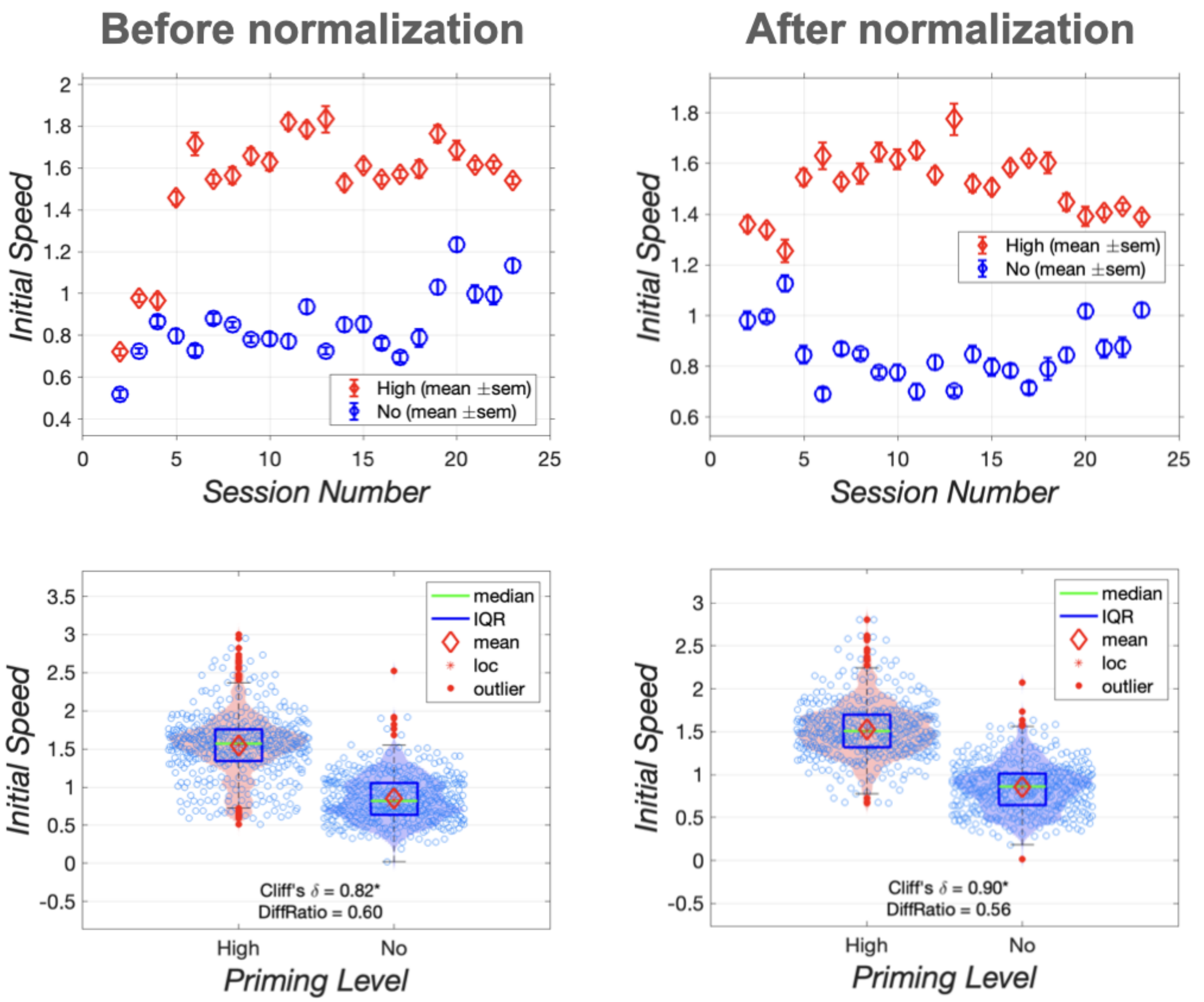


***Figure S12***: Effect of normalization to remove influence of session-to-session fluctuation in average vigor. Data are from rat AM46.

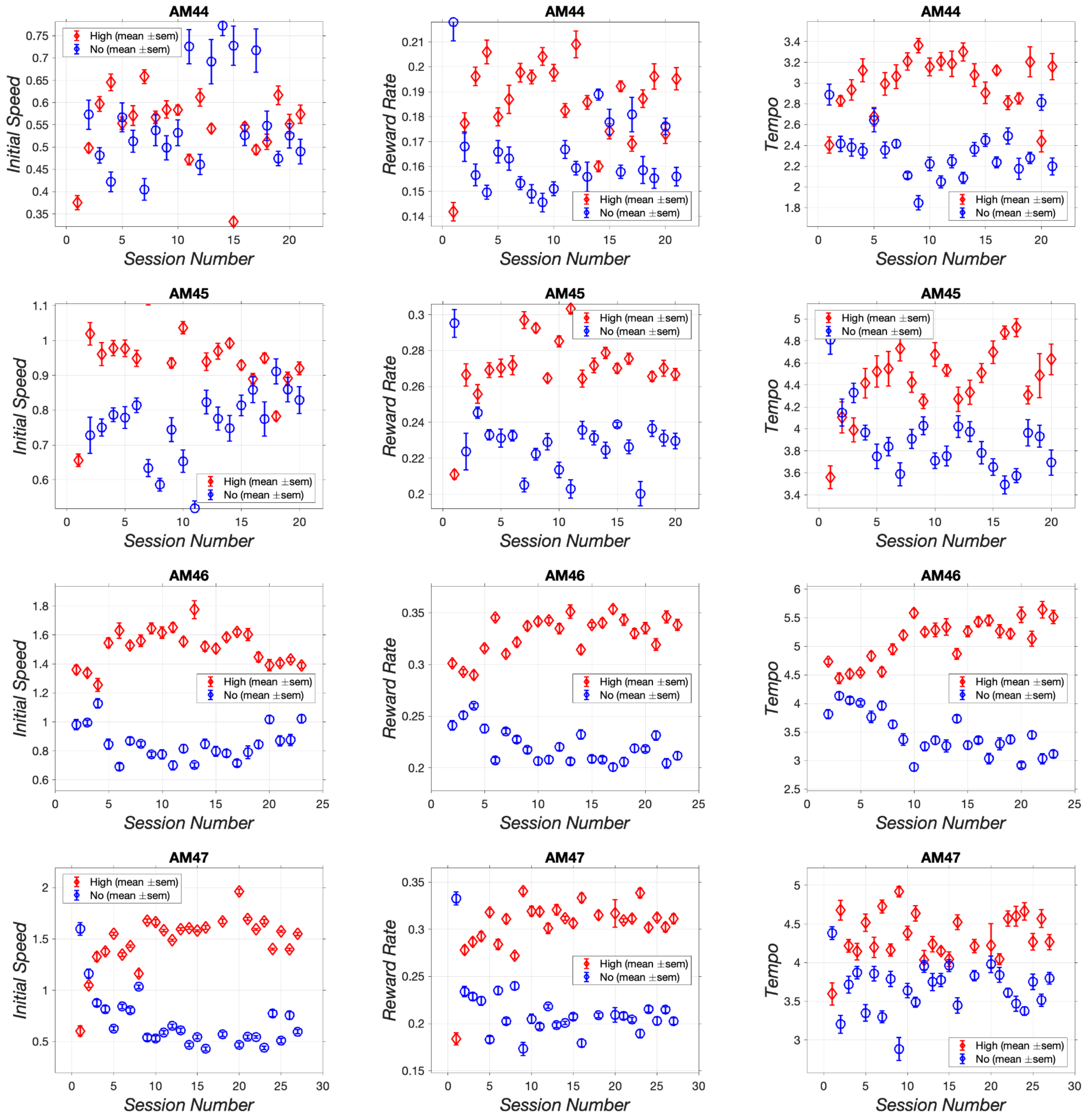


***Figure S13a***: Session-by-session vigor scores collapsed over trials for rats AM44-AM47.

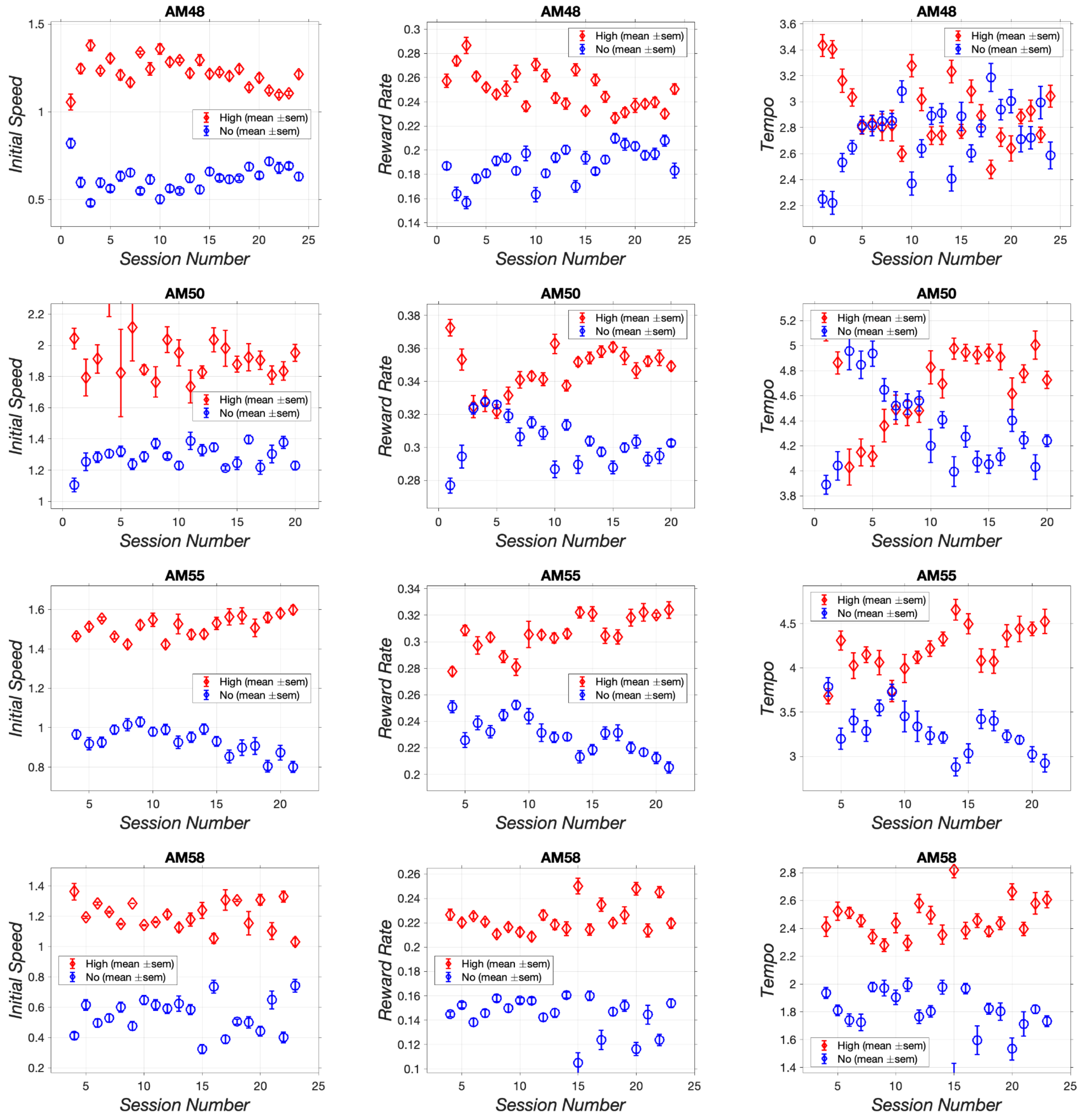


***Figure S13b:*** Session-by-session vigor scores collapsed over trials for rats AM48, AM50, AM55, and AM58.

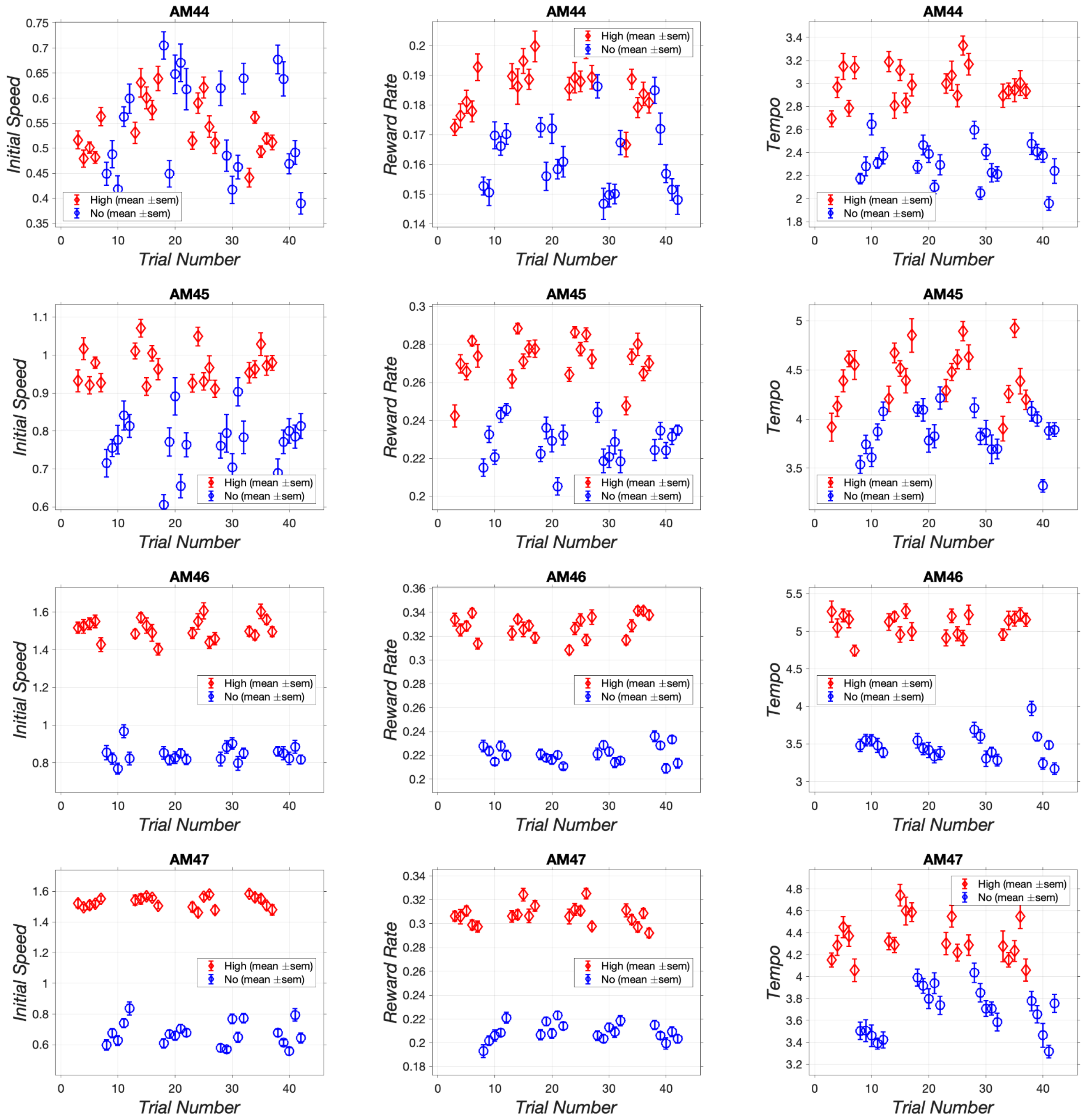


***Figure S14a:*** Trial-by-trial vigor scores collapsed over sessions for rats AM44-AM47.

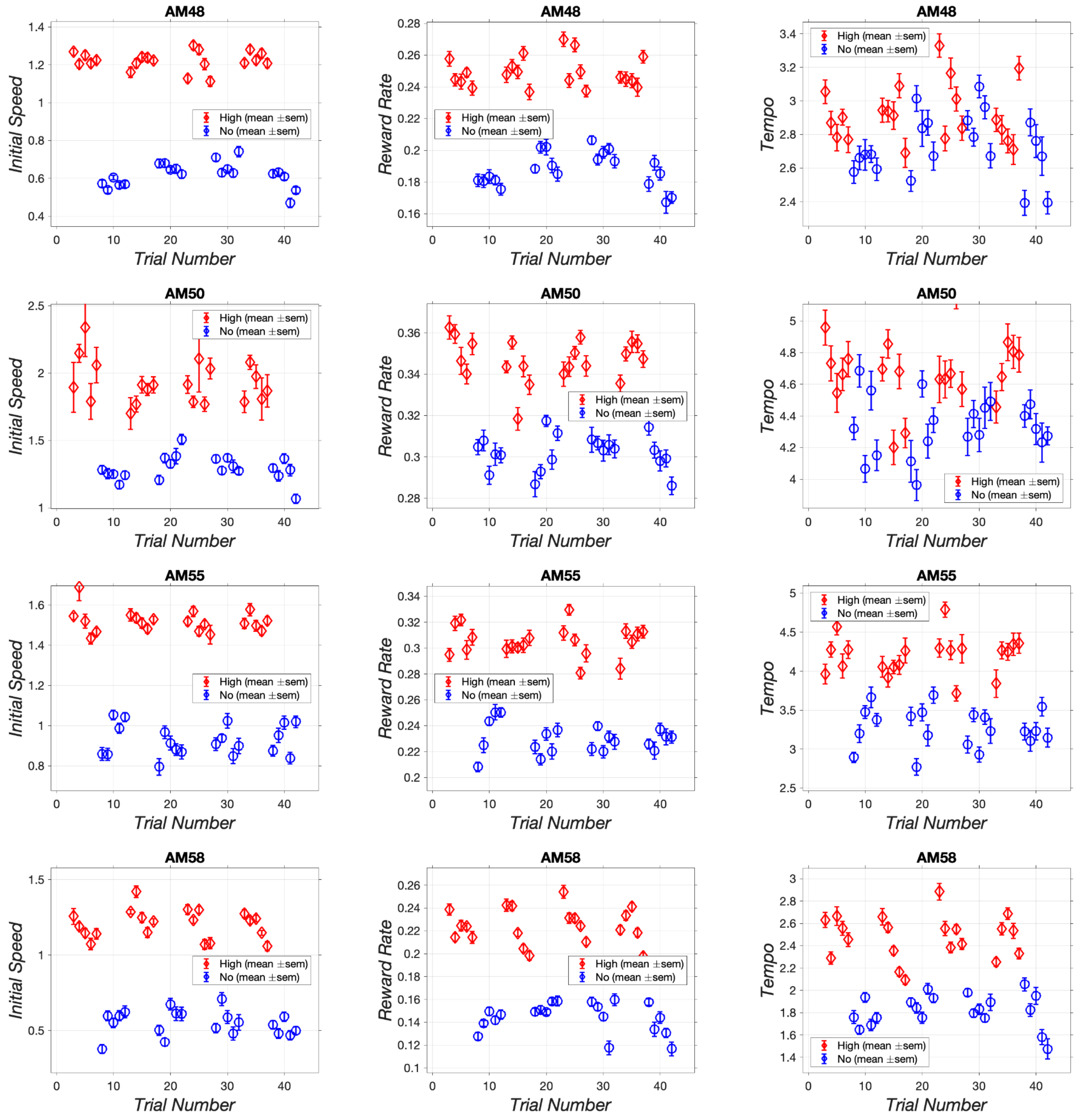


***Figure S14b:*** Trial-by-trial vigor scores collapsed over sessions for rats AM48, AM50, AM55, and AM58.

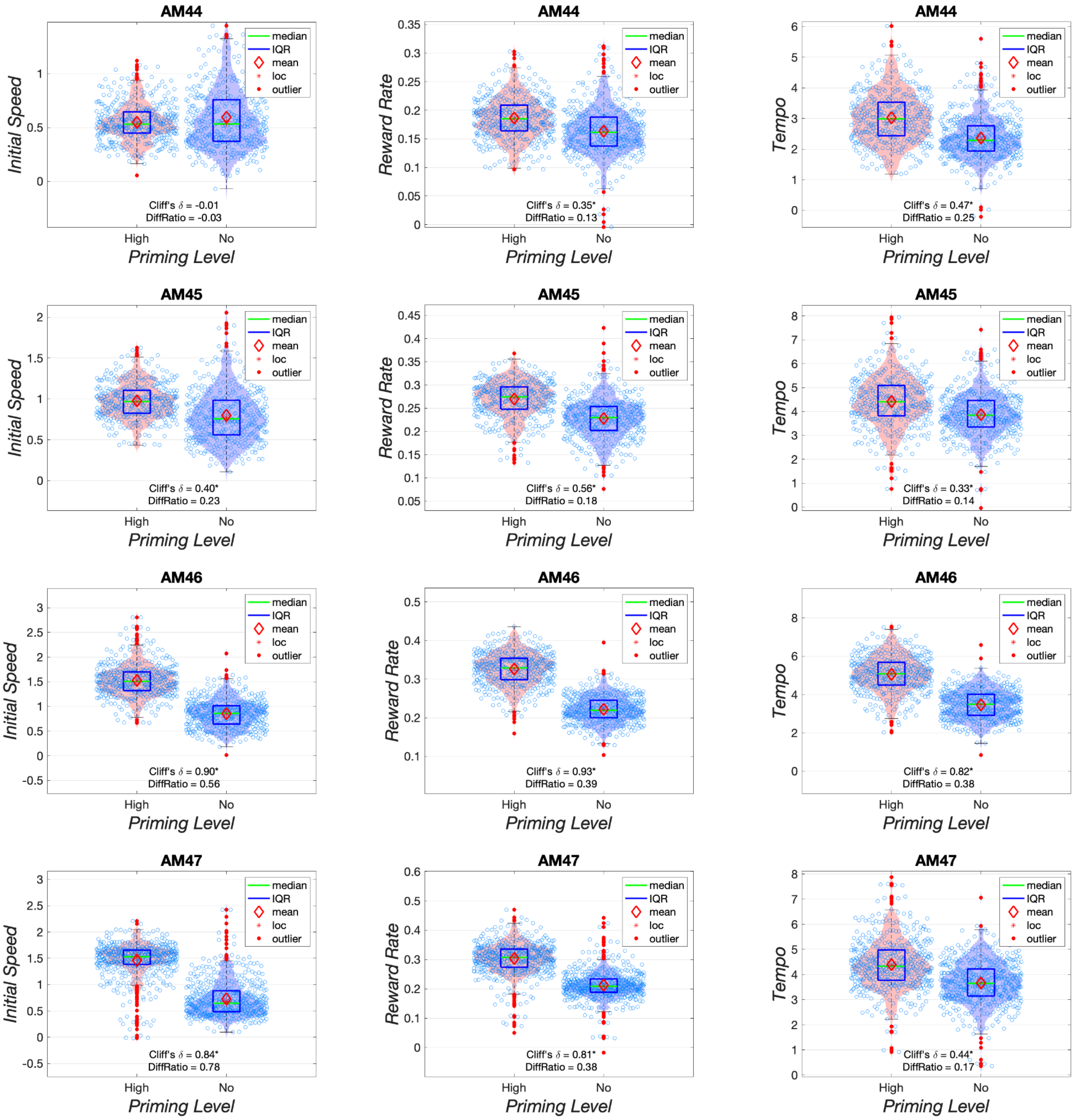


***Figure S15a:*** Distributions of vigor scores for rats AM44-AM47.

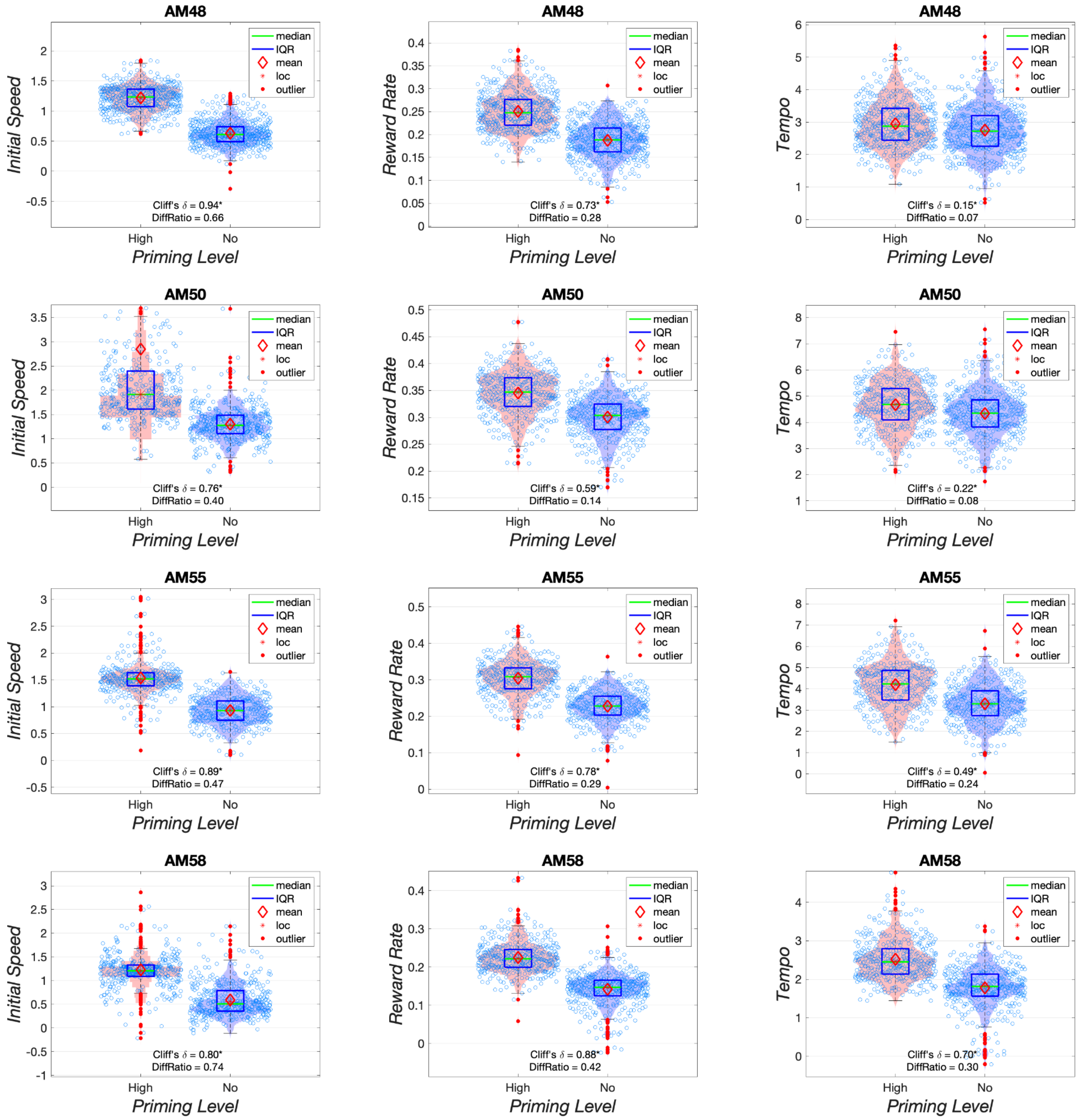


***Figure S15b:*** Distributions of vigor scores for rats AM48, AM50, AM55, and AM58.

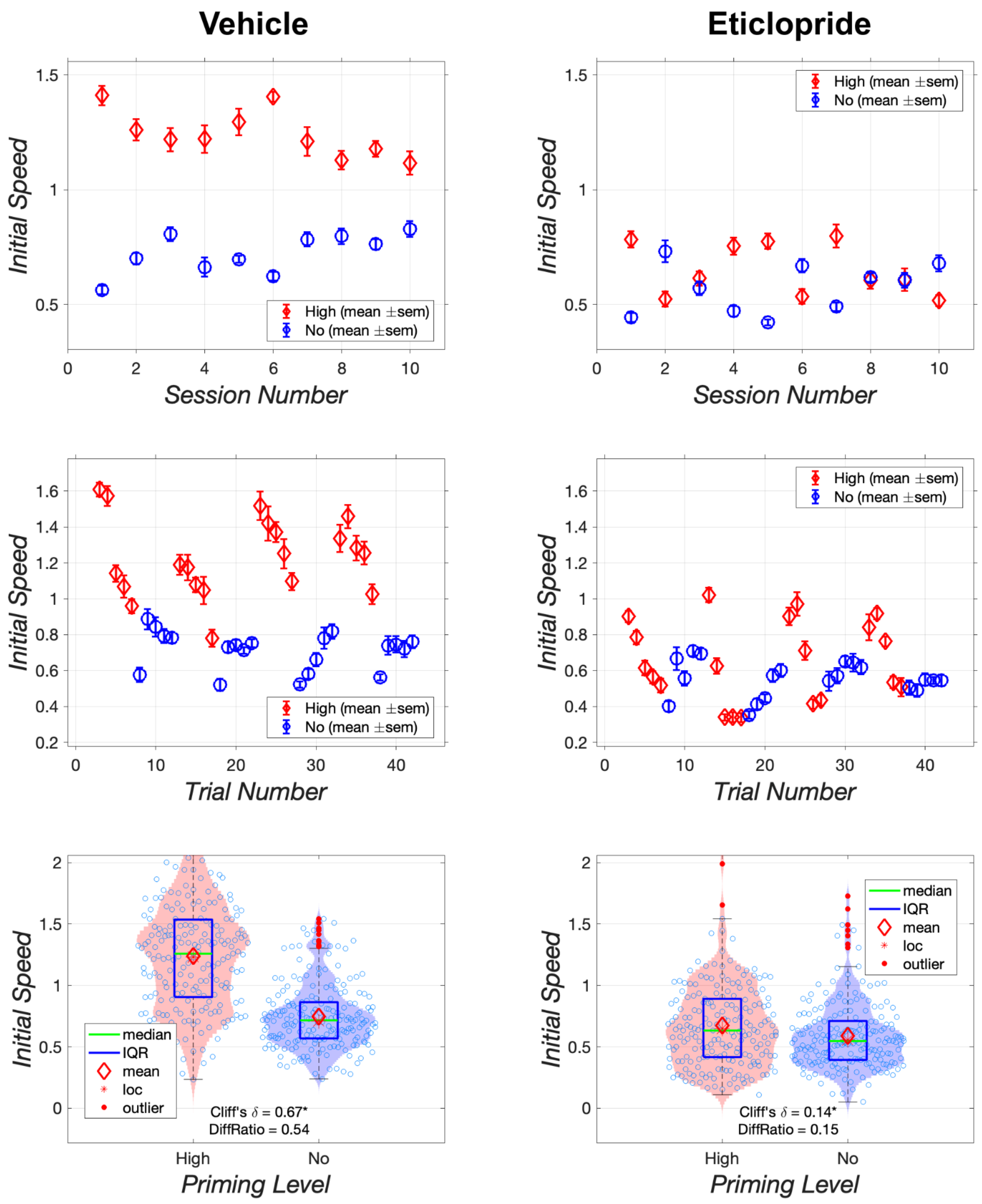


***Figure S16:*** Effect of vehicle and 0.1 mg/kg of eticlopride on the initial speed to initiate responding by rat AM55. The upper row shows results collapsed over trials, whereas the middle row shows results collapsed over sessions.

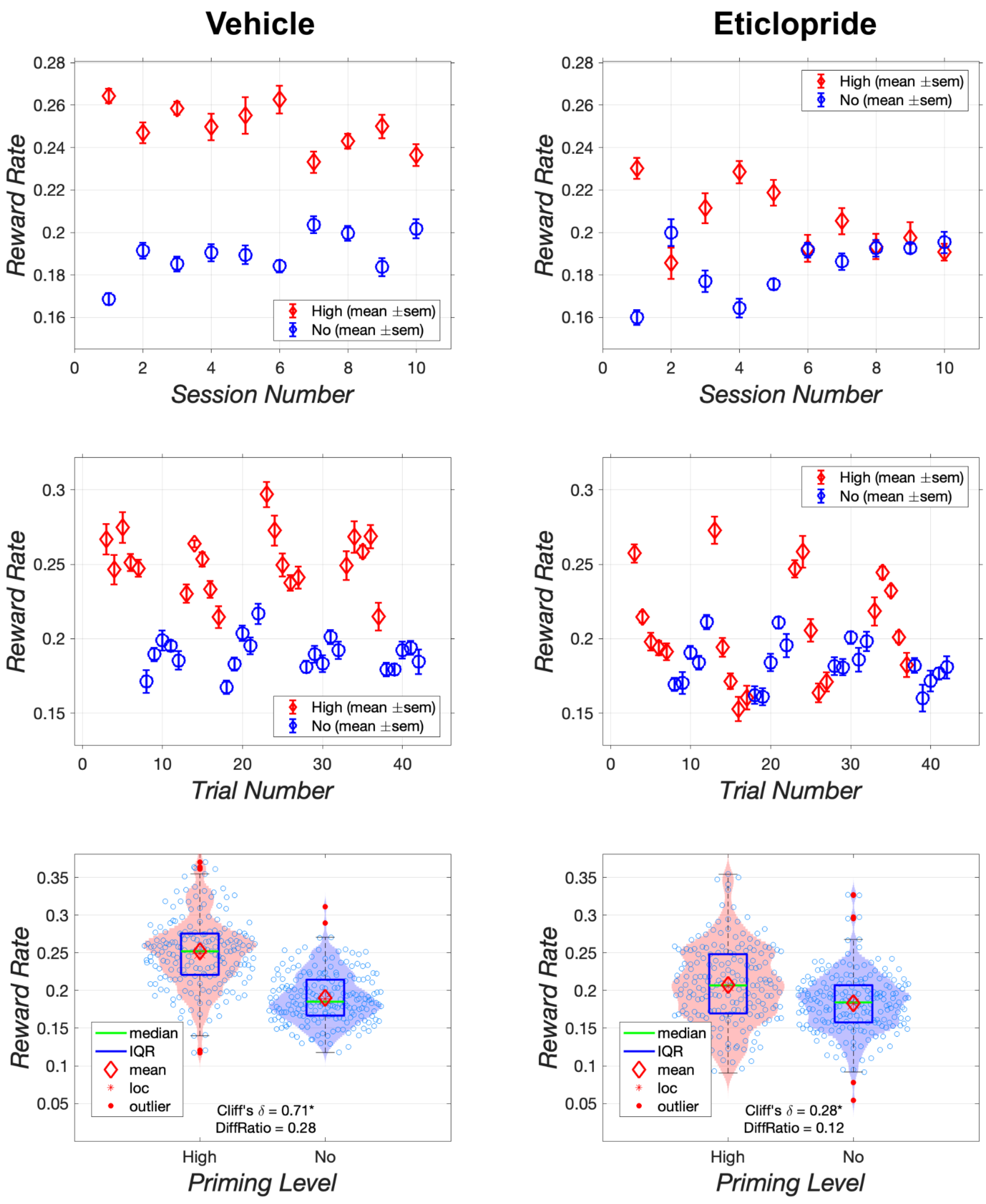


***Figure S17:*** Effect of vehicle and 0.1 mg/kg of eticlopride on reward rate in rat AM55. The upper row shows results collapsed over trials, whereas the middle row shows results collapsed over sessions.

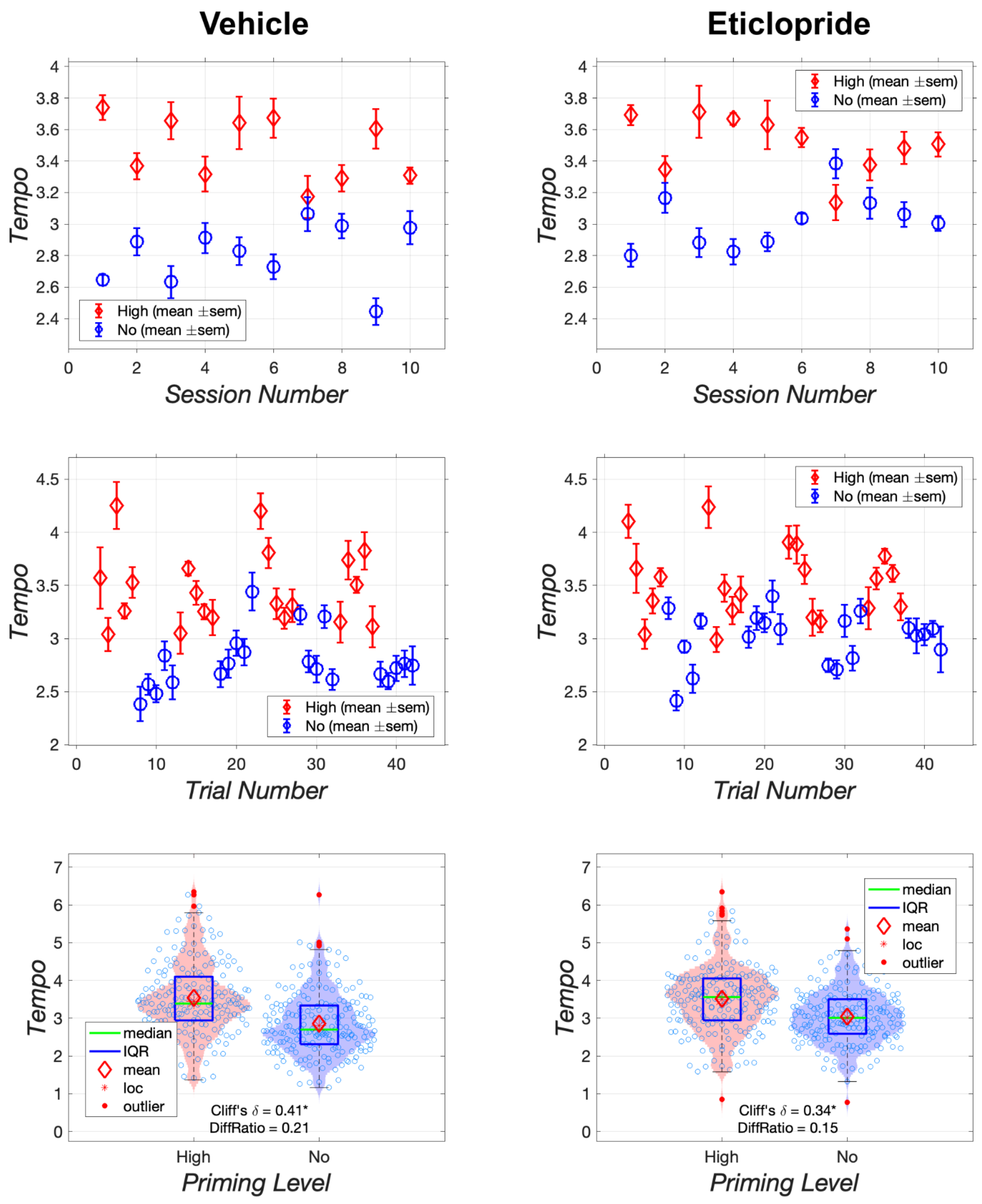


***Figure S18:*** Effect of vehicle and 0.1 mg/kg of eticlopride on the tempo of responding during the setup phase in rat AM55. The upper row shows results collapsed over trials, whereas the middle row shows results collapsed over sessions.

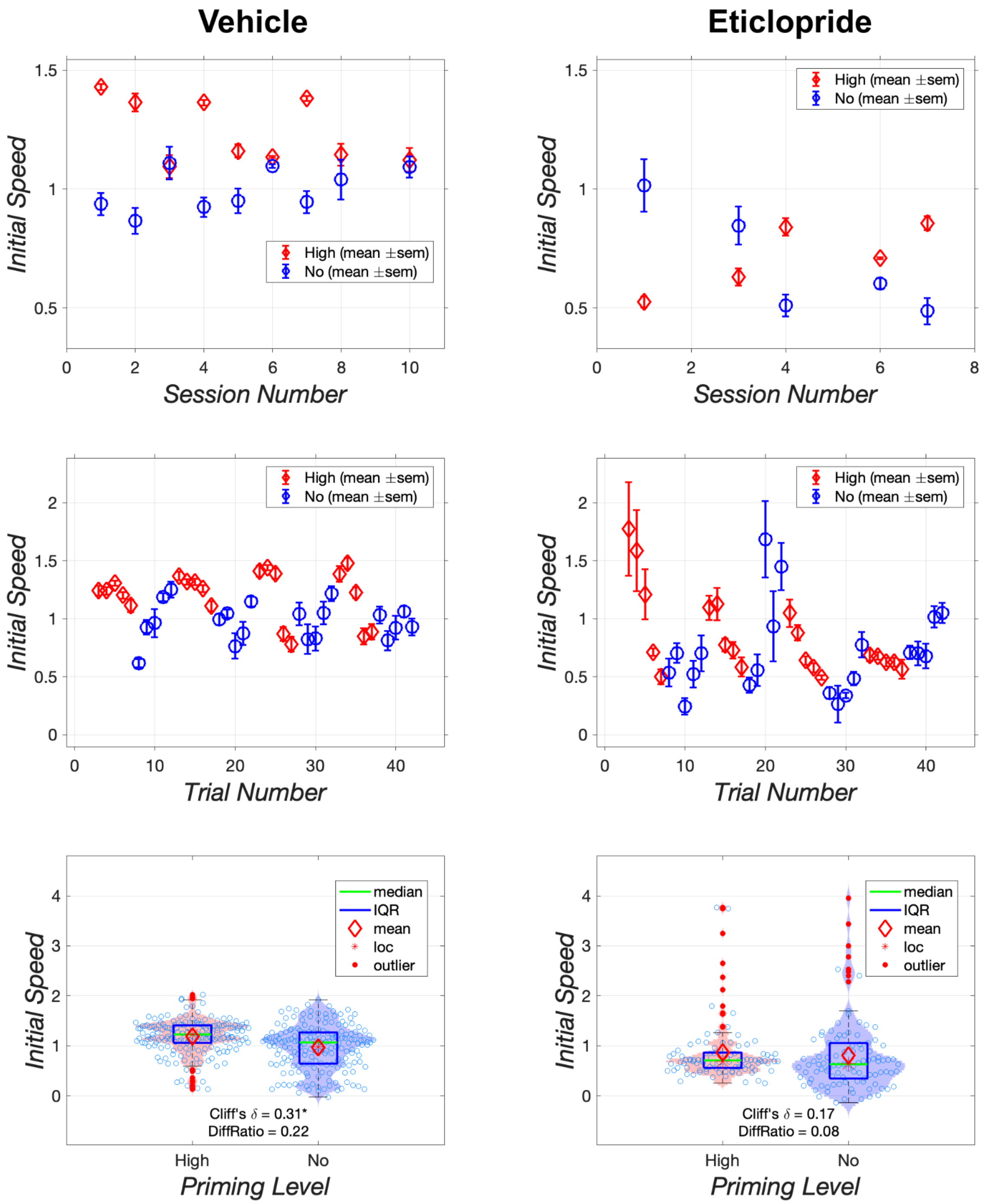


***Figure S19:*** Effect of vehicle and 0.1 mg/kg of eticlopride on the initial speed to initiate responding by rat AM58. The upper row shows results collapsed over trials, whereas the middle row shows results collapsed over sessions.

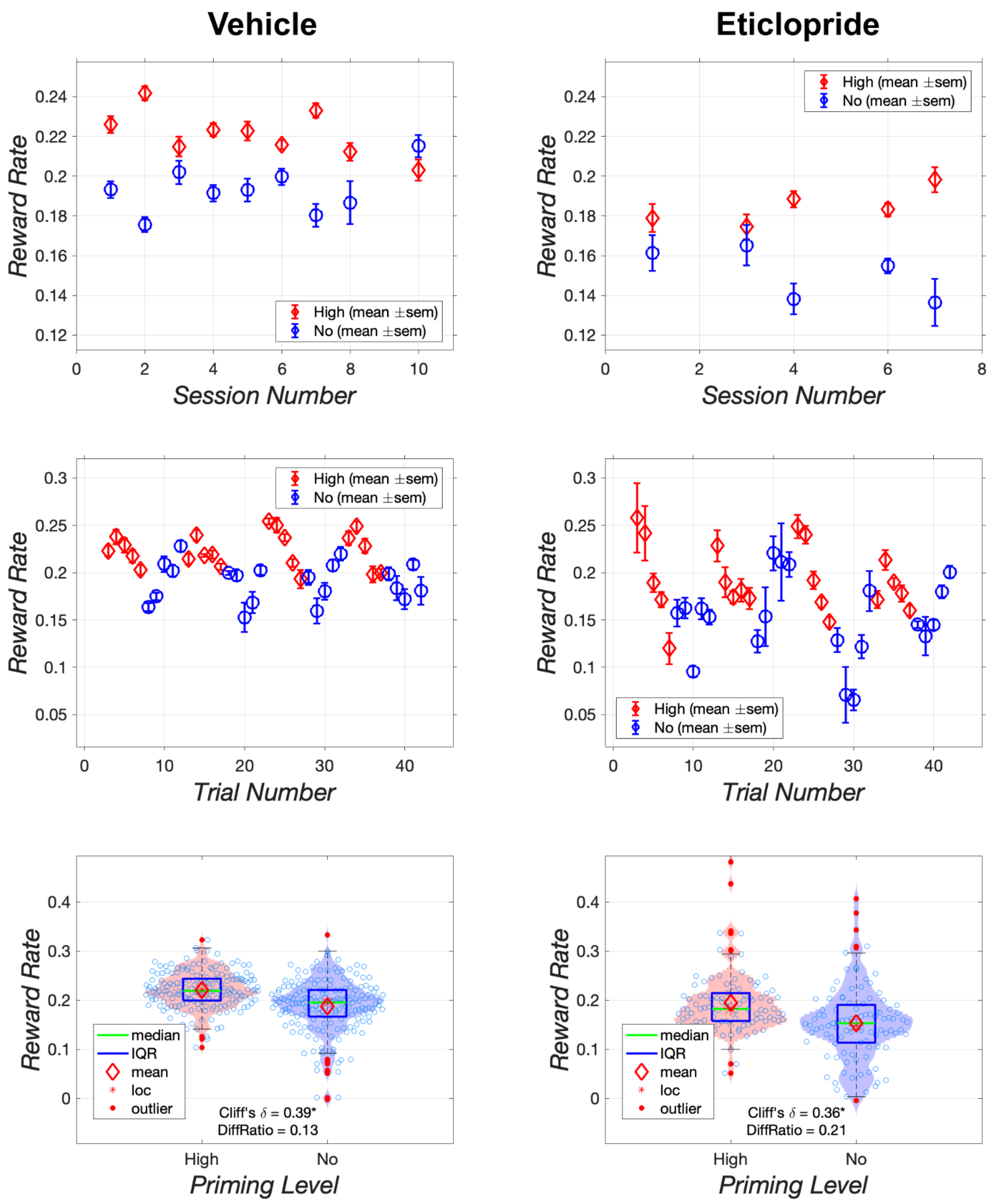


***Figure S20:*** Effect of vehicle and 0.1 mg/kg of eticlopride on reward rate in rat AM58. The upper row shows results collapsed over trials, whereas the middle row shows results collapsed over sessions.

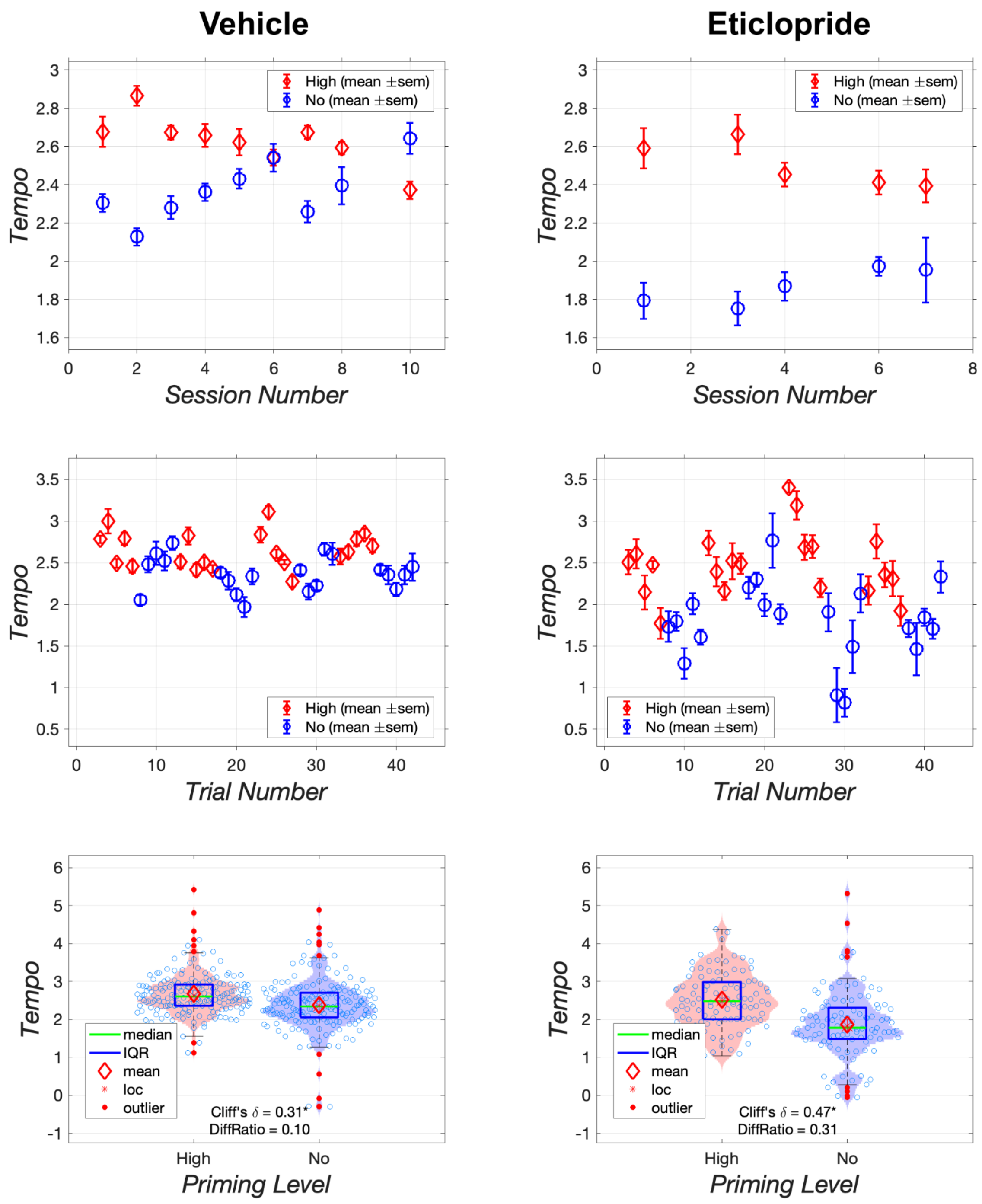


***Figure S21:*** Effect of vehicle and 0.1 mg/kg of eticlopride on the tempo of responding during the setup phase in rat AM58. The upper row shows results collapsed over trials, whereas the middle row shows results collapsed over sessions.

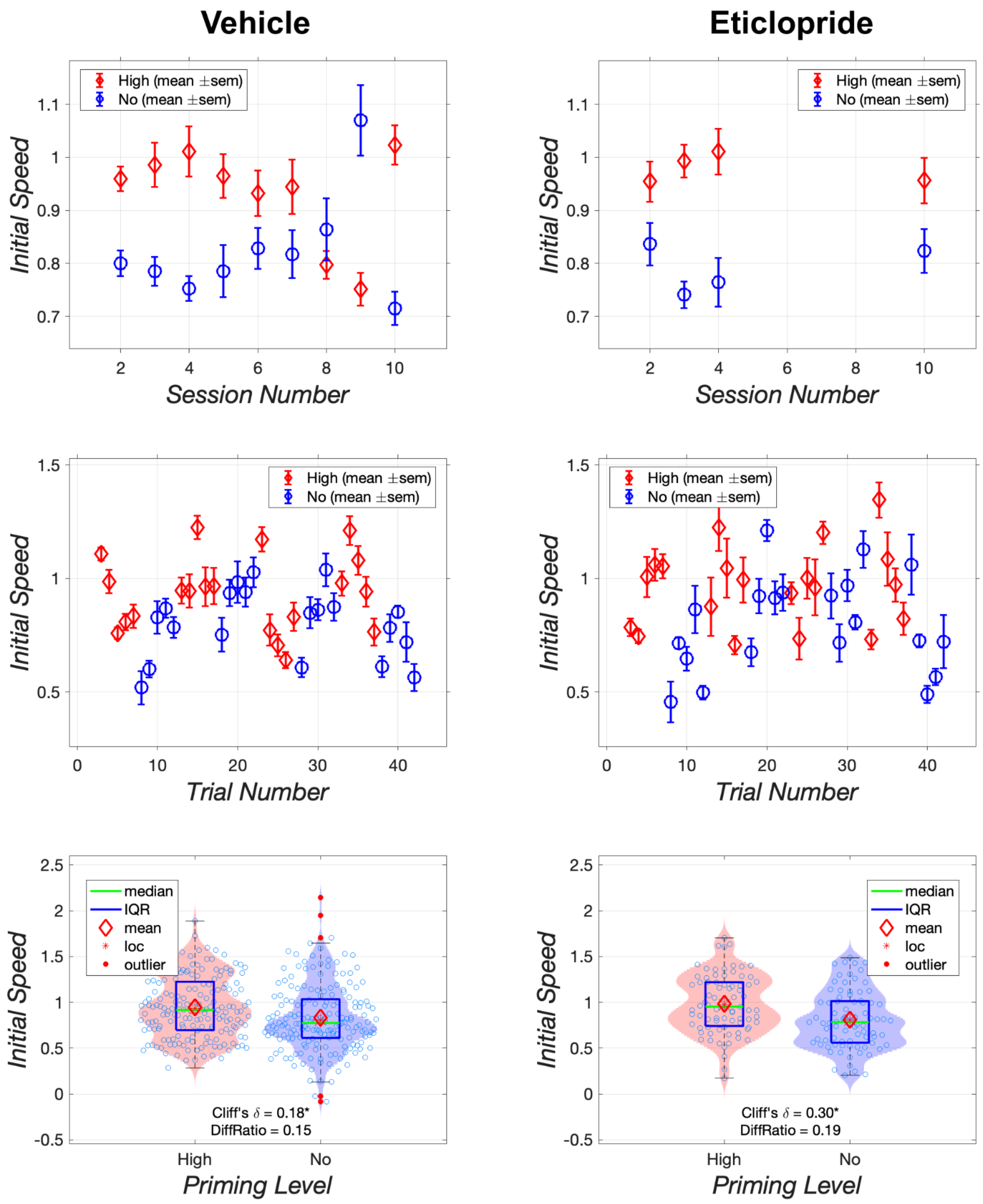


***Figure S22:*** Effect of vehicle and 0.05 mg/kg of eticlopride on the initial speed to initiate responding by rat AM48. The upper row shows results collapsed over trials, whereas the middle row shows results collapsed over sessions.

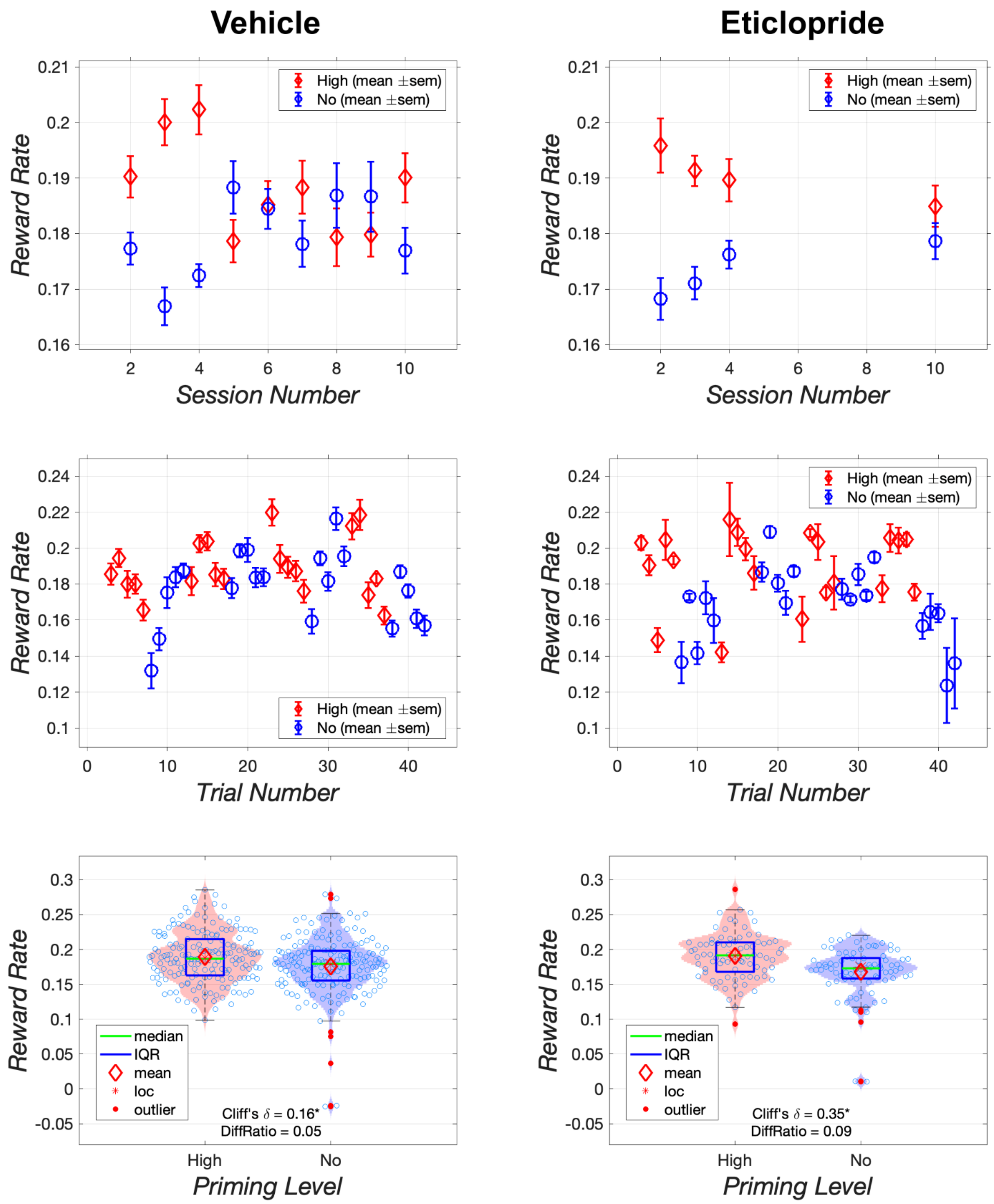


***Figure S23:*** Effect of vehicle and 0.05 mg/kg of eticlopride on reward rate in rat AM48. The upper row shows results collapsed over trials, whereas the middle row shows results collapsed over sessions.

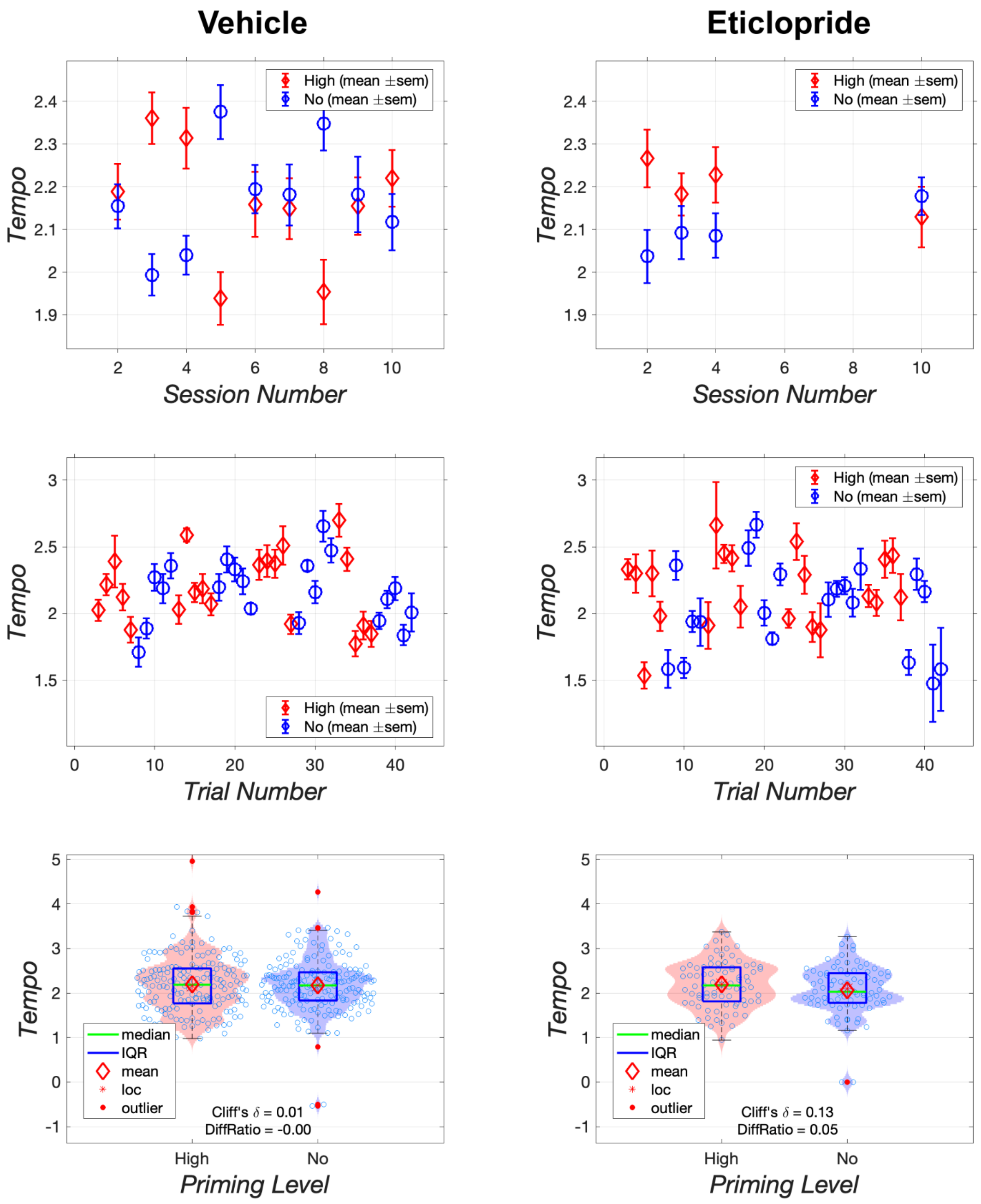


***Figure S24:*** Effect of vehicle and 0.05 mg/kg of eticlopride on the tempo of responding during the setup phase in rat AM48. The upper row shows results collapsed over trials, whereas the middle row shows results collapsed over sessions.

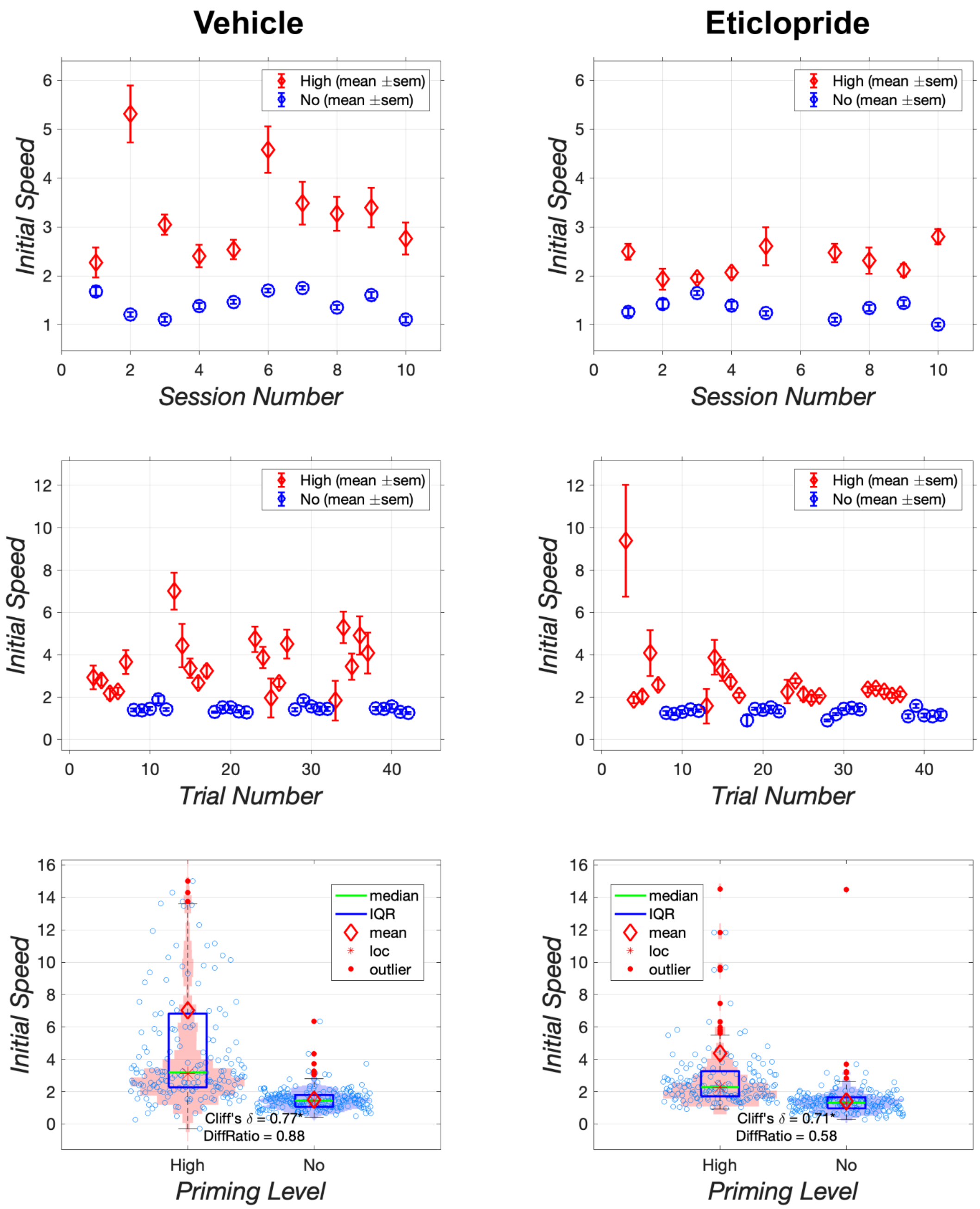


***Figure S25:*** Effect of vehicle and 0.05 mg/kg of eticlopride on the initial speed to initiate responding by rat AM50. The upper row shows results collapsed over trials, whereas the middle row shows results collapsed over sessions.

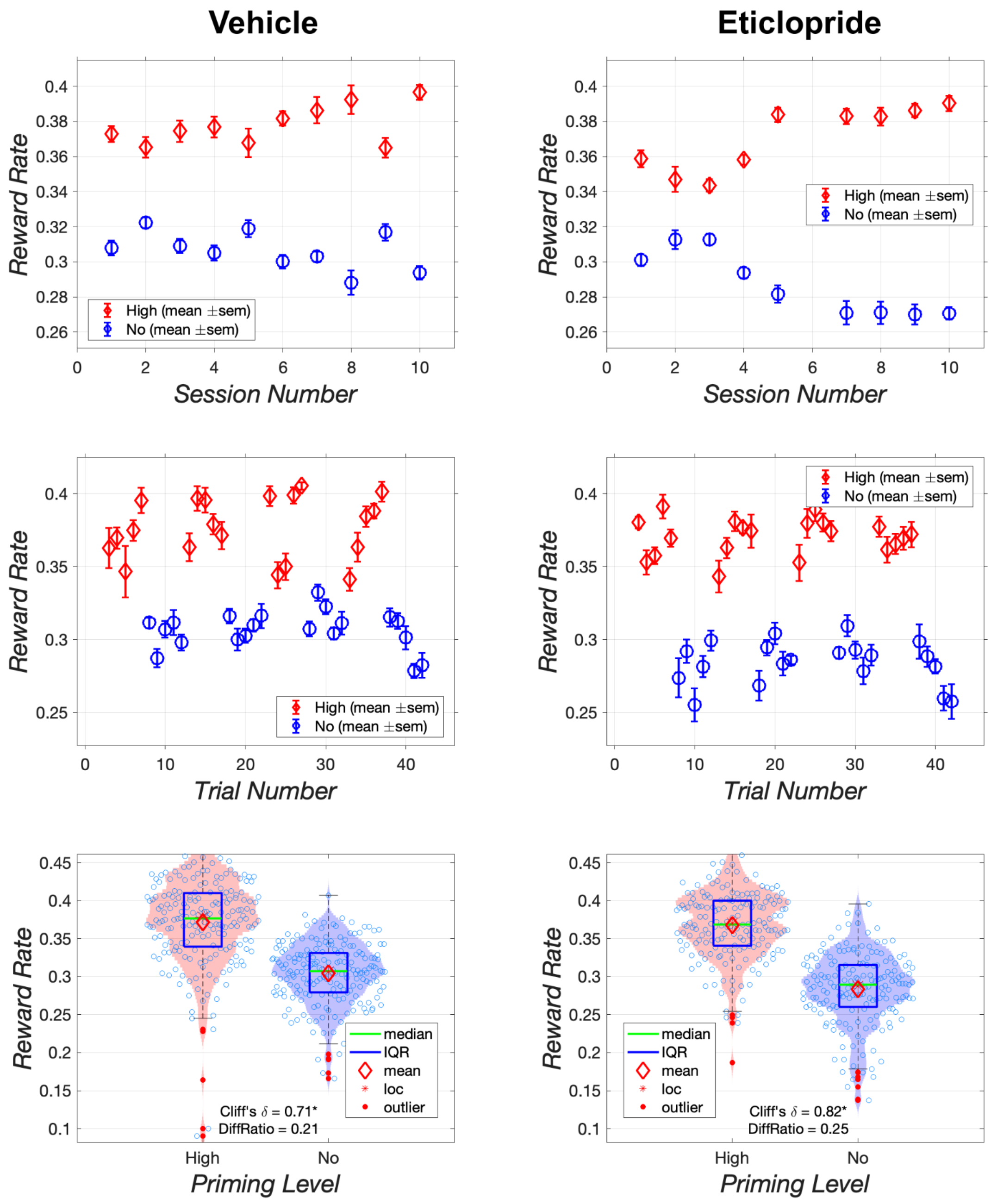


***Figure S26:*** Effect of vehicle and 0.05 mg/kg of eticlopride on reward rate in rat AM50. The upper row shows results collapsed over trials, whereas the middle row shows results collapsed over sessions.

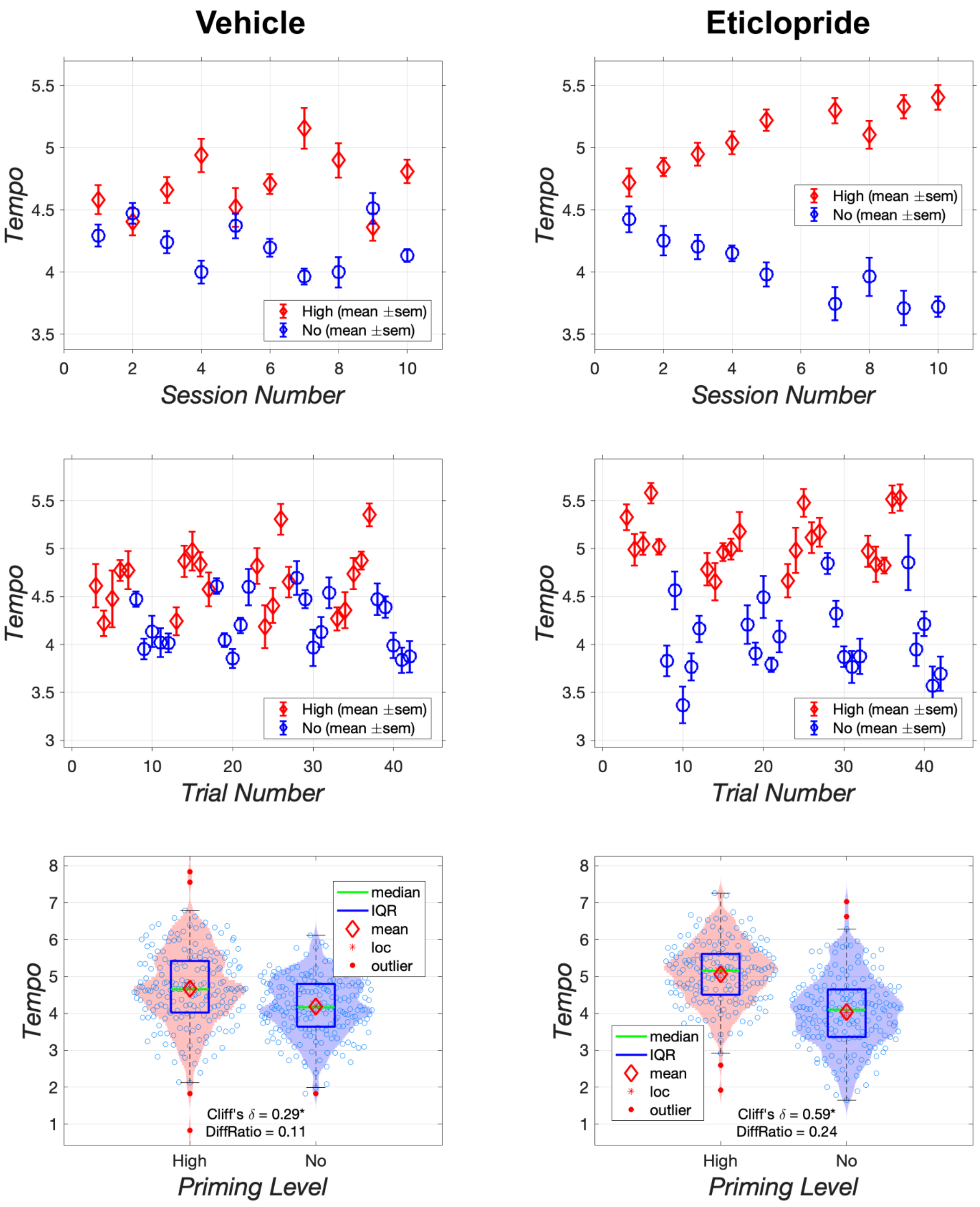


***Figure S27:*** Effect of vehicle and 0.05 mg/kg of eticlopride on the tempo of responding during the setup phase in rat AM50. The upper row shows results collapsed over trials, whereas the middle row shows results collapsed over sessions.
